# Optical Control of G-Actin with a Photoswitchable Latrunculin

**DOI:** 10.1101/2023.07.17.549222

**Authors:** Nynke A. Vepřek, Madeline H. Cooper, Laura Laprell, Emily Jie-Ning Yang, Sander Folkerts, Ruiyang Bao, Thomas G. Oertner, Liza A. Pon, J. Bradley Zuchero, Dirk H. Trauner

## Abstract

Actin is one of the most abundant proteins in eukaryotic cells and a key component of the cytoskeleton. A range of small molecules have emerged that interfere with actin dynamics by either binding to polymeric F-actin or monomeric G-actin to stabilize or destabilize filaments or prevent their formation and growth, respectively. Amongst these, the latrunculins, which bind to G-actin and affect polymerization, are widely used as tools to investigate actin-dependent cellular processes. Here, we report a photoswitchable version of latrunculin, termed opto-latrunculin (**OptoLat**), which binds to G-actin in a light-dependent fashion and affords optical control over actin polymerization. **OptoLat** can be activated with 390 – 490 nm pulsed light and rapidly relaxes to the inactive form in the dark. Light activated **OptoLat** induced depolymerization of F-actin networks in oligodendrocytes and budding yeast, as shown by fluorescence microscopy. Subcellular control of actin dynamics in human cancer cell lines was demonstrated by live cell imaging. Light-activated **OptoLat** also reduced microglia surveillance in organotypic mouse brain slices while ramification was not affected. Incubation in the dark did not alter the structural and functional integrity of microglia. Together, our data demonstrate that **OptoLat** is a useful tool for the elucidation of G-actin dependent dynamic processes in cells and tissues.

## INTRODUCTION

Actin is one of the most abundant proteins in eukaryotic cells and essential to their integrity and dynamics. The actin cytoskeleton is maintained through a dynamic interplay between globular, monomeric G-actin and filamentous, polymeric F-actin (Fig. 1a). This homeostasis is regulated by numerous actin binding proteins that are involved in a wide variety of signaling cascades.^1–3^ Actin dynamics can also be altered by small molecules, which can have stabilizing or destabilizing effects on Factin.^4^ Stabilizers, such as phalloidin and jasplakinolide, bind to the filaments and prevent depolymerization.^5–7^ The cytochalasins bind to the terminal monomers on a filament (barbed end), preventing further polymerization. Amongst the actin destabilizing natural products, latrunculins A and B stand out by their unique binding site and mechanism of action. Latrunculins A and B bind G-actin in the nucleotide binding cleft between subdomains 2 and 4 (Fig. 1b/c).^8^ The binding of latrunculin in this site prevents conformational changes that are crucial for the incorporation of G-actin into the polymerizing filament.^8^ The main mechanism of action is therefore the sequestration of actin monomers. Additionally, latrunculin A can be involved in the active severing of filaments.^9^

**Figure 1:**
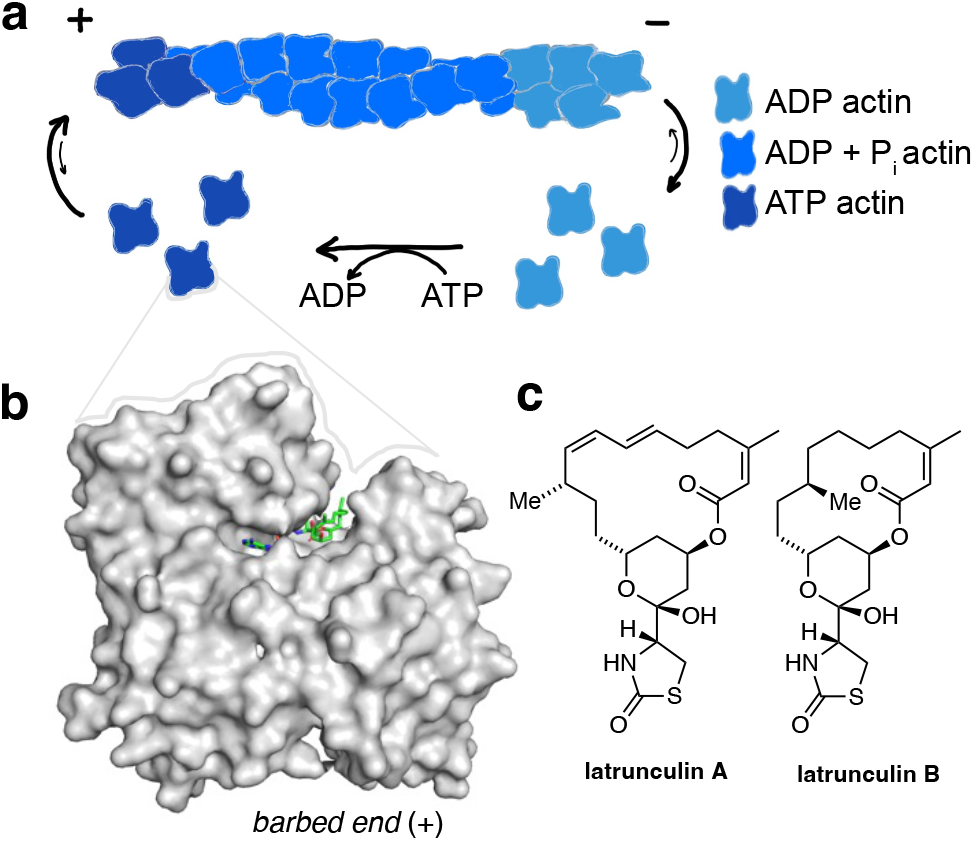
Actin turnover is inhibited by Latrunculin binding to Gactin. a) Schematic representation of actin turnover (treadmilling) b) Latrunculins bind G-actin above the ATP binding cleft (pdb: 2q0u). c) Chemical structures of latrunculins A and B.

The study of actin dynamics in cells and tissues is challenging because of the many factors control the remodeling of the dynamic actin networks. To address some of these challenges, we have previously introduced photoswitchable versions of the actin stabilizer jasplakinolide that allow for the reversible stabilization of actin filaments with sub-cellular precision.^10,11^

Although jasplakinolide-based probes have been used to the study the effect of F-actin stabilization on cytoskeletal function, actin dynamics in cells also depend on the highly regulated Gactin pool.^12^ G-actin dependent remodeling of actin, for instance during cell migration, takes place within seconds-minutes.^13^ The investigation of such processes requires tools that allow for the modulation of the G-actin pool at the time scales of the biological processes under investigation. Here, we describe optical probes that are complementary to the F-actin stabilizers previously introduced and derived from the G-actin binders latrunculin A and B. Our photoswitchable version of latrunculin, **OptoLat**, can sequester G-actin monomers in a light dependent, reversible fashion leading to spatially and temporally confined breakdown of actin networks. We show that actin dynamics can not only be altered in cultured human cancer cell lines with **OptoLat**, but also in mammalian oligodendrocytes and microglia, as well as the budding yeast *Saccharomyces cerevisiae*.

## RESULTS

### Design and Synthesis

We based the design of opto-latrunculins on published x-ray crystal structures and known structureactivity relationship studies.^14–18^ Latrunculins bind G-actin right next to the nucleoside binding cleft between subdomains 2 and 4 and inhibit the conformational twist that is essential for incorporation of ATP-bound G-actin into F-actin (Fig 2a).^8^ Their thioazolidinone core and hemiacetal engages in hydrogen bonding with the protein whereas the lipophilic macrocycle is predominantly involved in hydrophobic interactions (Fig. 2a).^17–19^

**Figure 2:**
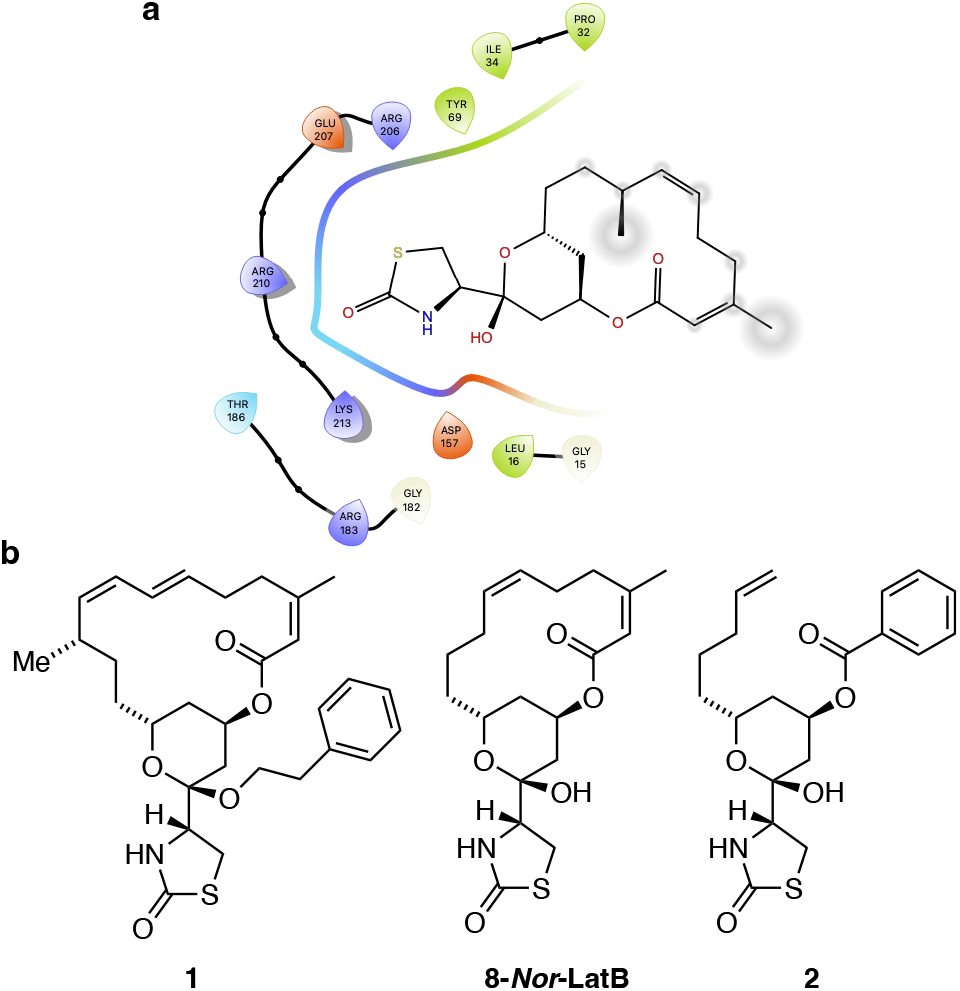
**a)** Interactions of latrunculin B with the G-actin. b) Chemical structures of modified or truncated latrunculins identified in SAR studies.

**Figure 2:**
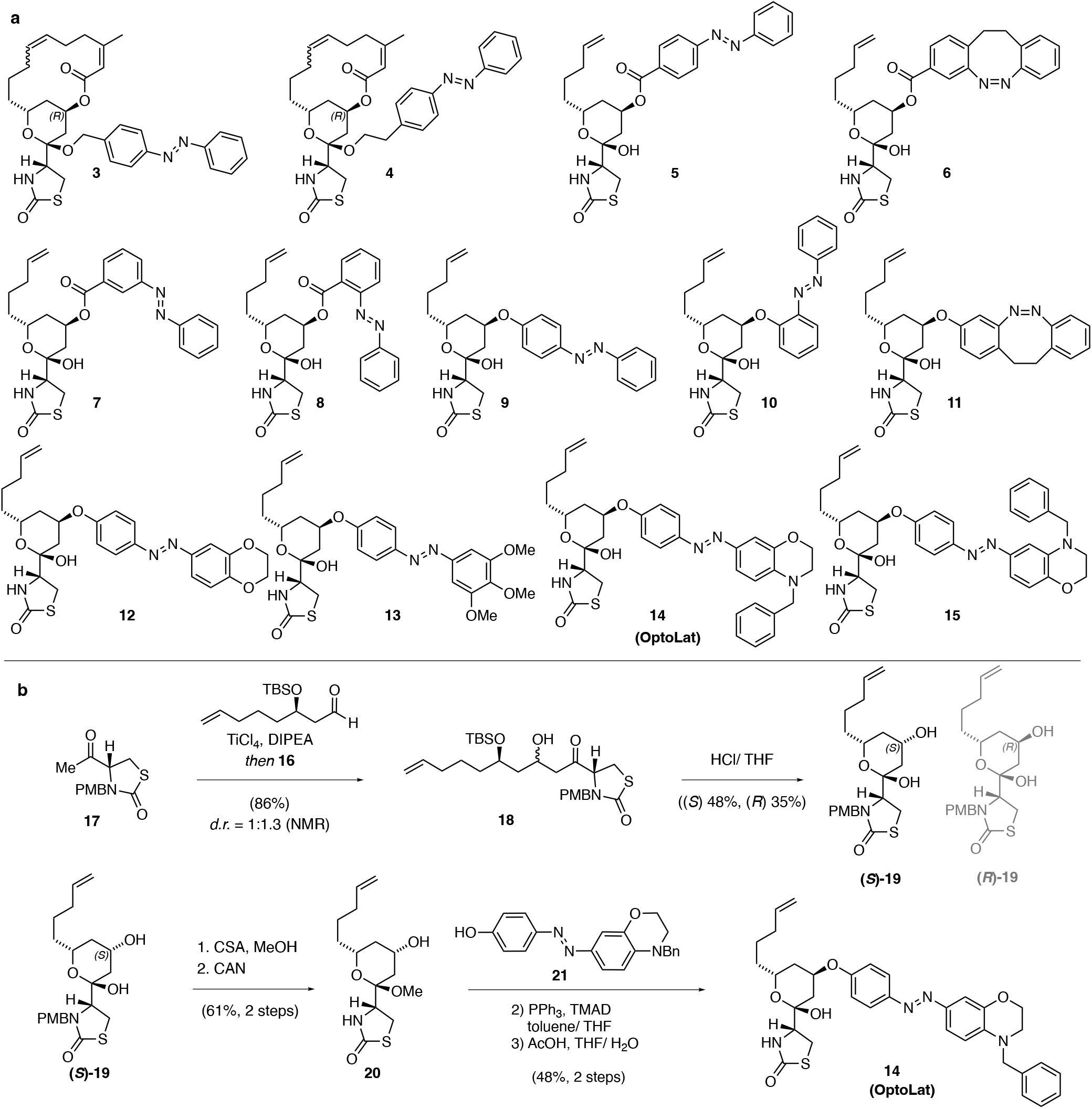
Photoswitchable latrunculin derivatives and synthesis of OptoLat. a) Photoswitchable latrunculin derivatives synthesized. b) and c) Synthetic route towards **OptoLat**.

Structure-activity relationship studies have identified certain modifications of latrunculins that retain bioactivity. For instance, acetal **1**, derived from latrunculin itself was shown to be a 1.0 μM inhibitor in biochemical studies.^16^ The fully synthetic derivative, **8-*Nor*-latrunculin B**, was found to cause actin disruption at μM concentrations.^20^ *Seco* derivatives such as the benzoate **2** were also found to be active, albeit with slightly reduced potency.^20^

Based on this knowledge, we designed the photoswitchable derivatives shown in Fig. 3a. These include photoswitchable acetals such as **3** and **4**, which retain the macrocycle and were based on **1**, as well as azobenzene and diazocine esters such as **5** and **6**, inspired by **2**. To increase the steric clash between the molecule and protein by photoisomerization of the diazine bond, we also installed *meta* and *ortho* derivatives **7** and **8**. Concerned about the stability of esters, we explored derivatives bearing aryl ethers that had not been previously described. In addition to metabolic stability, aryl ethers would also bring the photoswitch deeper into the protein pocket which could increase steric effects. To this end, we synthesized azobenzenes **9** and **10** and diazocine **11**. In the next iteration, we aimed at increasing the bulk of the photoswitch by introducing different functionalities such as oxirane, trimethoxy and *N*-benzyl aryls on the photoswitch, generating aryl ethers **12** – **15**. Amongst these **14**, bearing a *para N*-benzyl azobenzene photoswitch, emerged as the most useful compound and was termed **OptoLat**.

The synthesis of **OptoLat** (**14**) (Fig. 3b) started with a titanium tetrachloride mediated aldol addition of enantiomerically pure aldehyde **16** to PMB-protected thioazolidinone **17**. As described by Fürstner, acid mediated silyl deprotection resulted in cyclization and gave a separable (1.25:1) mixture of hemiacetals (*S*) and (*R*)-**19**.^21^ (*S*)-**19** was converted into the methyl acetal following oxidative PMB deprotection.^21,22^ Subsequent Mitsunobu reaction with azobenzene **21** under optimized conditions using TMAD, followed by hydrolysis, gave **OptoLat** in good overall yield.

Photoswitchable acetals **3** and **4** were procured from the corresponding hemiacetals as described in the Supporting Information. Aryl ethers **5** – **6** were made by Mitsunobu reactions from (*S*)-**19**. Ether derivatives **9, 11** – **13** and **15** were obtained by similar synthetic strategies as **OptoLat. 10** was accessed by S_N_Ar from (*R*)-**19**.

### Initial photophysical and biological evaluation

All photoswitchable latrunculins were characterized for their photophysical properties (see Supporting Information). Due to its *para* amino substituent, **OptoLat** showed the most red-shifted absorption spectrum (λ_max_ trans ca. 450 nm) and fastest thermal relaxation (t_1/2_ < 2 s in DMSO).^23,24^ *trans* **OptoLat** can be efficiently activated with wavelengths between 390 and 490 nm (Fig. 4e). The activation efficiency was determined by NMR with *in situ* irradiation using ultra high-power LEDs (see SI). The highest ratio of *cis* isomer was reached with 460 nm (ca. 67%). Upon irradiation with 390 nm ca. 46% *cis* and 520 nm ca. 26% *cis* were obtained (Fig. 4f). The *cis* isomer was shortlived (t_1/2_ ca. 1.6 s, Fig. 4g) which resulted in complete deactivation of the photoswitch in the absence of light within seconds. Switching was fully reversible and did not show fatigue (Fig. 4h).

**Figure 4:**
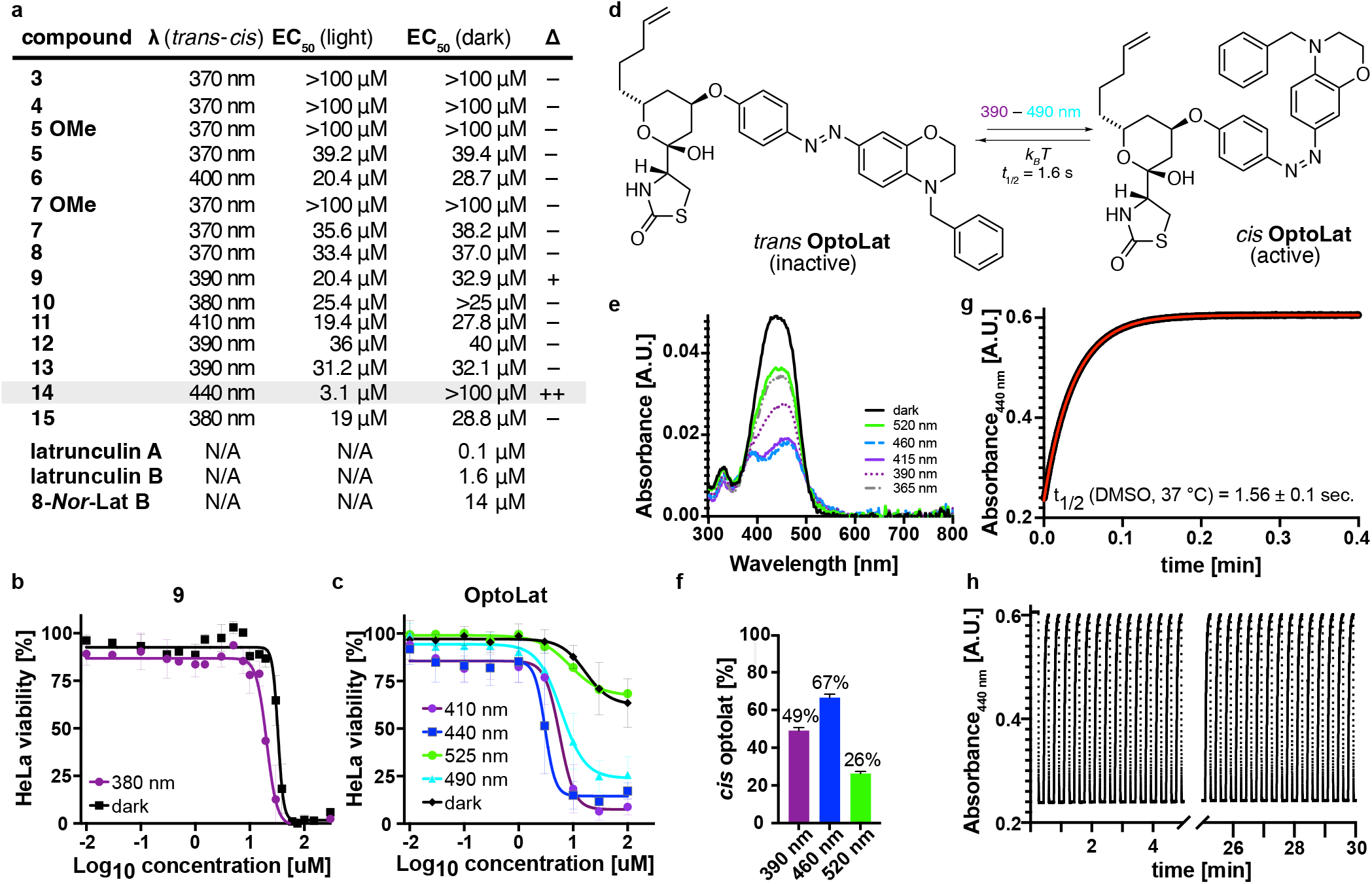
Initial biological evaluation with cell proliferation assays and photophysical properties of OptoLat. a) Antiproliferative activity of photoswitchable latrunculin analogs and reference compounds. b) Cell viability dose-response curves for **9** under dark and illuminated conditions. c) Cell viability dose-response curves for **OptoLat** (**14**) under illumination with different wavelengths and in the dark. d) Photochemical and thermal isomerization of OptoLat. e) UV-Vis absorption spectrum of **OptoLat**. f) Percentage of *cis* **OptoLat** measured by NMR with *in situ* irradiation at different wavelegths. g) Thermal relaxation of of *cis* **OptoLat** in the dark in DMSO solution at 37 °C. h) switching of **OptoLat** is reversible and fatigue-resistant over multiple cycles.

Next, we evaluated our photoswitches using light-dependent cell proliferation (MTT) assays in HeLa cells. Acetals **3** and **4**, derived from phenethyl latrunculin A (**1)**, were found to be completely inactive. *Seco* esters **5 – 8**, proved to be active, albeit with high EC_50_ values compared to macrocyclic reference compounds (e.g. **8-*Nor*-latB**). However, they did not produce any light-dependent antiproliferative activity. Their methyl acetals, which are synthetic intermediates, were completely inactive (**5 OMe** and **7 OMe**). Azoaryl ether **9** showed a small difference in light versus dark activity (Fig. 3b). However, other modifications of the photoswitch including the incorporation of an *ortho* azobenzene (**10**) and a diazocine (**11**), as well as more bulky substituents in **12** and **13** did not lead to a widening in the photopharmacological window. This was also true for “iso”Optolat **15** which has increased steric bulk by virtue of its benzyl substituent. Gratifyingly, its isomer **OptoLat** (**14**), which has desirable photophysical and thermal properties, proved to be virtually inactive in the dark but became highly active upon irradiation with blue light (Fig. 3c/d). Therefore, our detailed biological investigations focused on this photoswitchable form of latrunculins A/B. Cell proliferation assays across different cell lines showed that **OptoLat** has an EC_50_ of 2.5-4.1 μM (see SI). As such, it is about three times less active than **latrunculin B** itself. Compared to **8-*Nor*-lat B**, however, **OptoLat** showed 2.5 times higher antiproliferative activity in our HeLa assays (Fig. 4a/c)

### Shaping actin-based structures in oligodendrocytes and budding yeast cells

Next, we investigated, whether the strong antiproliferative effect indeed resulted from the destabilization of actin networks in different cell types. HeLa cells treated with **OptoLat** and kept in the dark showed normal actin structures, which broke down upon pulsed irradiation with 410 nm within 24 h. The quantification of these results, however, was difficult since breakdown of the actin network also affects the size and shape in HeLa cells. We therefore turned to oligodendrocytes, glial cells that develop extensive actin networks during their maturation but do not require actin to maintain their structural integrity.^25^ This feature makes them an ideal cell type for testing actin perturbing agents like **OptoLat**. We treated rat oligodendrocytes with escalating doses of **OptoLat**. Following 6 hours of **OptoLat** activation (pulsed irradiation with 430 nm), we observed a significant decrease in filamentous actin at all **OptoLat** doses as measured by phalloidin staining (Fig. 5a). The degree of actin disassembly was similar to the degree of disassembly seen in response to treatment with 100 nM Latrunculin A (Fig. 5b). We observed no difference between levels of filamentous actin in oligodendrocytes treated with **OptoLat** that were kept in the dark, or DMSO, demonstrating the lack of basal activity of *trans* **OptoLat**.

**Figure 5:**
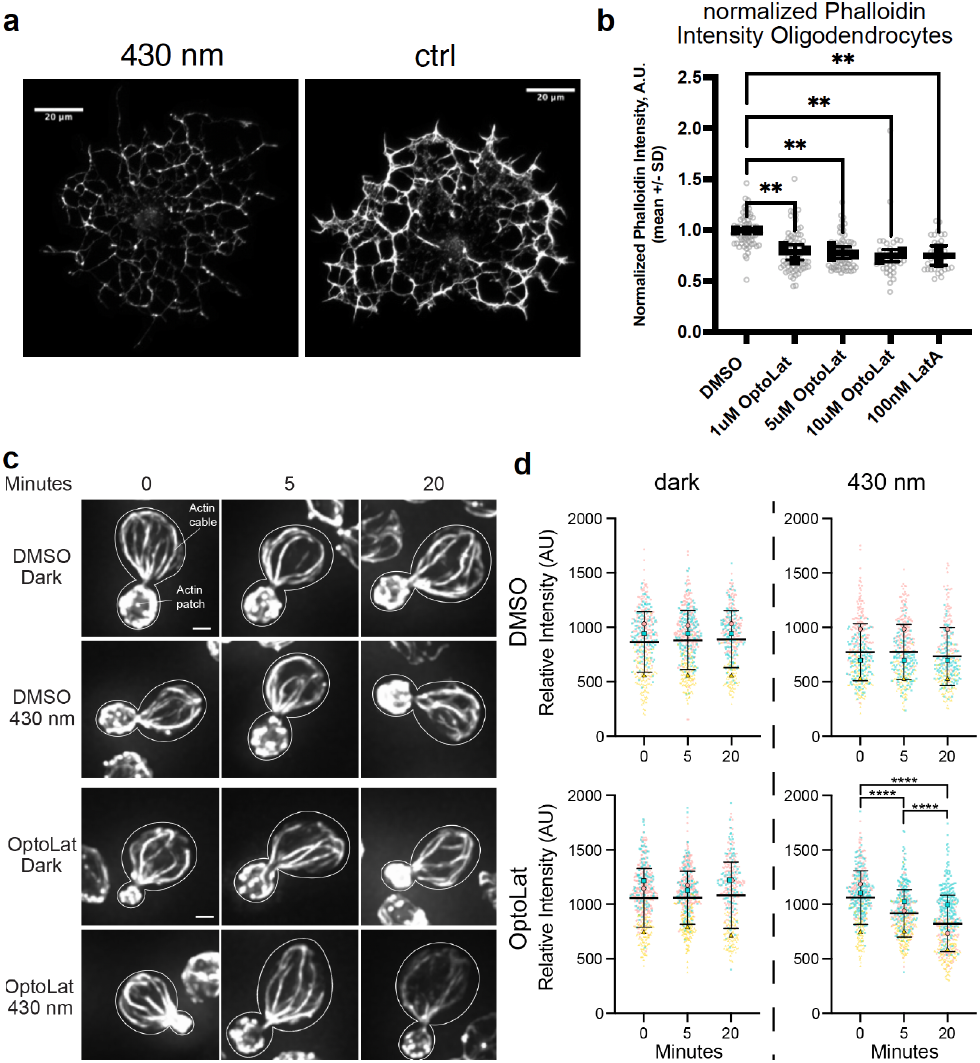
Light dependent reduction of actin content using **OptoLat** a) Depletion of F-actin in oligodendrocytes visualized using phalloidin staining after irradiating with 430 nm light b) Quantification of F-actin depletion in oligodendrocytes. ** *p* ≤ 0.01, values were calculated using one-way ANOVA followed by Dunnett’s multiple comparison test. c) Mid-log phase yeast cells were incubated in the presence of absence of 100 μM **OptoLat** in the dark or upon exposure to pulsed blue light (430 nm) for 0 – 20 min. Cells were then fixed and stained for F-actin using AlexaFluor488 (AF488)-phalloidin. c) Maximum projections of the actin cytoskeleton in the presence and absence of **OptoLat** activation. Cell outlines are shown in white. Scale bar: 1 μm. d) Quantitation of AF488-phalloidin-stained F-actin in actin cables. Results are given as the mean ± standard deviation (SD) from 3 independent trials. n > 80 cells/time point/condition/trial. The mean (large symbols) and all datapoints (small symbols) of 3 independent trials are shown, with differently shaped and colored symbols for each trial. Statistical significance was determined by the two-way ANOVA, with Tukey’s multiple comparisons test. **** *p* ≤ 0.0001.

Since actin is highly conserved across species, we wondered whether **OptoLat** is also active in yeast cells. In budding yeast (*S. cerevisiae*), there are three major actin-containing structures: actin patches, contractile rings, and actin cables. Actin patches are endosomes bearing an F-actin coat that localize to the bud (daughter cell)^26^. Contractile rings drive separation of mother and daughter cells during cell division, in yeast as in other eukaryotes, and localize to bud neck. Actin cables are actin bundles that assemble in the bud and extend along the mother-bud axis. They are essential for cell proliferation for their function as tracks for transport of organelles and other cellular constituents from mother to daughter cells during cell division^27,28^. Finally, actin cables are dynamic structures that undergo rapid turnover and treadmilling from their site of assembly in the bud toward the mother cell^29^. As a result, they highly sensitive destabilization by latrunculin^30^.

One of the advantages of a fast-relaxing photoswitchable actin destabilizer is that it could allow for the modulation of actin dynamics with temporal precision. Therefore, we monitored the effect of **OptoLat** on yeast actin cables by measuring the fluorescence intensity of F-actin in fluorochrome-coupled phalloidin-stained actin cables. Treatment of budding yeast cells with **OptoLat** in the dark or with DMSO in the dark or with illumination at 430 nm had no detectable effect on actin cables. By contrast, activation of **OptoLat** with pulsed blue light (430 nm) resulted in a rapid, time-dependent decrease in the Factin content in actin cables (Fig. 5c,d). Indeed, we observe a significant decrease in actin cables within 5 min of **OptoLat** activation and a further decrease in those structures after 20 min of **OptoLat** activation.

### Wound healing Assays

Since the remodeling of actin networks is essential to cell migration,^31–34^ we next investigated whether **OptoLat** reduces the migration of invasive cancer cells in a light dependent fashion, without showing significant toxicity. In a wound healing assay, **OptoLat** did not affect wound closure in the dark (Fig. 6). Under pulsed-irradiated conditions however, **OptoLat** significantly reduced wound closure. *cis* **OptoLat** was not cytotoxic at concentrations that strongly impaired cell migration (9 μM, 24 h, 440 nm) (Fig. 6c).

**Figure 6:**
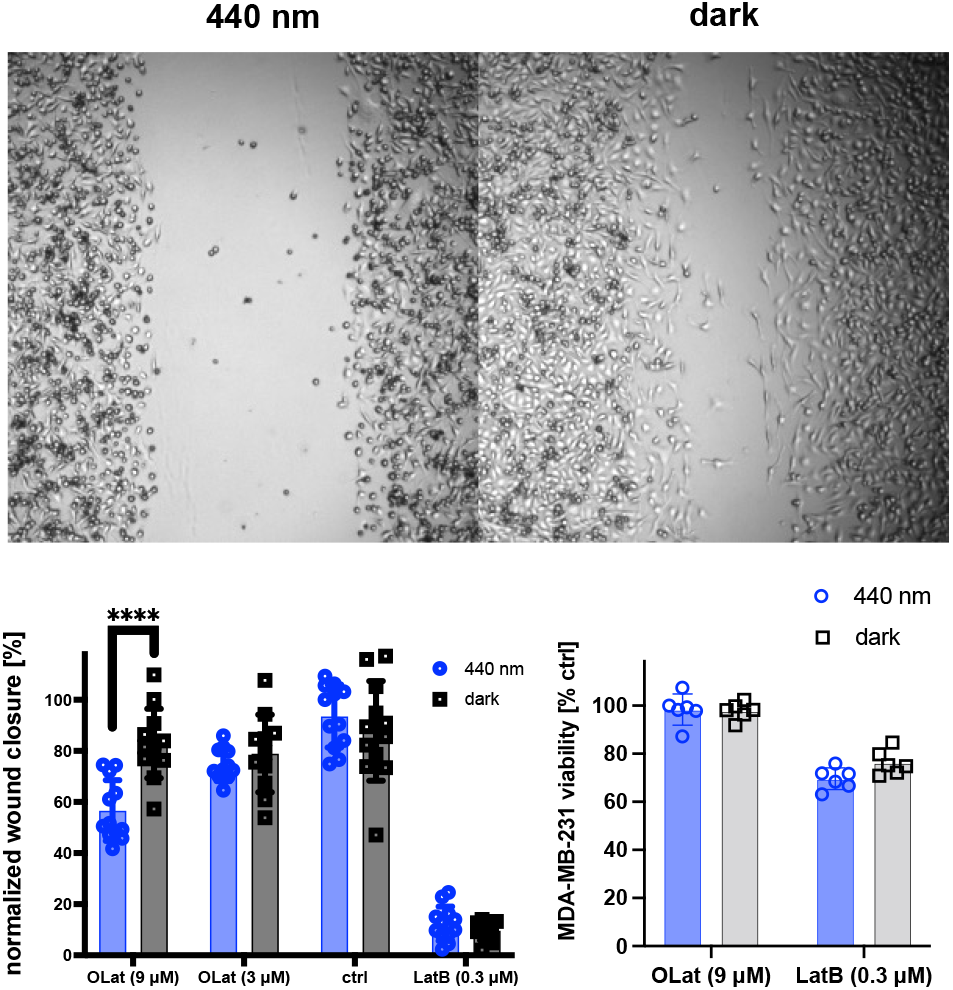
Light-dependent wound closure of MDA-MB-231 cells. a) Representative images of wound closure under irradiation with 440 nm and in the dark. b) Quantification of wound closure after 24 h. c) Cell viability after 48 h under treatment conditions. **** *p* ≤ 0.0001. Statistical analysis was performed using Mann-Whitney test.

### Reversible perturbation of actin dynamics in networks in live cells

The actin networks in lamellipodia show very high turnover rates as these are required to drive cell migration.^31–33^ To further assess the temporal and spatial precision of **OptoLat**, we investigated the ability of **OptoLat** to modulate actin dynamics in the lamellipodia of migrating cells with subcellular resolution. MDA-MB-231 cells that stably express mCherry LifeAct, a fluorescent protein that labels F-actin,^35^ were incubated with **OptoLat** overnight and kept in the dark. Under these conditions, the cells were directly compared to vehicle controls and behaved normally. The local pulsed irradiation of a region of interest (ROI) in the lamellipodium lead to a reduction of protrusion dynamics, which was followed by a breakdown of the actin network. Upon keeping the cells in the dark, actin networks fully recovered (Fig. 7, Supplementary Movie). This recovery was quantified through normalization of LifeAct fluorescence in the ROI (Fig. 7b). F-actin fluorescence was steeply reduced upon **OptoLat** activation. The destructed network persisted for several minutes, before a full recovery was observed. Irradiation with 440 nm light alone did not lead to the destruction of the actin network in control cells (Fig 7, DMSO control). The dynamic changes of actin turnover upon light irradiation can also be clearly visualized using kymographs (Fig. 7c).

**Figure 7:**
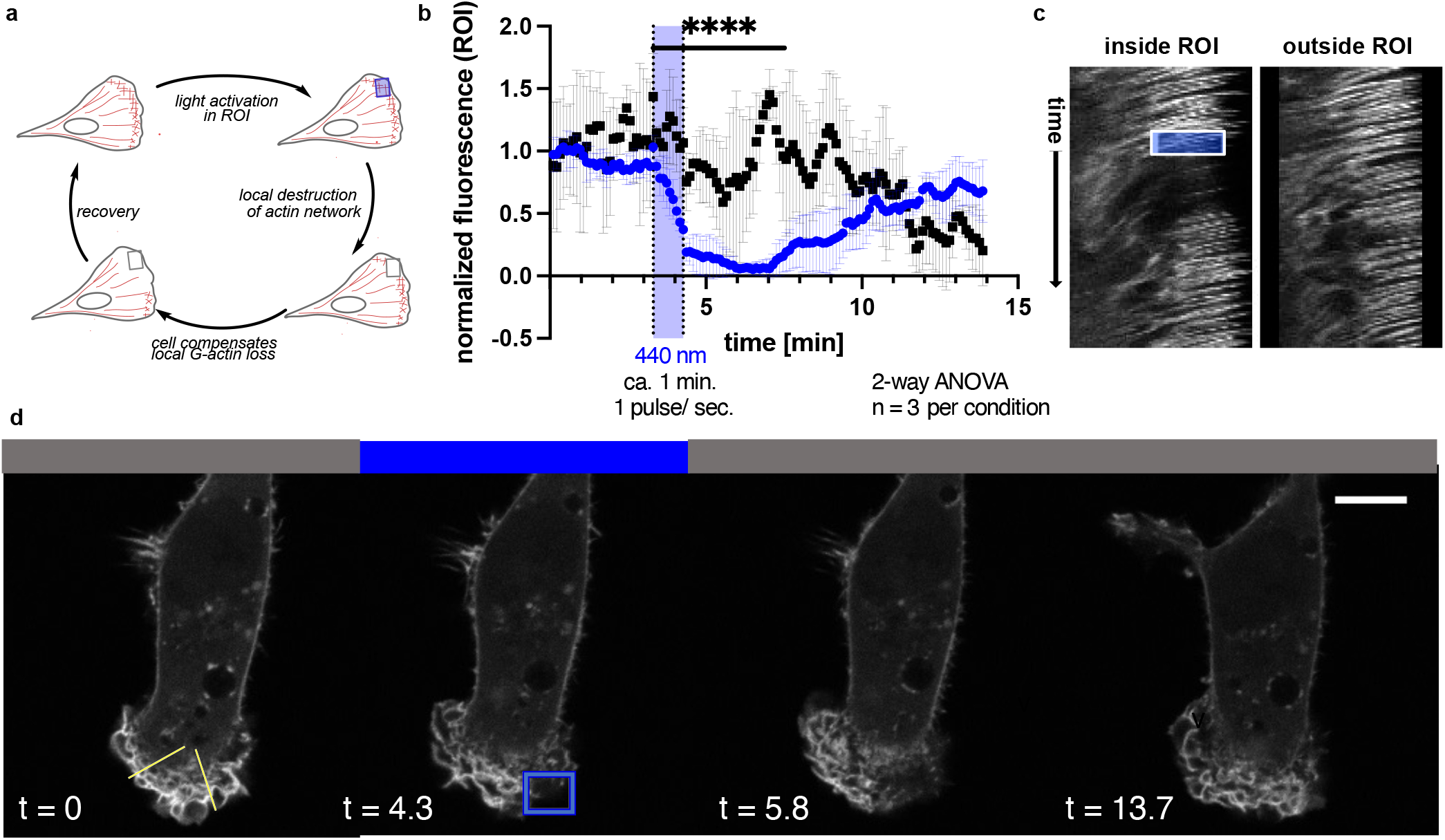
Cell Migration and local subcellular remodeling of actin networks. a) Schematic of local **OptoLat** activation, actin network destruction and recovery. ROI = region of irradiation. b) Quantification of fluorescence inside and outside the ROI for n=3 per condition. Black squares = region outside of the ROI; blue circles = within the ROI. c) Kymographs inside and outside of the ROI along the yellow lines in d). The box represents the duration and extent of irradiation in the ROI. d) Time course images of a representative cell treated with OptoLat. Yellow lines in the left panel indicate selections for the kymographs (left: outside of the ROI; right: within the ROI).

### Effects on microglia in brain slices

Having shown that **OptoLat** was effective in cultured cells, we tested whether our novel tool could also be applied in complex tissues. As an example, we investigated microglial surveillance activity in organotypic brain slices in the presence of **OptoLat** under dark and light-activated conditions. Microglia are highly branched immune cells that constantly scan the brain parenchyma by extending and retracting filopodia-like structures (surveillance). These processes are highly dynamic and depend on actin remodeling, while the overall shape of the cell (ramification) depends on microtubule networks.^36^

To investigate whether **OptoLat** affects the behavior of microglial cells, we prepared hippocampal slice cultures from mice expressing a red fluorescent protein exclusively in microglia (Cx3Cr1Cre-tdT). We incubated slices cultures with **OptoLat** overnight and imaged microglia 60-80 um below the surface of the slice culture surface with two-photon microscopy (Fig. 8a). From maximum intensity projections taken at two time points (Δt = 2 min) we quantified the cell’s shape (ramification) and its motility (surveillance). The ramification index is a measure of the ratio of the cell’s perimeter to its area (Fig. 8b), normalized to that of a circle of the same area. The surveillance index is a measure of the area (pixels) surveyed per unit time (Fig. 8c). In the dark, ramification and surveillance of microglia conditioned with **OptoLat** was not different from vehicle controls. **OptoLat** activation by pulsed blue light did not affect ramification (Fig. 8d), but induced a significant reduction in microglia surveillance (Fig. 8e, f). The effect persisted for approx. 10 minutes after the light was turned off, after which surveillance slowly recovered. In the absence of **OptoLat**, pulsed blue light had no effect on surveillance (Fig. 8e, f)^37^ These experiments demonstrate that **OptoLat** can not only be used in single layer cell culture, but also in complex tissue. The compound is well tolerated and does not seem to trigger an adverse immune response in the dark, which would have caused the retraction of microglia processes during the **OptoLat** incubation period.

**Figure 8:**
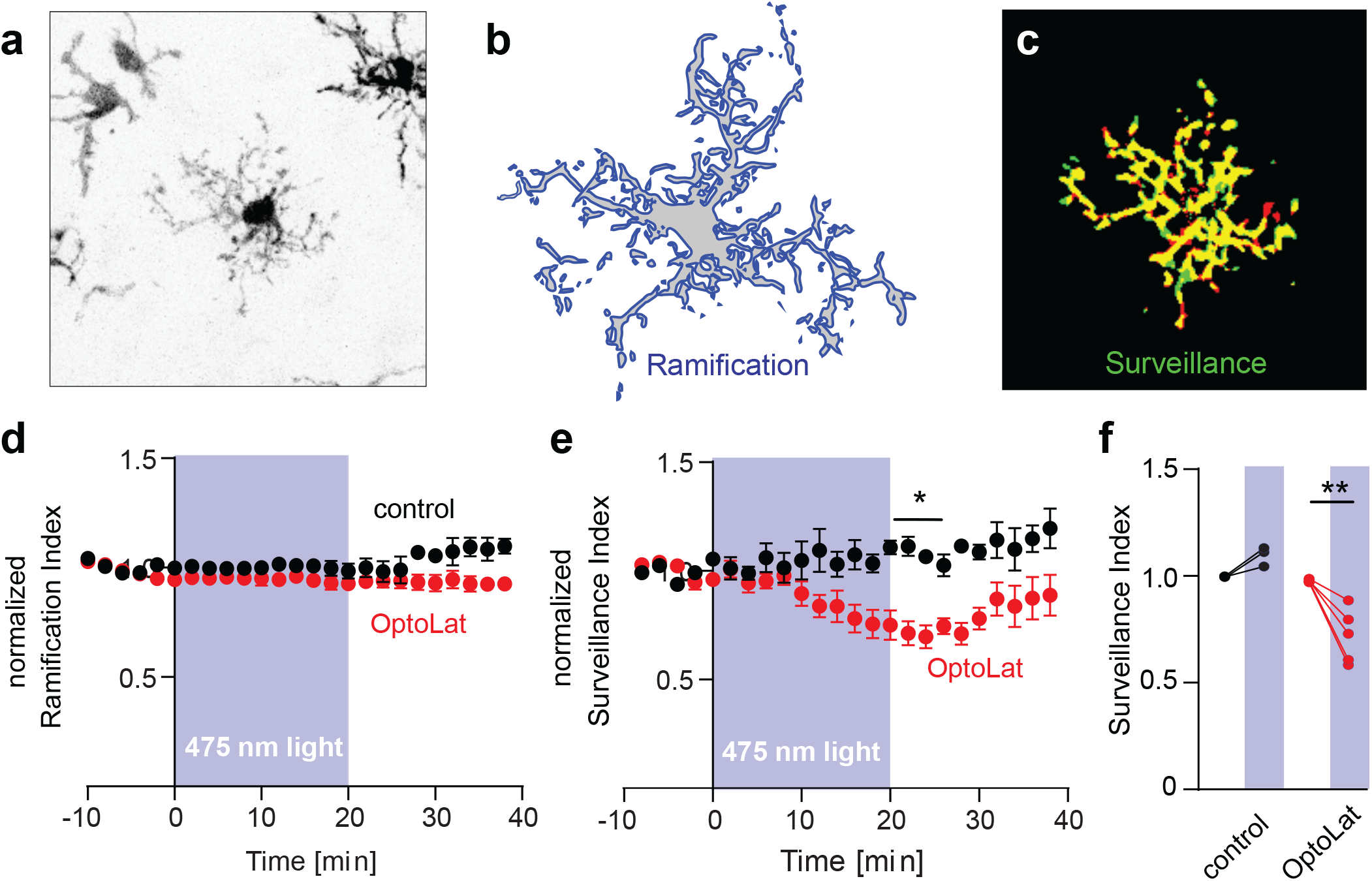
Light-induced reduction of microglia surveillance in brain slice culture. a) Fluorescence image (contrast inverted) of microglia in hippocampal tissue culture expressing tdTomato. b) Ramification of microglia was calculated by dividing its perimeter by the surface area. c) Surveillance was quantified as the sum of areas covered only at one time point (red + green pixels). d) Ramification of microglia was not affected upon activation of OptoLat. Red: Tissue incubated with OptoLat and illuminated for 20 min. Black: Tissue without OptoLat, illuminated for 20 min. Mean ± SEM, normalized to baseline. e) Surveillance index of microglia incubated with OptoLat (red) and without OptoLat (black). Reduction of surveillance after ca 10 minutes irradiation in presence of OptoLat, followed by recovery. Mean ± SEM, normalized to baseline. f) Statistical analysis of light-induced changes in surveillance index in cultures without (control, n = 3 cells/slice cultures) and with OptoLat (n = 5 cells/slice cultures). Baseline (−10 to 0 min) was compared to post-light period (20-22 min). 2-way ANOVA with Bonferroni Post-hoc test. Adjusted p-value: 0.0028.

## DISCUSSION

Photoswitchable modulators of the cytoskeleton allow for light-induced precise spatial and temporal modulation of cytoskeletal dynamics. Recently, we have developed light-responsive stabilizers of F-actin derived from the natural product jasplakoinlide.^10,11^ Here, we report generation of a light-activated actin destabilizer, the photoswitchable latrunculin derivative **OptoLat**.

Our design of photoswitchable latrunculins was based on the known SAR and several X-ray structures of latrunculin A/B bound to G-actin. This also enabled computational docking studies that pointed to the need for a large photoswitch for optimal difference in activity. The systematic evaluation of candidates with the desired photophysical and thermal properties ultimately resulted in our most effective compound, **OptoLat**. This blue-light sensitive, fast-relaxing photoswitch allows for the disassembly of actin structures with high spatiotemporal precision. It is cell permeant and performs well in a variety of cell types, such as yeast cells, human cancer cell lines, rodent oligodendrocytes, and microglia embedded in rodent brain slices. It extends the toolset available for the temporally and spatially restricted modulation of actin dynamics and complements other optogenetic^38,39^ and photopharmacological^10,11^ approaches toward controlling the actin cytoskeleton. Its ease of application, enabled by its inactivity in the dark and its rapid, reversible activation by irradiation with blue light, opens a myriad of possible applications, from *in vitro* and cellular assays to the investigation of actin dynamics in more complex tissues, or organisms.

Finally, **OptoLat** serves as a template for “latruncologs”, i.e. structurally simplified latrunculin A/B derivatives that are synthetically accessible in ca. 10 steps. The stable aryl ether moiety can be introduced via Mitsunobu reaction of nucleophilic aromatic substitution enabling the synthesis of a wide range of analogs that can be easily diversified. The systematic exploration of these analogs, which can be further functionalizes, e.g. with fluorophores, is under active investigation in out laboratories.

## ACKNOWLEDGEMENTS

This work was supported by grants from the National Institutes of Health (NIH) (GM122589 and AG051047) to LP. These studies used the Confocal and Specialized Microscopy Shared Resource of the Herbert Irving Comprehensive Cancer Center at Columbia University Irving Medical Center, funded in part through the NIH/NCI Cancer Center Support Grant P30CA013696. We thank the Studienstiftung des deutschen Volkes for a PhD fellowship (to N.A.V.) and the National Institutes of Health (Grant R01GM126228, to D.T.) for financial support. We also thank the Stanford Medical Scientist Training Program [T32 GM007365-45] and Stanford Bio-X Interdisciplinary Graduate Fellowship (M.H.C.), the McKnight Endowment Fund for Neuroscience (J.B.Z.), the National Multiple Sclerosis Society Harry Weaver Neuroscience Scholar Award (J.B.Z.), the Myra Reinhard Family Foundation (J.B.Z.), the National Institutes of Health R01NS119823 (J.B.Z.), and the Koret Family Foundation (J.B.Z.).

