## Supplementary Data for "Optical Control of G-Actin with a Photoswitchable Latrunculin"

for

### General Methods, Chemistry

All reactions were carried out under magnetic stirring and if air and/or moisture sensitive in flame-dried glass ware using standard *Schlenk* technique. Temperatures for reaction conditions were recorded as external bath temperature of heating blocks or heating baths. Low temperature reactions were carried out in a *Dewar* vessel using acetone/ dry ice (–78 °C), DI water/ NaCl (–10 °C), DI water/ ice (0 °C). When low temperatures were maintained for extended periods of time, a JULABO FT902 – Cryostat 9650892 with an isopropanol cooling bath was used.
High temperature reactions were conducted using metal heating blocks, silicon oil baths or sand baths. When reactions were heated to temperatures near boiling point, reflux condensers or pressure tubes were used.
Dry solvents were purchased as Acros Organics as Extra dry reagents over 4 Å molecular sieves or dried by passing the degassed solvents through activated alumina columns. Reagents were distilled according to standard procedures.
All reagents were purchased from commercial sources (ABCR, Acros Organics, Alfa Aesar, Ark Pharm, Combi-Blocks, Fisher Scientific, Oakwood, OxChem, Sigma Aldrich, Strem, TCI) and used without further purification unless otherwise noted.
Reactions were monitored using thin analytical thin layer chromatography (TLC) or liquid chromatography- mass spectrometry (LCMS) as specified below.

**Analytical Thin Layer Chromatography** (TLC) was performed on glass plates precoated with silica gel (0.25 mm, 60-Å pore size, Merck) and visualized under UV light at 254 nm/ 366 nm. Staining was performed with aqueous Ceric Ammonium Molybdate (IV) solution (CAM; 2.0 g $Ce(NH_4)_4(SO_4)_4 \cdot 2\ H_2O$, 48 g $(NH_4)_6Mo_7O_{24} \cdot 4\ H_2O$, 60 ml conc. sulfuric acid, 940 ml $H_2O$), or aqueous $KMnO_4$ solution (7.5 g $KMnO_4$, 50 g $K_2CO_3$, 6.25 ml aqueous 10% NaOH, 1000 ml $H_2O$) and subsequent heating using a heat gun (150–600 °C).

**Liquid Chromatography-Mass Spectrometry** (LCMS) was performed on an Agilent Technologies 1260 II Infinity system with an LC Kinetex column 2.6 µm C18 (50 x 3 mm) connected to an Agilent Technologies 6120 Quadrupole mass spectrometer with ESI ionization source.

**Flash Column Chromatography** was performed on Merck Silica Gerudan (60 Å pore size, 40–63 µm, Merck KGaA) and either performed manually with positive nitrogen or air pressure or using a Teledyne Isco Combiflash EZprep flash purification system.

**High Pressure Liquid Chromatography** (HPLC) was performed on a 260 Infinity Agilent Technologies system with a semipreparative column Phenomenex Gemini 5 µm C18 (150 x 10 mm) flowrate 8 ml/min and preparative column Phenomenex Gemini 5 µm C18 (150 x 30 mm) flowrate 72 ml/min; mobile phase gradients of acetonitrile in $H_2O$ (each containing 0.1% formic acid unless otherwise specified); UV-Vis detection with a 0.3 mm preparative flow cell.

**Nuclear Magnetic Resonance Spectra** (NMR) were recorded on a Bruker Avance III HD 400 MHz spectrometer equipped with a CryoProbeTM (400 MHz for $^{1}H$ and 101 MHz for $^{13}C$ spectra), a Bruker Avance III 400 NMR Spectrometer (377 MHz for $^{19}F$ and 400 MHz for high temperature $^{1}H$ spectra) and a Bruker Avance 600 MHz spectrometer equipped with a CPTCI-cryoprobehead (600 MHz $^{1}H$ and 150 MHz $^{13}C$ spectra)
Chemical shift signals are reported as parts per million (ppm, *δ* scale) and referenced to the residual non deuterated solvent signals. NMR spectra were processed and analyzed using MestReNova (Mestrelab Research; Versions 11 and 14)

**High-Resolution Mass Spectrometry** (HRMS) was performed on an Agilent Technologies 6224 Accurate-Mass TOF/LC/MS spectrometer with electrospray ionization (ESI) or atmospheric pressure chemical ionization (APCI) ionization sources.

**Infrared Spectroscopy** (IR) was performed in a ThermoScientific Nicolet-6700 Fourier Transform Infrared Spectrometer (FTIR) and the absorption signals are reported in wavenumbers [$cm^{-1}$].

**Optical Rotation** was determined on a Jasco P-2000 Polarimeter using the Sodium-D-line (598 nm) at room temperature (*T* = 20 – 28 °C) and concentration (*c* in g/mol); path length l = 1 dm; HPLC grade solvents were used. Specific optical rotations were calculated following:

$$[\alpha]_D^T = \frac{\alpha}{c \cdot d}\frac{10^{-1} \cdot \mathrm{deg} \cdot \mathrm{cm}^2}{\mathrm{g}}$$

Optical properties can be influenced by the *cis*:*trans* ratio of azobenzene and diazocine photoswitches and are therefore not reported for photoswitch containing reaction products.

**UV-Vis Spectra** was performed using a Varian Cary 60 UV-Visible Spectrophotometer using disposable UV-cuvettes (BRAND UV-Cuvette Disposable Spectrophotometer/Photometer Cuvettes, BrandTech; Ultra-Micro Cuvettes Vol. 70 – 850 µl, window height 8.5 mm, pathlength 10 mm; VWR cat# 47743-834)). Sample temperatures were controlled using an Agilent Technologies PCB 1500 Water Peltier system and samples were irradiated orthogonally using a Cairn Research Optoscan Monochromator with Optosource High Intensity Arc Lamp with a 75 W UXL-S50A lamp from USHIO Inc. Japan (15 nm full width at half maximum) or prizmatix (U)HP-LEDs: 365 nm, 415 nm, 460 nm, 520 nm.
Thermal half-lives were determined by first order exponential decay fit using GraphPad Prism 9 for macOS (San Diego, CA, USA).

**In-Situ NMR irradiation** was achieved with HP and UHP LEDs connected to an optical fiber (with the last 7 cm stripped, last 5 cm sanded) that was inserted into the compound solution in DMSO-$d_6$ using a coaxial NMR insert.

**All yields** are isolated yields, unless otherwise specified.

### Synthetic Protocols

#### Synthesis of Hex-5-enal:

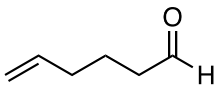

A flame-dried 500 ml *Schlenk* flask was equipped with a stir bar and charged with oxalyl chloride (10.4 ml, 121 mmol, 1.50 eq.) in DCM (90 ml). The solution was cooled to –78 °C and DMSO (17.0 ml, 240 mmol, 3.00 eq.) was added over 10 minutes. The reaction mixture was stirred for 10 min. before a solution of hex-5-enol (8.00 g, 79.9 mmol, 1.00 eq.) in DCM (15 ml) was added over 15 min. After 1.5 h, $NEt_3$ (33.3 ml, 240 mmol, 3.00 eq.) was added dropwise. Upon addition, the reaction was allowed to warm to 0 °C and water (150 ml) was added. The mixture was stirred for 45 min. at 0 °C, the layers were separated and the aqueous layer was extracted with DCM (2 x 150 ml). The combined organic layers were washed with LiCl (10% aq., 3 x 100 ml), dried over $MgSO_4$, filtered and concentrated (700 mbar, 35 °C water bath). Purification was performed by flash column chromatography (Pent:$Et_2O$ = 10:0 to 9.5:0.5). Upon slow evaporation, residual solvent was removed by fractional distillation. The title compound was obtained as a colorless, volatile liquid (4.00 g, 40.8 mmol, 51%).

**$R_f$** ($SiO_2$, Pent:$Et_2O$ = 9:1; $KMnO_4$) = 0.5

**$^1$H-NMR (400 MHz, $CDCl_3$):** $\delta$ [ppm] = 9.78 (d, $J$ = 1.8 Hz, 1H), 5.77 (ddt, $J$ = 16.9, 9.8, 6.7 Hz, 1H), 5.14 – 4.91 (m, 2H), 2.45 (t, $J$ = 7.3 Hz, 2H), 2.10 (q, $J$ = 7.2 Hz, 2H), 1.74 (p, $J$ = 7.4 Hz, 2H).

**$^{13}$C-NMR (101 MHz, $CDCl_3$):** $\delta$ [ppm] = 202.7, 137.7, 115.7, 43.3, 33.1, 21.3.

The analytical data matched those previously reported.[1]

---

[1] Lininger, M.; Neuhaus, C.; Hofmann, T.; Fransioli-Ignazio, L.; Jordi, M.; Drueckes, P.; Trappe, J.; Fabbro, D.; Altmann, K.-H. *ACS Med. Chem. Lett.* **2011**, *2*, 22–27.

**Synthesis of (*R*)-4-isopropylthiazolidine-2-thione:**

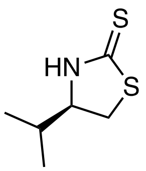


D-Valinol (10.6 g, 0.11 mol, 1.00 eq.) was dissolved in EtOH (30 ml) and $CS_2$ (15.7 ml, 262 mmol, 2.60 eq.) was added. A solution of KOH (15.3 g, 272 mmol, 2.70 eq.) in EtOH:$H_2O$ (1:1, 120 ml) was added dropwise at room temperature and the reaction was then heated to 105 °C for 3 d. The reaction was cooled to room temperature and $N_2$ was lead through the solution for 1 h to remove volatile byproducts, which were quenched by passing through a sodium hypochloride solution. All volatiles were removed under reduced pressure and the residues were diluted with DCM (200 ml) and acidified with HCl (120 ml, 1.0 M) under cooling and vigorous stirring. The layers were separated, and the aqueous layer was extracted with dichloromethane (3 x 200 ml). The combined organic layers were washed with brine, dried over $MgSO_4$, filtered and concentrated. (*R*)-4-isopropylthiazolidine-2-thione was obtained as yellow solid (14.0 g, 86.8 mmol, 86%) and used without further purification.

**$R_f$** ($SiO_2$, Hex:EtOAc = 5:1, $KMnO_4$) = 0.3

**$^1$H-NMR (400 MHz, $CDCl_3$):** $\delta$ [ppm] = 8.28 (s, 1H), 4.05 (td, $J$ = 8.2, 6.4 Hz, 1H), 3.49 (dd, $J$ = 11.1, 8.3 Hz, 1H), 3.30 (dd, $J$ = 11.1, 8.2 Hz, 1H), 1.97 (h, $J$ = 6.7 Hz, 1H), 1.01 (d, $J$ = 6.8 Hz, 3H), 0.98 (d, $J$ = 6.8 Hz, 3H).

**$^{13}$C-NMR (101 MHz, $CDCl_3$):** $\delta$ [ppm] = 201.2, 70.2, 36.1, 32.1, 18.9, 18.4.

The analytical data matched those previously reported.[2]

---

[2] Kasun, Z. A.; Gao, X.; Lipinski, R. M.; Krische, M. J. *J. Am. Chem. Soc.* 2015, 137, 8900–8903.

**Synthesis of Crimmins' Auxiliary:**

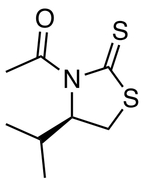

NaH (3.82 g, 95.5 mmol, 1.10 eq.) was dissolved in dry THF (45 ml), degassed and cooled to 0 °C. A solution of (*R*)-4-isopropylthiazolidine-2-thione (14.0 g, 86.8 mmol, 1.00 eq.) in THF (45 ml) was added slowly. After stirring for 20 min. at 0 °C, acetyl chloride (6.80 ml, 95.3 mmol, 1.10 eq.) was added dropwise to the reaction. After stirring at 0 °C for 20 min. the reaction mixture was allowed to warm to room temperature. Full consumption of the starting material was observed by TLC after 1 h. HCl (100 ml, 1 M) and Ethyl acetate (100 ml) were added. The layers were separated, and the aqueous layer was extracted with Ethyl acetate (2 x 100 ml). The combined organic layers were washed with brine, dried over $MgSO_4$ filtered and concentrated. After purification on flash column chromatography ($SiO_2$; Hex:EtOAc = 20:1 to 10:1) Crimmins' auxiliary was obtained as a yellow oil (14.6 g, 71.8 mmol, 83%).

$\mathbf{R}_f$ ($SiO_2$, Hex:EtOAc = 9:1; $KMnO_4$) = 0.5

**$^1$H-NMR (400 MHz, $CDCl_3$):** $\delta$ [ppm] = 5.15 (dd, $J$ = 7.9, 6.3 Hz, 1H), 3.50 (dd, $J$ = 11.5, 8.1 Hz, 1H), 3.02 (d, $J$ = 11.3 Hz, 1H), 2.77 (s, 3H), 2.37 (h, $J$ = 6.9 Hz, 1H), 1.06 (d, $J$ = 6.8 Hz, 3H), 0.97 (d, $J$ = 6.9 Hz, 3H).

**$^{13}$C-NMR (100 MHz, $CDCl_3$):** $\delta$ [ppm] = 203.4, 170.9, 71.4, 30.9, 30.5, 27.1, 19.2, 17.9.

**HRMS (ESI$^+$, *m/z*):** $[M+H]^+$ for $C_8H_{13}NOS_2^+$ calcd.: 203.0439, found: 203.0432.

**IR (ATR):** $\tilde{\nu}$ [cm$^{-1}$] = 2962 (w), 2874 (w), 1692 (s), 1467 (w), 1410 (w), 1363 (s), 1305 (m), 1271 (s), 1244 (s), 1201 (s), 1099 (m), 1072 (s), 1030 (s), 1008 (s), 973 (s), 922 (m), 898 (w), 884 (w), 855 (s), 802 (w), 778 (w), 755 (w).

$[\alpha]_D$ = –355 ($c$ = 0.01, $CHCl_3$)

The analytical data matched those previously reported.[3]

**Synthesis of (R)-3-hydroxy-1-((R)-4-isopropyl-2-thioxothiazolidin-3-yl)oct-7-en-1-one (SI-1):**

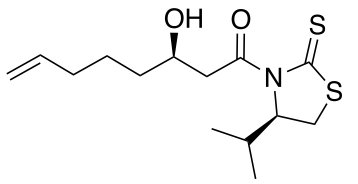


In a flame-dried *Schlenk* flask Crimmin's auxiliary (2.90 g, 14.3 mmol, 1.50 eq.) was dissolved in dry distilled DCM (75 ml), degassed and cooled to –50 °C. After keeping the starting material at –50 °C for 10 min., a solution of $TiCl_4$ in DCM (1.72 ml of a 1.0 M solution, 15.2 mmol, 1.60 eq.) was added dropwise and the reaction was stirred for 20 min. Then a solution of DIPEA (2.63 ml, 15.2 mmol, 1.60 eq.) in DCM (25 ml) was added dropwise and the reaction was warmed to –40 °C and stirred for 1 h. Next, the reaction was cooled to –78 °C and stirred for 10 min. Hex-5-enal (0.93 g, 9.51 mmol, 1.00 eq.) was added dropwise as a solution in DCM (24 ml) whereupon the reaction was warmed to –50 °C and stirred for 16 h. The reaction was quenched with pH = 7.5 phosphate buffer solution and after separation of the two layers, the aqueous layer was extracted with DCM (3 x 100 ml). The combined organic layers were washed with brine, dried over $MgSO_4$, filtered and evaporated. Purification by flash column chromatography ($SiO_2$, Hex:EtOAc = 20:1 to 5:5) furnished the title compound as a yellow oil (1.90 g, 6.31 mmol, 66%).

**$R_f$** ($SiO_2$, Hex:EtOAc = 7:3, $KMnO_4$) = 0.2

**$^1$H-NMR (400 MHz, $CDCl_3$):** $\delta$ [ppm] = 5.81 (ddt, *J* = 17.0, 10.2, 6.7 Hz, 1H), 5.16 (ddd, *J* = 7.6, 6.2, 1.1 Hz, 1H), 5.09 – 4.92 (m, 2H,), 4.13 (dddd, *J* = 9.8, 7.4, 4.6, 2.4 Hz, 1H,), 3.64 (dd, *J* = 17.7, 2.4 Hz, 1H), 3.52 (dd, *J* = 11.5, 7.9 Hz, 1H), 3.12 (dd, *J* = 17.7, 9.4 Hz, 1H), 3.03 (dd, *J* =

[3] McNutley, J.; McLeod, D.; Jenkins, H. *A. Eur. J. Org. Chem.* **2016**, 688–692.

11.5, 1.1 Hz, 1H), 2.51 ($s_{br}$, 1H), 2.36 (dq, $J$ = 13.6, 6.8 Hz, 1H), 2.09 (dt, $J$ = 7.9, 6.5, 2H), 1.66 – 1.39 (m, 4H), 1.07 (d, $J$ = 6.9 Hz, 3H), 0.98 (d, $J$ = 7.0 Hz, 3H).

**$^{13}$C-NMR (100 MHz, CDCl$_3$):** $\delta$ [ppm] = 203.2, 173.4, 138.7, 114.9, 71.5, 68.0, 45.7, 35.9, 33.7, 31.0, 30.7, 25.0, 19.2, 18.0.

**HRMS (ESI$^+$, *m/z*):** $[M+H]^+$ for $C_{14}H_{23}NO_2S_2^+$: calcd.: 302.1243, found: 302.1244.

**IR (ATR):** ṽ [cm$^{-1}$] = 3430 (w), 2963 (w), 1930 (w), 1684 (m), 1640 (w), 1468 (w), 1437 (w), 1391 (w), 1364 (m), 1305 (m), 1279 (m), 1256 (s), 1155 (s), 1122 (m), 1092 (s), 1037 (s), 990 (m), 910 (m), 883 (m), 752 (s), 722 (m).

$[\alpha]_D$ = –326 ($c$ = 0.02, CHCl$_3$)

The analytical data matched those previously reported.[4]

**Synthesis of (R)-3-((tert-butyldimethylsilyl)oxy)-1-((R)-4-isopropyl-2-thioxothiazolidin-3-yl)oct-7-en-1-one (SI-2):**

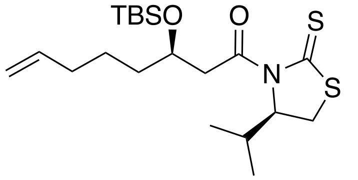


**Aldol product SI-1** (3.07 g, 11.8 mmol, 1.00 eq.) was dissolved in dry DCM (62 ml) and cooled to –78 °C. 2,6-Lutidine (4.10 ml, 35.5 mmol, 3.00 eq.) and TBSOTf (4.10 ml, 17.8 mmol, 1.50 eq.) were added dropwise. The reaction was stirred at –78 °C for 1 h. Upon addition of sat. aq. $NH_4Cl$ solution (60 ml) the reaction was let come to room temperature. The layers were separated and the aqueous layer was extracted with DCM (3 x 80 ml). The combined organic layers were

[4] Symkenberg, G.; Kalesse, M. *Angew. Chem. Int. Ed.* **2014**, *53*, 1795 –1798.

washed with brine, dried over $MgSO_4$, filtered and evaporated. Purification by flash column chromatography ($SiO_2$; Hex:EtOAc = 30:1 to 10:1) furnished the title compound as yellow oil (4.00 g, 10.7 mmol, 90%).

**$R_f$** ($SiO_2$, Hex:EtOAc = 10:1) = 0.5

**$^1$H-NMR (400 MHz, $CDCl_3$):** $\delta$ [ppm] = 5.79 (ddt, $J$ = 16.9, 10.2, 6.6 Hz, 1H), 5.08 – 4.92 (m, 2H), 4.35 – 4.25 (m, 1H), 3.57 (dd, $J$ = 17.0, 7.8 Hz, 1H), 3.47 (dd, $J$ = 11.5, 7.8 Hz, 1H), 3.14 (dd, $J$ = 17.0, 4.1 Hz, 1H), 3.02 (dd, $J$ = 11.5, 1.1 Hz, 1H), 2.45 – 2.30 (m, $J$ = 6.8 Hz, 1H), 2.10 – 1.98 (m, 2H), 1.56 – 1.39 (m, 4H), 1.06 (d, $J$ = 6.8 Hz, 3H), 0.97 (d, $J$ = 7.0 Hz, 3H), 0.85 (s, 9H), 0.07 (s, 3H), 0.03 (s, 3H).

**$^{13}$C-NMR (100 MHz, $CDCl_3$):** $\delta$ [ppm] = 203.0, 172.2, 138.8, 114.7, 71.8, 69.2, 45.6, 37.2, 33.9, 31.0, 30.9, 26.0, 24.3, 19.3, 18.2, 18.0, –4.3, –4.5.

**HRMS (ESI$^+$, *m/z*):** $[M+H]^+$ for $C_{20}H_{38}NO_2S_2Si^+$: calcd.: 416.2108, found: 416.2113.

**IR (ATR):** ṽ [$cm^{-1}$] = 2928 (m), 2855 (w), 1696 (m), 1641 (w), 1471 (w), 1462 (w), 1391 (w), 1371 (m), 1306 (m), 1279 (m), 1254 (s), 1158 (s), 1123 (m), 1190 (s), 1038 (s), 1005 (m), 990 (m), 939 (m), 910 (m), 869 (m), 834 (s), 809 (s), 774 (s), 733 (m).

**Synthesis of methyl (R)-3-((tert-butyldimethylsilyl)oxy)oct-7-enoate (SI-3):**

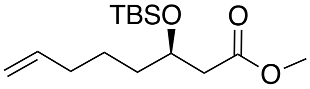


**SI-2** (6.49 g, 15.6 mmol, 1.00 eq.) was dissolved in MeOH (213 ml) and imidazole (14.3 g, 210 mmol, 13.5 eq.) was added. The reaction was stirred at room temperature over night and $NH_4Cl$ (sat. aq., 100 ml) and ethyl acetate (100 ml) were added and the layers separated. The aqueous phase was extracted with ethyl acetate (3 x 100 ml) and the combined organic layers were washed with brine, dried over $MgSO_4$, filtered and concentrated. Purification by flash column chromatography ($SiO_2$, Hex:EtOAc = 40:1 to 10:1) gave methyl ester **12** as a pale-yellow oil (4.02 g, 14.0 mmol, 90%).

**$R_f$** ($SiO_2$, Hex:EtOAc = 20:1; $KMnO_4$) = 0.5

**$^1$H-NMR (400 MHz, $CDCl_3$):** $\delta$ [ppm] = 5.79 (ddt, $J$ = 16.9, 10.1, 6.6 Hz, 1H), 5.04 – 4.92 (m, 2H) 4.18 – 4.09 (m, 1H), 3.66 (s, 3H), 2.44 (dd, $J$ = 6.3, 4.7 Hz, 2H), 2.05 (q, $J$ = 6.5, 5.9 Hz, 2H), 1.55 – 1.33 (m, 4H), 0.86 (s, 9H), 0.06 (s, 3H), 0.03 (s, 3H).

**$^{13}$C-NMR (100 MHz, $CDCl_3$):** $\delta$ [ppm] = 172.4, 138.8, 114.8, 69.4, 51.6, 42.7, 37.1, 33.8, 25.9, 24.3, 18.1, –4.4, –4.7.

**HRMS (ESI$^+$, *m/z*):** $[M+H]^+$ for $C_{15}H_{31}O_3Si^+$: calcd.: 287.2037, found: 287.2040.

**IR (ATR):** ṽ [cm$^{-1}$] = 2929 (m), 2857 (w), 1740 (m), 1642 (w), 1473 (w), 1463 (w), 1437 (w), 1361 (w), 1252 (m). 1199 (m), 1168 (m), 1081 (m), 1033 (m), 1005 (m), 939 (w), 910 (m), 834 (s), 810 (m), 774 (s).

$[\alpha]_D$ = –14 ($c$ = 0.01, $CHCl_3$)

**Synthesis of (R)-3-((tert-butyldimethylsilyl)oxy)oct-7-enal (16):**

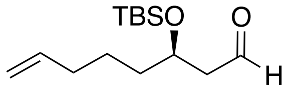


**SI-3** (4.01 g, 14.0 mmol, 1.00 eq.) was dissolved in dry DCM (212 ml) and cooled to –78 °C. DIBAL-H (25% in toluene, 18.8 ml, 28.0 mmol, 2.00 eq.) was added dropwise. The reaction as stirred for 2 h at –78 °C, before MeOH (80 ml) was added slowly and the reaction was stirred for another 30 min. Next, aqueous Rochelle's salt solution (200 ml) was added and the reaction was warmed to room temperature. The layers were separated and the aqueous phase was extracted with DCM (3 x 200 ml). The combined organic layers were washed with brine, dried over $MgSO_4$, filtered and concentrated. Purification of the crude product was performed by flash column chromatography ($SiO_2$, Hex:EtOAc = 30:1) and gave the title compound as a colorless oil (3.23 g, 12.6 mmol, 90%).

**$R_f$** ($SiO_2$, Hex:EtOAc = 30:1) = 0.26

**$^1$H-NMR (400 MHz, $CDCl_3$):** $\delta$ [ppm] = 9.81 (t, $J$ = 2.5 Hz, 1H), 5.78 (ddt, $J$ = 16.9, 10.2, 6.6 Hz, 1H), 5.05 – 4.93 (m, 2H), 4.19 (p, $J$ = 5.8 Hz, 1H), 2.52 (dd, $J$ = 5.8, 2.5 Hz, 2H), 2.05 (q, $J$ = 6.9 Hz, 2H), 1.62 – 1.36 (m, 4H), 0.87 (s, 9H), 0.07 (s, 3H), 0.05 (s, 1H).

**$^{13}$C-NMR (101 MHz, $CDCl_3$):** $\delta$ [ppm] = 202.5, 138.6, 114.9, 68.2, 51.0, 37.3, 33.7, 25.9 , 24.5, 18.1, –4.3, –4.6.

**HRMS (ESI$^+$, *m/z*):** $[M+H]^+$ for $C_{14}H_{28}O_2Si^+$: calcd.: 257.1931, found: 257.1934.

**IR (ATR):** $\tilde{\nu}$ [$cm^{-1}$] = 2929 (m), 2857 (w), 1710 (m), 1642 (w), 1473 (w), 1463 (w), 1411 (w), 1361 (w), 1298 (w), 1252 (m), 1083 (m), 1005 (m), 939 (m), 910 (m), 834 (s), 809 (m), 773 (s).

$[\alpha]_D$ = +1 ($c$ = 0.01, $CHCl_3$)

**Synthesis of ethyl (R)-2-oxothiazolidine-4-carboxylate (SI-4):**

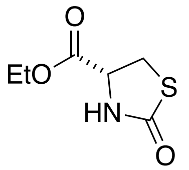


Ethyl L-cysteinate hydrochloride (5.93 g, 31.9 mmol, 1.00 eq.) was dissolved in dry THF (100 ml) and degassed/ backfilled with $N_2$. CDI (5.70 g, 35.1 mmol, 1.10 eq.) was added in portions at r.t.

and the reaction was stirred for 16 h. The reaction mixture was filtered over a plug of celite, rinsed with THF and concentrated under reduced pressure. Purification by flash column chromatography ($SiO_2$; Hex:acetone = 9:1 to 7:3) gave the title compound as a pale yellow oil (4.48 g, 25.6 mmol, 80%).

**$R_f$** ($SiO_2$, Hex:acetone = 6:2; $KMnO_4$) = 0.2

**$^1$H-NMR (400 MHz, $CDCl_3$):** $\delta$ [ppm] = 6.04 (s, 1H), 4.42 (ddd, *J* = 8.0, 5.5, 1.0 Hz, 1H), 4.28 (q, *J* = 7.1 Hz, 2H), 3.74 – 3.59 (m, 2H), 1.31 (d, *J* = 7.2 Hz, 3H).

**$^{13}$C-NMR (100 MHz, $CDCl_3$):** $\delta$ [ppm] = 174.4, 170.0, 62.6, 56.1, 31.9, 14.3.

**HRMS (ESI$^+$, *m/z*):** $[M+H]^+$ for $C_6H_{10}NO_3S^+$: calcd.: 176.0376, found: 176.0374.

The analytical data agree with the literature.[5]

**Synthesis of ethyl (*R*)-2-oxothiazolidine-4-carboxylate (SI-5):**

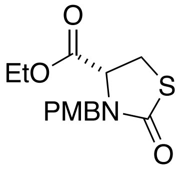


PMBCl (7.40 ml, 52.1 mmol, 120 eq.) was added dropwise to a suspension of **SI-4** (7.60 g, 43.4 mmol, 1.00 eq.), $K_2CO_3$ (8.99 g, 65.1 mmol, 1.50 eq.) and NaI (1.95 g, 13.0 mmol, 0.30 eq.) in DMF (115 ml) at r.t. After 10 h, more PMBCl (2.17 ml, 15.4 mmol, 0.70 eq.) was added and the reaction stirred for another 12 h. The reaction was diluted with Ethyl acetate (40 ml) and water (60 ml), the layers were separated, and the aqueous layer was extracted with EtOAc (2 x 50 ml). The combined organic layers were washed with LiCl (10% aq., 3 x 100 ml) and brine, dried over

[5] Fürstner, A.; De Souza, D.; Turet, L.; Fenster, M. D. B.; Parra-Rapado, L.; Wirtz, C.; Mynott, R.; Lehmann C.W. *Chem. Eur. J.* **2007**, *13*, 115 – 134.

$MgSO_4$, filtered and concentrated. The crude mixture was purified by flash column chromatography ($SiO_2$, Hex:EtOAc = 9:1 to 7:3) and the title compound was obtained as a pale-yellow oil (4.12 g, 13.9 mmol, 61%).

**$R_f$** ($SiO_2$, Hex:EtOAc = 7:3, $KMnO_4$) = 0.34

**$^1$H-NMR (400 MHz, $CDCl_3$):** $\delta$ [ppm] = 7.15 (d, $J$ = 8.3 Hz, 2H), 6.86 (d, $J$ = 8.4 Hz, 2H), 5.08 (d, $J$ = 14.8 Hz), 4.24 (q, $J$ = 7.2 Hz, 1H), 4.15 – 4.05 (m, 1H), 3.99 (d, $J$ = 14.8 Hz, 1H), 3.80 (s, 3H), 3.47 (dd, $J$ = 11.4, 8.5 Hz, 1H), 3.33 (dd, $J$ = 11.4, 3.1 Hz, 1H), 1.30 (t, $J$ = 7.1 Hz, 3H).

**$^{13}$C-NMR (100 MHz, $CDCl_3$):** $\delta$ [ppm] = 171.7, 170.1, 159.5, 129.9, 127.7, 114.4, 62.3, 59.4, 55.4, 47.4, 29.2, 14.3.

**HRMS (ESI$^+$, *m/z*):** $[M+Na]^+$ for $C_{13}H_{15}NO_3SNa^+$: calcd.: 288.0665, found: 288.0678.

**IR (ATR):** $\tilde{\nu}$ [cm$^{-1}$] = 3460 (w), 2936 (w), 2836 (w), 1739 (m), 1739 (m) 1673 (s), 1611 (m), 1585 (w), 1512 (s), 1442 (m), 1390 (m), 1302 (m), 1244 (s), 1210 (s), 1172 (s), 1110 (m), 1027 (s), 814 (m). 760 (m), 724 (m).

**Synthesis of (*R*)-N-methoxy-3-(4-methoxybenzyl)-*N*-methyl-2-oxothiazolidine-4-carboxamide (SI-6):**

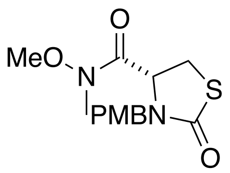


Ethyl (*R*)-2-oxothiazolidine-4-carboxylate (**SI-5**) (9.69 g, 32.8 mmol, 1.00 eq.) was dissolved in dioxane (54 ml) and a solution of KOH (5.52 g, 98.4 mmol, 3.00 eq.) in $H_2O$ (36 ml) was added. After 1.5 h, the reaction was diluted with diethyl ether (50 ml) and the pH was adjusted to 2 with aq. HCl (aq., 1 M). The layers were separated and the aqueous phase was extracted with $Et_2O$

(3 x 50 ml). The combined organic layers were washed with brine, dried over $MgSO_4$, filtered and concentrated. The carboxylic acid was used crude and therefore redissolved in dry acetonitrile (129 ml) and degassed. EDCI (9.43 g, 49.2 mmol, 1.50 eq.) and MeNH(OMe) · HCl (3.84 g, 39.4 mmol, 1.20 eq.) were added at r.t. and the reaction was stirred for 14 h. Water (100 ml) and ethyl acetate (100 ml) were added. The layers were separated and the aqueous layer was extracted with ethyl acetate (3 x 100 ml). The combined organic layers were washed with brine, dried over $MgSO_4$, filtered and concentrated. The colorless oil was purified by flash column chromatography ($SiO_2$, Hex:acetone = 8:2 to 7:3) furnishing the title compound (6.90 g, 22.2 mmol, 68% over 2 steps) as a colorless oil.

**$R_f$** ($SiO_2$, Hex:acetone = 7:3; $KMnO_4$) = 0.7

**$^{1}$H-NMR (400 MHz, $CDCl_3$):** $\delta$ [ppm] = 7.21 – 7.12 (m, 2H), 6.93 – 6.79 (m, 2H), 5.14 (d, $J$ = 14.7 Hz, 1H), 4.40 (dd, $J$ = 8.9, 5.2 Hz, 1H), 3.85 (d, $J$ = 14.7 Hz, 1H), 3.79 (s, 3H), 3.53 – 3.41 (m, 1H), 3.38 (s, 3H), 3.21 (s, 3H), 3.15 (dd, $J$ = 11.2, 5.0 Hz, 1H).

**$^{13}$C-NMR (100 MHz, $CDCl_3$):** $\delta$ [ppm] = 172.4, 169.4, 159.5, 130.2, 127.8, 114.3, 61.4, 57.6, 55.5, 47.1, 32.7, 28.2.

**HRMS (ESI$^+$, *m/z*):** $[M+Na]^+$ for $C_{14}H_{18}N_2O_4SNa^+$: calcd.: 333.0879, found: 333.0878.

**IR (ATR):** $\tilde{\nu}$ [cm$^{-1}$] = 2937 (w), 2836 (w), 1669 (s), 1611 (m), 1585 (w), 1512 (s), 1441 (m), 1392 (m), 1326 (w), 1302 (m), 1245 (s), 1173 (s), 1110 (m), 1029 (m), 996 (m), 955 (m), 887 (w), 834 (m), 812 (m), 752 (m).

**Synthesis of (*R*)-4-acetyl-3-(4-methoxybenzyl)thiazolidin-2-one (17):**

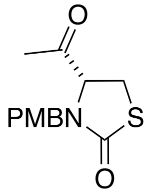


Weinreb amide **SI-6** (6.84 g, 22.0 mmol, 1.00 eq.) was dissolved in dry distilled THF (90 ml) and cooled to 0 °C. A solution of MeMgBr (29.4 ml, 88.2 mmol, 4.00 eq.; 3.0 M in $Et_2O$) was added dropwise and the reaction was stirred at 0 °C for 2 h. $NH_4Cl$ (sat. aq., 60 ml) at 0 °C. The reaction was warmed to room temperature and diluted with $H_2O$ (60 ml) and EtOAc (100 mL). The layers were separated and the organic layer was extracted with ethyl acetate (3 x 100 ml). The combined organic layers were washed with brine, dried over $MgSO_4$, filtered and concentrated. The title compound was obtained as an off-white solid (5.60 g, 21.1 mmol, 96%) and used without further purification.

$\mathbf{R}_f$ ($SiO_2$, Hex:acetone = 7:3; $KMnO_4$) = 0.6

**$^1$H-NMR (400 MHz, $CDCl_3$):** $\delta$ [ppm] = 7.13 (d, $J$ = 8.2 Hz, 2H), 6.89 – 6.82 (m, 2H), 5.02 (d, $J$ = 14.6 Hz, 1H), 4.09 (dd, $J$ = 9.4, 3.9 Hz, 1H), 3.91 (d, $J$ = 14.7 Hz, 1H), 3.80 (s, 3H), 3.50 (dd, $J$ = 11.5, 9.4 Hz, 1H), 3.11 (dd, $J$ = 11.5, 4.0 Hz, 1H), 2.14 (s, 3H).

**$^{13}$C-NMR (100 MHz, $CDCl_3$):** $\delta$ [ppm] = 204.9, 171.8, 159.7, 130.1, 127.3, 114.5, 65.7, 55.5, 47.7, 27.9, 26.4.

**IR (ATR):** $\tilde{\nu}$ [cm$^{-1}$] = 3013 (w), 2911 (w), 2842 (w), 1719 (m), 1658 (s), 1612 (m), 1585 (w), 1512 (s), 1463 (w), 1439 (m), 1393 (m), 1356 (m), 1309 (w), 1285 (m), 1240 (s), 1202 (m), 1185 (s), 1176 (s), 1161 (s), 1151 (s), 1110 (m), 1028 (s). 1012 (m), 997 (s), 942 (m), 902 (m), 839 (m), 825 (m), 812 (s), 799 (m), 763 (m), 707 (m).

$[\alpha]_D$ = –48 ($c$ = 0.01, EtOH)

The analytical data matched those previously reported.[6]

**Synthesis of (4R)-4-((5R)-5-((tert-butyldimethylsilyl)oxy)-3-hydroxydec-9-enoyl)-3-(4-methoxybenzyl)17hiazolidine-2-one (18):**

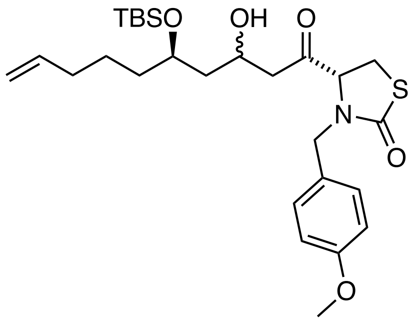


Thoazolidinone **17** (1.98 g, 7.48 mmol, 1.20 eq.) was dissolved in dry DCM (150 ml) and cooled to –78 °C. A solution of $TiCl_4$ (7.86 ml, 7.86 mmol, 1.26 eq., 1M solution in DCM) was added dropwise over 30 minutes and the reaction was stirred for 20 minutes. Next, a solution of dry DIPEA (2.30 mL, 13.4 mmol, 2.16 eq.) in dry DCM (7mL) was added dropwise over 30 minutes. The reaction was stirred at –78 °C for 1 h and then at 0 °C for 2 h, before being cooled to –60 °C. Then a solution of NMP (0.72 mL, 7.48 mmol, 1.20 eq.) in DCM (7mL) was added dropwise over 20 minutes, then stir for 0.5 h at –60 °C. A solution of aldehyde **16** (1.60 g, 6.23 mmol, 1.00 eq.) in dry DCM (7 mL) was added slowly over 1 h. The reaction was stirred at –60 °C for 16 h. $NH_4Cl$ (sat. aq., 100 ml) was added at –60 °C and the reaction was warmed to room temperature. The layers were separated, and the aqueous layer was extracted with DCM (2 x 100 mL). The combined organic layers were washed with brine, dried over $MgSO_4$, filtered and concentrated. Purification by flash column chromatography ($SiO_2$, Hex:EtOAc, 9.5:0.5 to 7:3) furnished the title compound as a light-brown oil (2.98 g, 5.67 mmol, 91%)

**R*f*** ($SiO_2$, Hex:EtOAc = 7:3; CAM) = 0.3

---

[6] Fürstner, A.; De Souza, D.; Turet, L.; Fenster, M. D. B.; Parra-Rapado, L.; Wirtz, C.; Mynott, R.; Lehmann C. W. *Chem. Eur. J.* **2007**, *13*, 115 – 134.

**Synthesis of (*R*)-4-((2*R*,4*R*,6*R*)-2,4-dihydroxy-6-(pent-4-en-1-yl)tetrahydro-2H-pyran-2-yl)-3-(4-methoxybenzyl)thiazolidin-2-one ((*R*)-SI-7) and (*R*)-4-((2*R*,4*S*,6*R*)-2,4-dihydroxy-6-(pent-4-en-1-yl)tetrahydro-2H-pyran-2-yl)-3-(4-methoxybenzyl)thiazolidin-2-one ((*S*)-SI-7):**

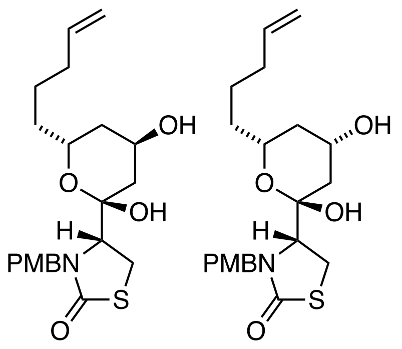


**(*R*)-SI-7** **(*S*)-SI-7**

Aldol Product **18** (1.69 g, 3.25 mmol, 1.0 eq.) was dissolved in THF (38.5 ml) and HCl (2 M, 3.86 ml, 7.71 mmol, 2.38 eq.) was added at room temperature. The reaction was stirred for 16 h until full consumption of the starting material was observed by TLC. $NaHCO_3$ (sat. aq., 50 ml) and DCM (50 ml) were added. The layers were separated, and the aqueous phase was extracted with DCM (2 x 50 ml). the combined organic layers were washed with brine, dried over $MgSO_4$, filtered and concentrated. The diastereomeric products were separated by flash column chromatography ($SiO_2$, Hex:EtOAc = 6:4 to 3:7) and (*R*)-**SI-7** was isolated as a white solid (459 mg, 1.13 mmol, 35%) and (*S*)-**SI-7** was isolated a white foam (632 mg, 1.55 mmol, 48%).

Diastereomer (*R*)-**SI-7:**

$\mathbf{R}_f$ ($SiO_2$, Hex:EtOAc = 5:5) = 0.34

**$^1$H-NMR (400 MHz, $CDCl_3$):** $\delta$ [ppm] = 7.24 – 7.16 (m, 2H), 6.88 – 6.81 (m, 2H), 5.92 – 5.75 (m, 1H), 5.24 (s, 1H), 5.13 – 4.93 (m, 3H), 4.49 – 4.44 (m, 1H), 4.34 (d, $J$ = 14.4 Hz, 1H), 4.29 – 4.16 (m, 1H), 3.80 (s, 3H), 3.64 – 3.56 (m, 1H), 3.42 – 3.28 (m, 2H), 2.34 ($s_{br}$, 1H), 2.20 – 2.02 (m, 3H), 1.90 – 1.74 (m, 2H), 1.68 – 1.42 (m, 4H).

**$^{13}$C-NMR (101 MHz, $CDCl_3$):** $\delta$ [ppm] = 173.2, 159.3, 138.7, 130.0, 130.0, 129.1, 115.0, 114.2, 100.6, 66.0, 64.8, 63.3, 55.4, 47.8, 37.9, 35.6, 33.8, 33.2, 27.1, 24.9.

Diastereomer (*S*)-**SI-7:**

$\mathbf{R_f}$ ($SiO_2$, Hex:EtOAc = 5:5) = 0.13

**$^1$H-NMR (400 MHz, $CDCl_3$):** $\delta$ [ppm] = 7.22 – 7.11 (m, 2H), 6.90 – 6.81 (m, 2H), 5.90 – 5.75 (m, 1H), 5.15 (d, $J$ = 14.5 Hz, 1H), 5.09 – 4.95 (m, 2H), 4.27 (d, $J$ = 14.5 Hz, 1H), 4.21 – 4.07 (m, 1H), 3.98 – 3.84 (m, 1H), 3.80 (s, 3H), 3.56 – 3.48 (m, 1H), 3.42 – 3.22 (m, 2H), 2.31 – 2.22 (m, 1H), 2.17 (s, 1H), 2.16 – 2.07 (m, 2H), 2.06 – 1.99 (m, 1H), 1.91 ($s_{br}$, 1H), 1.69 – 1.39 (m, 4H), 1.27 – 1.14 (m, 1H).

**$^{13}$C-NMR (101 MHz, $CDCl_3$):** $\delta$ [ppm] = 173.0, 159.3, 138.5, 129.8, 128.8, 115.1, 114.2, 101.1, 70.1, 64.8, 64.3, 55.4, 47.8, 40.8, 37.2, 35.6, 33.8, 31.1, 26.7, 25.1.

**Synthesis of (R)-4-((2R,4R,6R)-4-hydroxy-2-methoxy-6-(pent-4-en-1-yl)tetrahydro-2H-pyran-2-yl)-3-(4-methoxybenzyl)thiazolidin-2-one ((*R*)-SI-8):**

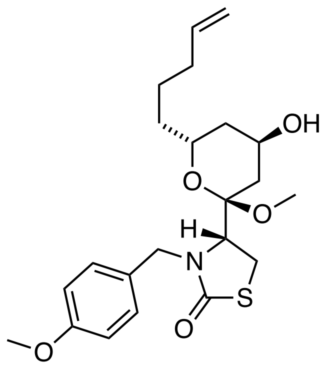


(*R*)-**SI-7** (0.37 g, 0.90 mmol, 1.00 eq.) was dissolved in MeOH (18 ml) and (+)-CSA (53.3 mg, 0.23 mmol, 0.26 eq.) was added. The reaction was stirred at r.t. for 16 h and $NaHCO_3$ (aq. sat., 30 ml) and EtOAc (50 ml) were added. The phases were separated and the aqueous phase was extracted with EtOAc (2 x 30 ml). The combined organic layers were washed with brine, dried over $Na_2SO_4$, filtered and concentrated. Purification by flash column chromatography ($SiO_2$, Hex:EtOAc = 9.5:0.5 to 5:5) furnished the title compound as a colorless solid (304 mg, 0.72 mmol, 80%).

**R*f*** ($SiO_2$, Hex:EtOAc = 4:6; CAM) = 0.33

**$^{1}$H-NMR (400 MHz, $CDCl_3$):** *δ* [ppm] = 7.19 (d, *J* = 8.6 Hz, 2H), 6.85 (d, *J* = 8.6 Hz, 2H), 5.92 – 5.78 (m, 1H), 5.12 – 4.98 (m, 3H), 4.21 (d, *J* = 14.4 Hz, 1H), 4.18 – 4.12 (m, 1H), 3.98 – 3.89 (m, 1H), 3.80 (s, 3H), 3.79 – 3.75 (m, 1H), 3.68 (d, *J* = 9.4 Hz, 1H), 3.31 – 3.21 (m, 2H), 3.16 (s, 3H), 2.21 – 2.11 (m, 2H), 2.11 – 2.04 (m, 1H), 1.90 – 1.79 (m, 2H), 1.79 – 1.39 (m, 5H).

**$^{13}$C-NMR (101 MHz, $CDCl_3$):** *δ* [ppm] = 172.9, 159.3, 138.4, 129.9, 128.7, 115.1, 114.2, 103.7, 66.3, 64.2, 59.0, 55.4, 47.8, 47.3, 37.9, 35.7, 33.9, 32.4, 25.4, 25.2.

**IR (ATR):** ṽ [$cm^{-1}$] = 3426 (w), 2938 (w), 1668 (s), 1612 (m), 1586 (w), 1512 (s), 1442 (w), 1403 (m), 1361 (w), 1303 (w), 1286 (w), 1247 (s), 1216 (m), 1199 (m), 1174 (m), 1120 (m), 1109 (m), 1074 (w), 1032 (s), 990 (m), 939 (w), 915 (m), 847 (w), 822 (w), 758 (w), 722 (w).

$[\alpha]_D$ = +34 (*c* = 0.01, $CHCl_3$)

**Synthesis of (R)-4-((2R,4S,6R)-4-hydroxy-2-methoxy-6-(pent-4-en-1-yl)tetrahydro-2H-pyran-2-yl)-3-(4-methoxybenzyl)thiazolidin-2-one ((*S*)-SI-8):**

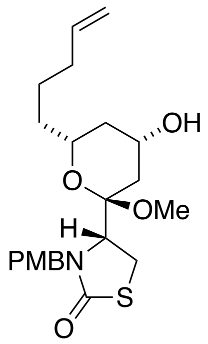

(*S*)-**SI-7** (0.56 g, 1.47 mmol, 1.00 eq.) was dissolved in MeOH (29 mL) and (+)-CSA (87.2 mg, 0.38 mmol, 0.26 eq.) was added. The reaction was stirred at r.t. for 16 h and $NaHCO_3$ (aq. sat., 30 mL) and EtOAc (50 mL) were added. The phases were separated and the aqueous layer was extracted with EtOAc (2 x 30 mL). The combined organic layers were washed with brine, dried over $Na_2SO_4$, filtered and concentrated. Purification by flash column chromatography ($SiO_2$, Hex:EtOAc = 9.5:0.5 to 5:5) furnished the title compound as a colorless foam (0.58 g, 1.37 mmol, 94%).

$\mathbf{R}_f$ ($SiO_2$, Hex:EtOAc = 4:6; CAM) = 0.29

**$^1$H-NMR (400 MHz, $CDCl_3$):** $\delta$ [ppm] = 7.24 – 7.16 (m, 2H), 6.92 – 6.77 (m, 2H), 5.85 (ddt, *J* = 16.9, 10.2, 6.7 Hz, 1H), 5.11 (d, J = 14.4 Hz, 1H), 5.09 – 4.96 (m, 2H), 4.23 (d, *J* = 14.4 Hz, 1H), 4.11 – 4.01 (m, 1H), 3.84 (dd, *J* = 8.5, 3.5 Hz, 1H), 3.80 (s, 3H), 3.63 – 3.55 (m, 1H), 3.31 – 3.21 (m, 2H), 3.06 (s, 3H), 2.22 (ddd, *J* = 12.6, 4.8, 1.8 Hz, 1H), 2.18 – 2.11 (m, 1H), 2.03 – 1.96 (m, 1H), 1.75 – 1.44 (m, 5H), 1.24 – 1.14 (m, 1H).

**$^{13}$C-NMR (100 MHz, $CDCl_3$):** $\delta$ [ppm] = 172.9, 159.3, 138.5, 130.0, 128.9, 115.1, 114.1, 103.2, 70.3, 64.8, 58.9, 55.4, 47.6, 47.3, 40.6, 37.2, 35.7, 33.9, 25. 5, 25.3.

**HRMS (ESI$^+$, *m/z*):** [$(M+Na)^+$] for $C_{33}H_{31}NO_5SNa^+$: calcd.: 444.1815, found: 444.1813.

[$(M+H)^+$] for $C_{33}H_{32}NO_5S^+$: calcd.: 422.1996, found: 422.1994.

**IR (ATR):** $\tilde{\nu}$ [$cm^{-1}$] = 3531 (w), 2920 (m), 1850 (w), 1672 (s), 1611 (w), 1586 (w), 1513 (s), 1441 (w), 1403 (w), 1363 (w), 1303 (w), 1288 (w), 1248 (m), 1215 (m), 1197 (m), 1175 (m), 1091 (m), 1030 (m), 938 (w), 907 (w), 847 (w), 821 (w), 760 (w), 728 (w).

$[\alpha]_D$ = +37 (*c* = 0.01, $CHCl_3$)

**Synthesis of (R)-4-((2R,4S,6R)-4-hydroxy-2-methoxy-6-(pent-4-en-1-yl)tetrahydro-2H-pyran-2-yl)thiazolidin-2-one (20):**

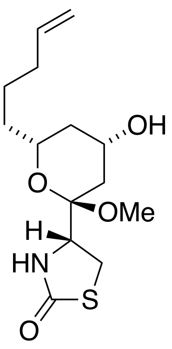

(*S*)-**SI-8** (72.0 mg, 0.17 mmol, 1.00 eq.) was dissolved in MeOH (28 mL) and MeCN (5.5 ml) and CAN (0.49 g, 0.85 mmol, 8.00 eq.) was added in one portion. The reaction was stirred for 16 h under nitrogen. $NaHCO_3$ (sat. aq., 30 ml), $H_2O$ (30 mL) and EtOAc (50 ml) were added, and the layers separated. The aqueous phase was extracted with EtOAc (2 x 50 mL). the combined organic layers were washed with brine, dried over $Na_2SO_4$, filtered and concentrated. Purification by flash column chromatography ($SiO_2$, Hex:EtOAc 7:3 to 9:1) furnished the title compound as a colorless solid (32.6 mg, 0.11 mmol, 63%, 85% brsm*.).

**The reaction stalled after 16 h and was therefore stopped, and the residual starting material was reisolated.*

$\mathbf{R_f}$ ($SiO_2$, Hex:EtOAc = 3:7; CAM) = 0.28

**$^1$H NMR (400 MHz, CDCl3):** $\delta$ [ppm] = 6.13 (s, 1H), 5.80 (ddt, $J$ = 16.9, 10.1, 6.6 Hz, 1H), 5.02 (d, $J$ = 17.0 Hz, 1H), 4.98 (d, $J$ = 10.5 Hz, 1H), 4.17 – 4.09 (m, 1H), 4.08 – 3.96 (m, 1H), 3.58 – 3.49 (m, 1H), 3.44 – 3.27 (m, 2H), 3.17 (s, 3H), 2.24 (s, 1H), 2.12 – 2.00 (m, 3H), 1.94 (ddd, $J$ = 12.5, 4.3, 2.0 Hz, 1H), 1.66 – 1.33 (m, 5H), 1.15 (q, $J$ = 11.7 Hz, 1H).

**$^{13}$C-NMR (101 MHz, $CDCl_3$):** $\delta$ [ppm] = 175.1, 138.6, 114.9, 101.6, 70.2, 64.8, 56.6, 47.9, 40.5, 36.2, 35.1, 33.8, 28.1, 25.0.

**HRMS (ESI$^+$, *m/z*):** [$(M+Na)^+$] for $NaC_{14}H_{23}NO_4S^+$: calcd.: 324.1240, found: 324.1237.

**IR (ATR):** ṽ [cm$^{-1}$] = 3221 (w), 2933 (w), 1674 (s), 1455 (w), 1365 (w), 1227 (w), 1138 (m), 1105 (m), 1029 (s), 955 (w), 909 (w), 852 (w), 720 (w).

$[\alpha]_D$ = +106 ($c$ = 0.01, $CHCl_3$)

**Synthesis of (2R,4R,6R)-2-methoxy-2-((R)-3-(4-methoxybenzyl)-2-oxothiazolidin-4-yl)-6-(pent-4-en-1-yl)tetrahydro-2H-pyran-4-yl (Z)-3-methylhepta-2,6-dienoate (SI-9):**

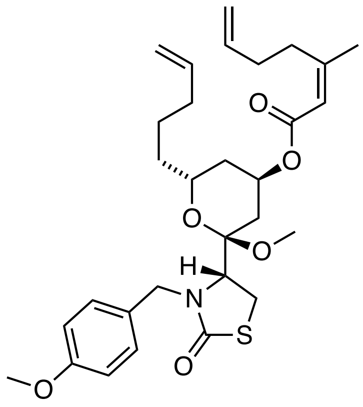


(*S*)-**SI-8** (200 mg, 0.47 mmol, 1.00 eq.) was dissolved in dry DCM (10.0 ml) and pyridine (0.08 ml, 0.95 mmol, 2.00 eq.) was added dropwise. The reaction mixture was cooled to –78 °C and $Tf_2O$ (1.68 ml, 0.57 mmol, 1.20 eq. in dry DCM (8.32 ml; 1.0 M) was added dropwise. The reaction was stirred at –30 to –50 °C for 2 h, until full conversion was observed by TLC, and turned yellow. The cold reaction mixture was poured into ice-cold 10% $KHSO_4$ solution (aq., 100 ml) and extracted with DCM (3 x 100 ml). the combined organic layers were washed with brine, dried over $Na_2SO_4$, filtered, concentrated (rotavap bath at r.t.) and dried under HV for at least 5 minutes.

In parallel, carboxylic acid **22**[7] (199 mg, 1.42 mmol, 3.00 eq.) was dissolved in dry THF (20 ml) and NaH (55.0 mg, 1.38 mmol, 2.90 eq.) was added as a suspension in THF (5 ml). The reaction mixture was heated to 55 °C for 1 h and cooled to 0 °C. The crude triflate was dissolved in dry THF (12 ml) and added dropwise, followed by the dropwise addition of [15]crown[5] until the reaction appeared clear (ca. 1.4 ml). The reaction was stirred at 0 °C for 1 h, then at r.t. for 18 h.

[7] Prepared according to She; J., Lampe; J. W., Polianski; A. B., Watson, P. S. *Tet. Lett.* **2009**, *50*, 298-301.

$NaHCO_3$ (sat. aq., 50 ml) and EtOAc (50 ml) were added and phases were separated. The aqueous layer was extracted with EtOAc (2 x 50 ml) and the combined organic layers were washed with brine, dried over $Na_2SO_4$, filtered and concentrated. The crude product was purified by flash column chromatography ($SiO_2$, Hex:$Et_2O$ = 9:1 to 7:3) and the title compound was obtained as colorless oil (235 mg, 0.43 mmol, 91%).
*Note: yield varied between 30 and 91%.

**$R_f$** (triflate **21**) ($SiO_2$, Hex:EtOAc = 9:1) = 0.18

**$R_f$** (product **23**) ($SiO_2$, Hex:EtOAc = 9:1) = 0.15

**LCMS:** (MeCN/$H_2O$, 0.1% FA: gradient 50-100% MeCN): $t_{ret}$ = 4.672 min.

**1H-NMR (400 MHz, $CDCl_3$):** δ = 7.20 (d, *J* = 8.6 Hz, 1H), 6.85 (d, *J* = 8.5 Hz, 1H), 5.84 (ddtd, *J* = 17.0, 10.9, 6.7, 4.6 Hz, 1H), 5.69 (d, *J* = 1.5 Hz, 1H), 5.19 (dd, *J* = 4.5, 2.5 Hz, 0H), 5.15 – 4.89 (m, 3H), 4.26 (d, *J* = 14.4 Hz, 0H), 3.97 – 3.86 (m, 0H), 3.80 (s, 1H), 3.80 – 3.78 (m, 0H), 3.25 – 3.20 (m, 1H), 3.10 (s, 1H), 2.73 (t, *J* = 7.8 Hz, 1H), 2.28 – 2.19 (m, 1H), 2.19 – 2.12 (m, 1H), 2.09 (dt, *J* = 15.2, 3.8, 2.0 Hz, 1H), 1.92 (dd, *J* = 11.4, 3.9 Hz, 1H), 1.89 (d, *J* = 1.3 Hz, 1H), 1.81 (dq, *J* = 16.5, 2.2 Hz, 1H), 1.77 – 1.44 (m, 2H).

**13C-NMR (101 MHz, $CDCl_3$):** *δ* [ppm] = 173.1, 165.8, 159.4, 159.1, 138.4, 137.9, 129.8, 129.7, 128.8, 117.1, 115.0, 114.9, 114.0, 114.0, 101.6, 66.1, 65.4, 59.1, 55.3, 47.5, 47.4, 35.5, 34.6, 33.9, 32.6, 32.2, 30.1, 25.4, 25.3, 25.1.

**HRMS (ESI$^+$, *m/z*):** $[M+Na]^+$ for $C_{30}H_{41}NNaO_6S^+$: calcd.: 566.2547, found: 566.2545.

**IR (ATR):** ṽ [$cm^{-1}$] = 3178 (w), 3075 (w), 2935 (w), 2860 (w), 2837 (w), 1704 (m), 1671 (s), 1641 (m), 1612 (m), 1586 (w), 1512 (s), 1442 (m), 1402 (m), 1379 (m), 1359 (m), 1329 (m), 1302 (m), 1284 (m), 1247 (s), 1215 (s), 1194 (s), 1174 (s), 1146 (s), 1127 (s), 1108 (s), 1092 (s), 1032 (s), 993 (s), 940 (m), 910 (s), 846 (m), 820 (m), 757 (m), 724 (m), 703 (m), 683 (m), 663 (m).

$[\alpha]_D$ = +15 (*c* = 0.005, $CHCl_3$)

**Synthesis of (R)-4-((1R,4Z,8Z,13R,15R)-15-methoxy-5-methyl-3-oxo-2,14-dioxa-bicyclo[11.3.1]heptadeca-4,8-dien-15-yl)-3-(4-methoxybenzyl)thiazolidin-2-one (SI-10):**

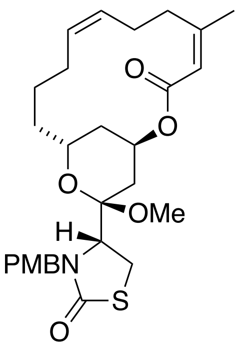


A flame-dried 50 ml *Schlenk* tube was charged with triene **SI-9** (30.0 mg, 55.4 µmol, 1.00 eq.) and dry DCM (degassed, 9.2 ml) were degassed with 3 FPT cycles, backfilled with $N_2$, and Grubbs *Z*-selective catalyst (12.3 mg, 19.4 µmol, 0.30 eq.) was weighed into a flame-dried vial, evacuated, and backfilled with nitrogen (2x) and added as a solution in dry, degassed DCM (1 ml). The reaction mixture was evacuated and backfilled with nitrogen (3x), and then, under slight vacuum, heated to 50 °C. After 24 h, 0.1 ml DMSO was added and the rection was stirred for 24 h open to air. The title compound was obtained as colorless oil (14.0 mg, 29.0 µmol, 45%, 62% brsm).
**Note: the isolated product yield was variable between 33-48%; use of higher catalyst loading only improved the yield up to 55%.*

Reactions with Grubbs 2nd generation catalyst were performed following the above procedure and yielded up to 64% of an (*Z*)/(*E*) product mixture: (*Z*):(*E*) = 40:60.

**$R_f$** ($SiO_2$, Hex:EtOAc = 7:3; CAM) = 0.24

**$^1$H-NMR (400 MHz, $CDCl_3$):** $\delta$ [ppm] = 7.25 – 7.20 (m, 2H), 6.88 – 6.84 (m, 2H), 5.64 (d, *J* = 1.4 Hz, 1H), 5.43-5.40 (m, 1H), 5.27 – 5.23 (m, 1H), 5.07 (d, *J* = 14.2 Hz, 1H), 4.29 (d, *J* = 14.2 Hz, 1H), 4.26 – 4.19 (m, 1H), 3.80 (s, 3H), 3.80 – 3.74 (m, 1H), 3.20 – 3.16 (m, 2H), 3.06 (s, 3H), 2.91 – 2.81 (m, 1H), 2.37 – 2.12 (m, 6H), 1.91 (d (br), *J* = 1.4 Hz, 3H), 1.89 – 1.76 (m, 4H), 1.67 – 1.53 (m, 2H), 1.46 (ddd, *J* = 14.0, 11.7, 2.4 Hz, 1H).

**$^{13}$C-NMR (100 MHz, $CDCl_3$):** $\delta$ [ppm] = 173.3, 166.2, 159.3, 156.0, 130.2, 130.1, 129.1, 129.0, 118.8, 114.1, 102.3, 67.2, 63.7, 59.1, 55.4, 47.7, 47.5, 34.4, 34.2, 31.4, 30.1, 27.1, 25.4, 25.1, 23.9, 21.7.

**HRMS (ESI$^+$, *m/z*):** $[M+NH_4]^+$ for $C_{28}H_{41}O_6N_2S^+$: calcd.: 533.2680, found: 533.2682.

**Synthesis of (R)-4-((1R,4Z,8Z,13R,15R)-15-methoxy-5-methyl-3-oxo-2,14-dioxabicyclo [11.3.1]heptadeca-4,8-dien-15-yl)thiazolidin-2-one (SI-10):**

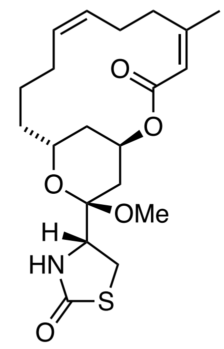


**SI-10** (17.3 mg, 33.5 µmol, 1.00 eq.) was dissolved in MeOH (2.0 ml) and MeCN (0.5 ml). CAN (92.0 mg, 0.17 mmol, 5.00 eq.) was added in one portion at room temperature. The reaction was stirred for 14 h. $NaHCO_3$ (sat. aq., 3 ml) and $H_2O$ (3 ml) were added slowly. The reaction was extracted with DCM (3 x 5 ml), dried over $Na_2SO_4$, filtered and concentrated. Purification by flash column chromatography ($SiO_2$, Hex:EtOAc = 8:2 to 7:3) furnished the title compound as a colorless solid (8.8 mg, 22.2 µmol, 66%).

**$R_f$** ($SiO_2$, hexanes/Ethyl acetate = 7/3) = 0.17

**$^1$H-NMR (400 MHz, $CDCl_3$):** $\delta$ [ppm] = 5.64 (d, $J$ = 1.7 Hz, 1H), 5.51 – 5.33 (m, 2H), 5.25 (p, $J$ = 3.1 Hz, 1H), 4.20 (dt, $J$ = 11.8, 7.3 Hz, 1H), 4.10 (t, $J$ = 8.1 Hz, 1H), 3.41 – 3.21 (m, 2H), 3.19 (s, 3H), 2.88 – 2.80 (m, 1H), 2.32 – 2.10 (m, 6H), 1.92 (s, 3H), 1.95 – 1.87 (m, 1H), 1.88 – 1.32 (m, 6H).

**$^{13}$C-NMR (100 MHz, CDCl$_3$):** $\delta$ [ppm] = 174.7, 166.2, 156.3, 130.0, 129.2, 118.7, 100.5, 67.1, 63.5, 56.7, 48.1, 34.5, 34.1, 30.9, 29.5, 28.1, 27.0, 25.2, 23.8, 21.7.

**Synthesis of (R)-4-((1R,4Z,8Z,13R,15R)-15-hydroxy-5-methyl-3-oxo-2,14-dioxabicyclo [11.3.1]heptadeca-4,8-dien-15-yl)thiazolidin-2-one (8-*Nor*-Latrunculin B):**

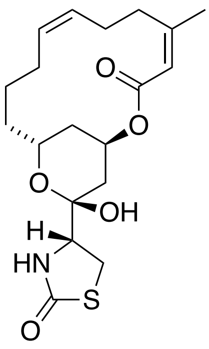


Methylketal **SI-10** (8.80 mg, 22.2 µmol, 1.00 eq.) was dissolved in $H_2O$ (0.6 ml) and AcOH (0.9 ml) and heated to 45 C. After 4 h, the reaction was cooled to room temperature and $NaHCO_3$ (sat. aq., 5 ml) was slowly added. the reaction was extracted with EtOAc (3 x 7 ml), the combined organic layers were washed with brine, dried over $Na_2SO_4$, filtered and concentrated. Purification by flash column chromatography ($SiO_2$, Hex:EtOAc = 9:1 to 6:4) furnished **8-*Nor*-Latrunculin B** as a colorless solid (3.00 mg, 7.90 µmol, 35%).

The *E*/*Z* mixture of methyl ketal **SI-10** was hydrolyzed analogously and gave (*E*/*Z*)-8-*Nor*-Latrunculin B (2.00 mg, 5.2 µmol, 41%).

**R*f*** ($SiO_2$, Hex:EtOAc = 5:5; CAM) = 0.38

**$^{1}$H-NMR (400 MHz, CDCl$_3$):** $\delta$ [ppm] = 5.68 (d, *J* = 1.5 Hz, 1H), 5.63 (s, 1H), 5.46 (p, *J* = 3.2 Hz, 1H), 5.41 – 5.25 (m, 2H), 4.26 (dddd, *J* = 11.6, 9.8, 3.8, 1.7 Hz, 1H), 3.85 – 3.76 (m, 1H), 3.63 (s, 3H), 3.46 (dd, *J* = 11.6, 8.8 Hz, 1H), 3.38 (dd, *J* = 11.7, 6.2 Hz, 1H), 2.54 (td, *J* = 13.5, 12.6, 3.7 Hz, 1H), 2.44 – 2.13 (m, 3H), 2.09 (ddd, *J* = 14.7, 3.1, 1.8 Hz, 1H), 2.03 – 1.95 (m, 1H), 1.95-1.90 (m, 1H), 1.92 (d (br), *J* = 1.4 Hz, 3H), 1.82 – 1.72 (m, 2H), 1.55 – 1.33 (m, 4H).

**$^{13}$C-NMR (100 MHz, $CDCl_3$):** $\delta$ [ppm] = 174.7, 165.5, 155.2, 129.8, 129.5, 118.0, 98.0, 68.6, 62.3, 61.6, 35.6, 35.0, 31.3, 30.9, 28.9, 26.9, 24.4, 23.9, 21.9.

**HRMS (ESI$^+$, *m/z*):** [$(M+H)^+$] for $C_{16}H_{27}NO_5SNa^+$ calcd.: 404.1502, found: 404.1517.

**IR (ATR):** $\tilde{\nu}$ [$cm^{-1}$] = 3310 (w), 2923 (w), 2853 (w), 1678 (s), 1447 (w), 1378 (w), 1347 (w), 1278 (m), 1186 (w), 1186 (w), 1141 (w), 1076 (w), 1054 (w), 1026 (w), 867 (w), 800 (w).

**Synthesis of (R)-4-((1R,4Z,13R,15R)-5-methyl-3-oxo-15-((4-(phenyldiazenyl)benzyl) oxy)-2,14-dioxabicyclo[11.3.1]heptadeca-4,8-dien-15-yl)thiazolidin-2-one (3):**

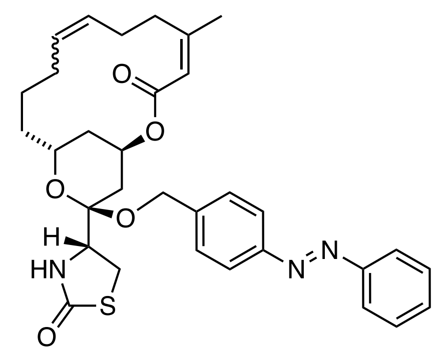


Inspired by a report in the literature[8], a flame-dried 2 ml vial with stir bar was charged with (*E*/*Z*)-**SI-10** (16.0 mg, 40.5 µmol, 1.00 eq.) and azobenzene **SI-25** (85.9 mg, 0.406 mmol, 10.0 eq.) and evacuated for 10 minutes. The starting materials were dissolved in dry DCM (1.3 ml) and $InCl_3$ (3 spatula tips) was added. The reaction instantly turned dark and was left stirring over night. A spatula tip of solid $NaHCO_3$ was added to the reaction and the mixture was filtered over a pad of silica and eluted with DCM and EtOAc. The crude product was purified by HPLC (semi-prep, 40-100% MeCN in $H_2O$ over 8 min, 3 min MeCN, no FA) and gave the title compound as an orange film (2.50 mg, 4.00 µmol, 10%).

[8] Airken, H. R. M.; Furkert, D. P. F.; Hubert, J. G.; Wood, J. M.; Brimble, M. A. *Org. Biomol. Chem.* **2013**, *11*, 5147–5155.

**LCMS** (50-100% MeCN in $H_2O$ over 5 min, 0.1% FA): $t_{ret}$ = 2.6 min (*cis*), 4.1 min (*trans*).

**HPLC** (semi-prep, 40-100% MeCN in $H_2O$ over 8 min, 3 min MeCN, no FA): $t_{ret}$ = 2.6 min.

**$^1$H-NMR (400 MHz, $CDCl_3$):** $\delta$ [ppm] = 7.97 – 7.82 (m, 4H), 7.62 – 7.44 (m, 5H), 5.76 – 5.66 (m, 1H), 5.52 – 5.40 (m, 1H), 5.27 – 5.16 (m, 3H), 4.79 – 4.69 (m, 1H), 4.65 – 4.56 (m, 1H), 4.42 – 4.31 (m, 1H), 4.29 – 4.14 (m, 1H), 3.42 – 3.30 (m, 1H), 3.30 – 3.17 (m, 1H), 2.64 – 2.51 (m, 1H), 2.30 (d, $J$ = 15.1 Hz, 1H), 2.14 – 1.85 (m, 4H), 1.82 (m, 3H) 1.81 –1.32 (m, 8H).

**$^{13}$C-NMR (101 MHz, $CDCl_3$):** δ [ppm] = 174.6, 166.8, 166.5, 157.1, 155.7, 152.7, 152.2, 152.0, 141.1, 141.0, 131.5, 131.3, 131.3, 130.2, 129.4, 129.3, 128.9, 128.7, 127.5, 127.3, 127.1, 123.1, 120.6, 118.1, 101.0, 67.4, 67.1, 65.8, 63.5, 61.7, 61.5, 57.4, 57.3, 35.7, 35.1, 33.5, 33.0, 32.7, 31.4, 31.2, 29.7, 29.8, 29.6, 28.2, 26.3, 24.9, 24.8, 23.9, 23.4, 22.9, 22.2.

NMRs: mixtures of *E*- and *Z*- olefin and *cis* and *trans* isomers.

**HRMS (ESI$^-$, *m/z*):** $[M–H]^-$ for $C_{32}H_{36}N_3O_5S^-$: calcd.: 574.2379, found: 574.2381.

**Synthesis of (*R*)-4-((1*R*,4*Z*,13*R*,15*R*)-5-methyl-3-oxo-15-(4-(phenyldiazenyl)phenethoxy) - 2,14-dioxabicyclo[11.3.1]heptadeca-4,8-dien-15-yl)thiazolidin-2-one (4):**

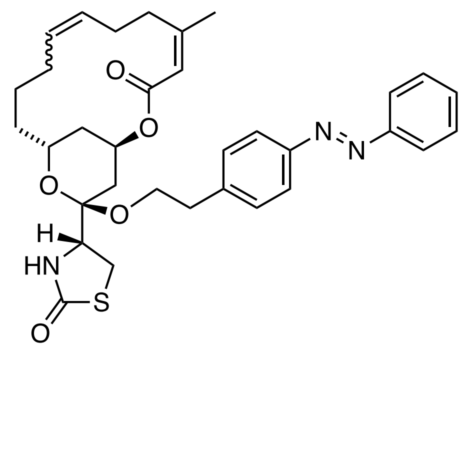

Inspired by a literature report[9], a flame-dried 2 ml vial with stir bar was charged with (*E*/*Z*)-**SI-10** (15.0 mg, 38.0 µmol, 1.00 eq.) and Azobenzene **SI-26** (86.0 mg, 0.38 mmol, 10.0 eq.) and evacuated for 10 minutes. The starting materials were dissolved in dry DCM (1.3 ml) and $InCl_3$ (1 spatula tip) was added. The reaction instantly turned dark and was left stirring for 11 h and another portion of $InCl_3$ (1 spatula tip) was added. A spatula tip of solid $NaHCO_3$ was added to the reaction and the mixture was filtered over a pad of silica (previously rinsed with $NEt_3$ and Hex) and eluted with Hex:EtOAc:$NEt_3$ (4:1:3). The crude product was purified by HPLC (semi-prep, 40-100% MeCN in $H_2O$ over 8 min, 3 min, no FA) and gave the title compound as an orange film (3.50 mg, 5.90 µmol, 16%).

**LCMS** (50-100% MeCN in $H_2O$ over 5 min, 0.1% FA): $t_{ret}$ = 2.9 min (*cis*), 4.3 min (*trans*).

**HPLC** (semi-prep, 40-100% MeCN in $H_2O$ over 8 min, 3 min, no FA): $t_{ret}$ = 3.0 min.

**$^1$H-NMR (600 MHz, $CDCl_3$):** $\delta$ [ppm] = 7.93 – 7.88 (m, 2H), 7.85 (dd, *J* = 8.4, 3.1 Hz, 2H), 7.56 – 7.44 (m, 3H), 7.30 (t, *J* = 8.6 Hz, 2H), 5.69 (dd, *J* = 17.3, 1.6 Hz, 1H), 5.52 – 5.32 (m, 4H), 5.25 – 5.18 (m, 1H), 4.48 – 4.25 (m, 1H), 4.05 – 3.99 (m, 1H), 3.82 – 3.76 (m, 1H), 3.68 – 3.54 (m, 1H), 3.09 – 2.79 (m, 5H), 2.56 – 2.43 (m, 1H), 2.36 – 1.95 (m, 8H), 1.95 – 1.87 (m, 5H), 1.84 – 1.30 (m, 12H).

**$^{13}$C NMR (151 MHz, $CDCl_3$):** $\delta$ [ppm] = 174.6, 174.6, 166.6, 166.2, 157.5, 155.9, 152.8, 151.6, 142.2, 142.2, 132.3, 131.1, 130.0, 129.8, 129.8, 129.4, 129.3, 129.2, 128.9, 123.2, 123.2, 123.0, 121.2, 121.1, 120.3, 120.3, 118.7, 118.1, 100.8, 100.6, 67.4, 67.0, 66.0, 63.7, 61.5, 56.8, 56.8, 36.7, 36.6, 36.1, 35.3, 34.7, 33.5, 33.4, 33.1, 32.4, 31.0, 29.9, 29.6, 29.4, 27.9, 27.2, 25.2, 25.1, 24.1, 23.8, 22.0.

NMRs: mixtures of *E*- and *Z*- olefin and *cis* and *trans* isomers. Rotamers observed.

**HRMS (ESI$^-$, *m/z*):** $[M–H]^-$ for $C_{33}H_{38}N_3O_5S^-$: calcd.: 588.2534, found: 588.2538.

[9] Airken, H. R. M.; Furkert, D. P. F.; Hubert, J. G.; Wood, J. M.; Brimble, M. A. *Org. Biomol. Chem.* **2013**, *11*, 5147–5155.

**Synthesis of (2R,4R,6R)-2-methoxy-2-((R)-2-oxothiazolidin-4-yl)-6-(pent-4-en-1-yl)tetra hydro-2H-pyran-4-yl-11,12-dihydrodibenzo[c,g][1,2]diazocine-3-carboxylate (SI-11):**

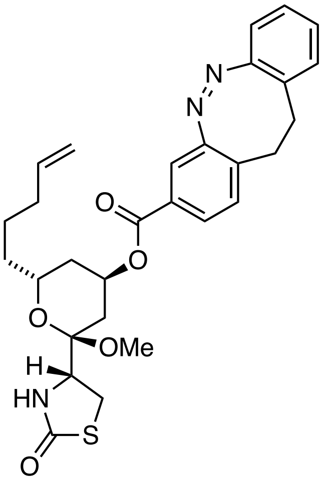


**20** (10.0 mg, 33.2 µmol, 1.00 eq.), diazocine-3-benzoic acid[10] (10.0 mg, 39.8 µmol, 1.20 eq.) and $PPh_3$ (17.4 mg, 66.4 µmol, 2.00 eq.) were dissolved in dry THF (0.66 ml). the reaction mixture was cooled to 0 °C and DEAD (40% in toluene, 32.0 µl, 69.7 µmol, 2.1 eq.) was added. After 10 min., the reaction was allowed to warm to r.t. and stirred for 1 h, concentrated and purified by flash column chromatography ($SiO_2$, Hex:EtOAc = 9:1 to 5:5). The title compound was separated from the inseparable side-product $H_2$DEAD by HPLC (semi-prep, 60-80% MeCN in $H_2O$ over 8 min., 3 min MeCN, no FA) and obtained as an orange film (8.9 mg, 16.6 µmol, 50%).

$\mathbf{R}_f$ ($SiO_2$, Hex:EtOAc = 5:5) = 0.5

**LCMS** (50-100% MeCN in $H_2O$ over 5 min, 0.1% FA): $t_{\text{ret}}$ 3.2 min.

**HPLC:** (semi-prep, 60-80% MeCN in $H_2O$ over 8 min, 3 min MeCN, no FA) $t_{ret}$ = 5.8 min.

**$^1$H-NMR (400 MHz, $CDCl_3$):** $\delta$ [ppm] = 7.69 (d, $J$ = 7.9 Hz, 1H), 7.50 (s, 1H), 7.15 (t, $J$ = 7.6 Hz, 1H), 7.10 – 6.95 (m, 3H), 6.90 – 6.80 (m, 1H), 5.80 (ddt, $J$ = 16.9, 10.1, 6.7 Hz, 1H), 5.47 (s, 1H), 5.35 (s, 1H), 5.08 – 4.93 (m, 2H), 4.15 – 4.07 (m, 1H), 4.00 – 3.90 (m, 1H), 3.37 – 3.18 (m, 6H),

---

[10] Maier M. S., Hüll K., Reynders M., Matsuura B. S., Leippe P., Ko T., Schäffer L., Trauner D. *J. Am. Chem. Soc.* **2019**, *141*, 17295–17304.

3.10 – 2.95 (m, 2H), 2.91 – 2.70 (m, 2H), 2.16 – 2.01 (m, 3H), 2.01 – 1.91 (m, 1H), 1.87 (d, $J$ = 14.4 Hz, 1H), 1.69 – 1.36 (m, 5H).

**$^{13}$C-NMR (101 MHz, CDCl$_3$):** $\delta$ [ppm] = 174.7, 165.0, 155.4, 155.3, 138.5, 133.7, 130.1, 129.8, 129.7, 128.3*, 127.6* 127.5, 127.1, 120.5, 119.0, 115.1, 100.0, 67.1, 66.2*, 66.1*, 56.6, 48.0*, 35.1, 34.4, 33.9, 32.1, 31.6, 29.7*, 29.5*, 28.1, 24.8.

* two peaks or split peaks that correspond to the same carbon. shift due to different conformation of *cis*-diazocine with respect to tetrahydropyran core

**HRMS (ESI$^+$, *m/z*):** [M+Na]$^+$ for $C_{29}H_{33}N_3O_5SNa^+$: calcd.: 558.2033, found: 558.2029.

**IR (ATR):** ṽ [cm$^{-1}$] = 3224 (w), 2937 (w), 1713 (s), 1684 (s), 1457 (w), 1351 (w), 1284 (m), 1256 (m), 1209 (w), 1160 (w), 1120 (s), 1095 (m), 1036 (m), 912 (w), 953 (w).

**Synthesis of (2*R*,4*R*,6*R*)-2-hydroxy-2-((*R*)-2-oxothiazolidin-4-yl)-6-(pent-4-en-1-yl)tetra hydro-2*H*-pyran-4-yl-11,12-dihydrodibenzo[*c*,*g*][1,2]diazocine-3-carboxylate (6):**

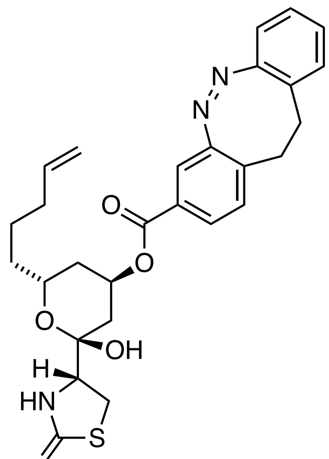


**SI-11** (6.90 mg, 12.9 µmol, 1.00 eq.) was dissolved in THF (1.00 ml) and $H_2O$ (0.45 ml) and AcOH (0.45 ml) was added. the reaction was heated to 45 °C and after 3 h, a second amount of AcOH (0.45 ml) was added and the reaction stirred for 3 h at 45 °C. The reaction was cooled to room temperature and $NaHCO_3$ (sat. aq., 5.00 ml) was added. the reaction was extracted with EtOAc (3 x 5 ml). The combined organic layers were washed with brine, dried over $Na_2SO_4$, filtered and concentrated. The crude product was purified by flash column chromatography ($SiO_2$, Hex:EtOAc = 6:4 to 0:10) and **40** was obtained as an orange film (1.40 mg, 2.70 µmol, 21%).

**R*f*** ($SiO_2$, Hex:EtOAc = 5:5; CAM) = 0.27 (minor), 0.45 (major).

**$^1$H-NMR (400 MHz, $CDCl_3$):** *δ* [ppm] = 7.62 (dd, *J* = 8.0, 1.8 Hz, 1H), 7.46 (d, *J* = 1.8 Hz, 1H), 7.16 (td, *J* = 7.6, 1.5 Hz, 1H), 7.10 (d, *J* = 8.0 Hz, 1H), 7.07 – 6.94 (m, 2H), 6.87 (dd, *J* = 7.8, 1.3 Hz, 1H), 5.77 (ddd, *J* = 16.9, 10.2, 6.1 Hz, 1H), 5.61 (s, 1H), 5.55 (s, 1H), 5.04 – 4.92 (m, 2H), 4.16 – 4.08 (m, 1H), 3.82 (dd, *J* = 8.9, 5.8 Hz, 1H), 3.57 (s, 1H), 3.49 (dd, *J* = 11.3, 9.3 Hz, 1H), 3.42 – 3.34 (m, 1H), 3.09 – 2.97 (m, 2H), 2.92 – 2.74 (m, 2H), 2.09 – 2.00 (m, 4H), 2.00 – 1.90 (m, 1H), 1.61 – 1.36 (m, 5H).

**$^{13}$C-NMR (101 MHz, $CDCl_3$):** *δ* [ppm] = 174.7, 155.6, 155.4, 138.5, 134.6, 130.4, 129.8, 128.6, 128.0, 127.6, 127.5, 127.2, 120.4, 119.1, 115.0, 97.5, 68.8, 65.4, 61.7, 35.0, 34.4, 33.7, 32.1, 31.5, 31.3, 28.9, 24.6.

**HRMS (ESI$^+$, *m/z*):** $[(M+H)–H_2O]^+$ for $C_{28}H_{31}N_3O_4S^+$: calcd.: 405.1952, found: 405.1972.

**IR (ATR):** ṽ [$cm^{-1}$] = 3281 (w), 3066 (w), 2926 (m), 2855 (w), 1680 (s), 1402 (w), 1350 (w), 1285 (s), 1256 (s), 1210 (m), 1158 (m), 1098 (s), 948 (s), 910 (w), 792 (w), 754 (m), 718 (w), 706 (w).

**Synthesis of (2*R*,4*R*,6*R*)-2-methoxy-2-((*R*)-3-(4-methoxybenzyl)-2-oxothiazolidin-4-yl)-6-(pent-4-en-1-yl)tetrahydro-2*H*-pyran-4-yl 4-(phenyldiazenyl)benzoate (SI-12):**

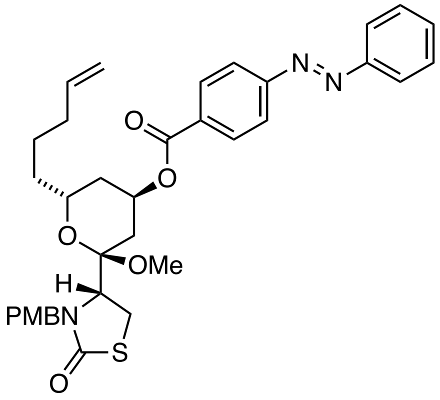


(*S*)-**SI-8** (30.0 mg, 71.2 µmol, 1.00 eq.), *para*-benzoic acid azobenzene (19.3 mg, 85.4 µmol, 1.20 eq.) and $PPh_3$ (37.3 mg, 0.15 mmol, 2.00 eq.) were dissolved in dry THF (1.40 ml). the reaction mixture was cooled to 0 °C and DEAD (40% in toluene, 68.0 µl, 0.15 mmol, 2.1 eq.) was added. After 10 min., the reaction was allowed to warm to r.t. and stirred for 3 h, concentrated and purified by flash column chromatography ($SiO_2$, Hex:EtOAc = 9:1 to 5:5). The title compound was obtained as an orange film (33.3 mg, 52.9 µmol, 74%).

**$^1$H-NMR (400 MHz, $CDCl_3$):** $\delta$ [ppm] = 8.24 – 8.18 (m, 2H), 7.99 – 7.92 (m, 4H), 7.58 – 7.50 (m, 3H), 7.25 – 7.21 (m, 2H), 6.91 – 6.84 (m, 2H), 5.93 – 5.81 (m, 1H), 5.45 ($s_{br}$, 1H), 5.14 (d, *J* = 14.4 Hz, 1H), 5.11 – 4.99 (m, 2H), 4.31 (d, *J* = 14.2 Hz, 1H), 4.13 – 4.04 (m, 1H), 3.86 (t, *J* = 6.0 Hz, 1H), 3.82 (s, 3H), 3.26 ($d_{br}$, *J* = 6.2 Hz, 2H), 3.19 (s, 3H), 2.31 – 2.12 (m, 3H), 2.05 – 1.94 (m, 2H), 1.85 – 1.48 (m, 5H).

**$^{13}$C-NMR (100 MHz, $CDCl_3$):** $\delta$ [ppm] = 173.2, 165.4, 159.3, 155.3, 152.7, 138.4, 132.6, 131.9, 130.8, 129.9, 129.3, 128.9, 123.3, 122.8, 115.2, 114.2, 101.9, 67.3, 66.4, 59.3, 55.4, 47.8, 47.6, 35.7, 34.7, 34.0, 30.4, 25.6, 25.3.
NMR spectra reported for major (*trans*) isomer.

**HRMS (ESI$^+$, *m/z*):** $[M+Na]^+$ for $NaC_{35}H_{39}N_3O_6S^+$: calc.: 652.2452, found: 652.2452.

**IR (ATR):** ṽ [cm⁻¹] = 2938 (w), 2835 (w), 1715 (s), 1672 (s), 1611 (w), 1513 (m), 1444 (w), 1404 (w), 1353 (w), 1276 (s), 1249 (s), 1217 (m), 1196 (m), 1175 (m), 1143 (m), 1120 (m), 1093 (s), 1035 (m), 1012 (m), 913 (w), 864 (w), 847 (w), 779 (m), 718 (w).

**Synthesis of (2*R*,4*R*,6*R*)-2-methoxy-2-((*R*)-2-oxothiazolidin-4-yl)-6-(pent-4-en-1-yl)tetra hydro-2*H*-pyran-4-yl 4-(phenyldiazenyl)benzoate (SI-13 or 5 OMe):**

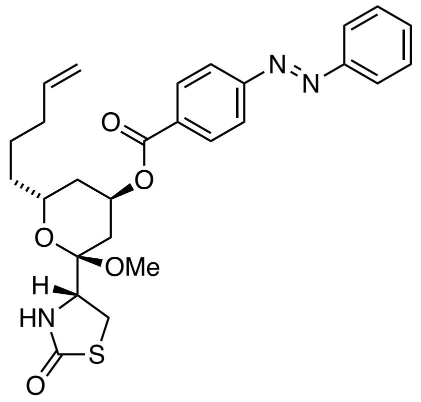


**SI-12** (33.3 mg, 52.9 µmol, 1.00 eq.) was dissolved in MeOH (2.70 ml) and MeCN (0.55 ml) and CAN (145 mg, 0.26 mmol, 5.00 eq.) was added. the reaction was stirred at room temperature over night. $NaHCO_3$ (sat. aq. 5 ml), $H_2O$ (5 ml) and EtOAc (5 ml) were added and the layers separated. The aqueous phase was extracted with EtOAc (3 x 10 ml). the combined organic layers were washed with brine, dried over $Na_2SO_4$, filtered and concentrated. Purification by flash column chromatography ($SiO_2$, Hex:EtOAc = 8:2 to 2:8) and HPLC (semi-prep, 70-90% MeCN in $H_2O$ over 8 min, 3 min MeCN, no FA) gave the title compound as an orange oil (12.8 mg, 25.1 µmol, 48%).

$\mathbf{R}_f$ ($SiO_2$, Hex:EtOAc = 7:3, CAM) = 0.40

**LCMS** (50-100% MeCN in $H_2O$ over 5 min, 0.1% FA): 2.5 min (*cis*) 3.9 min (*trans*).

**HPLC** (semi-prep, 70-90% MeCN in $H_2O$ over 8 min, 3 min MeCN, no FA): $t_{ret}$ = 3.3 min (minor), 6.1 min (major).

**$^1$H-NMR (400 MHz, $CDCl_3$):** $\delta$ = 8.58 (s, 1H), 8.18 – 8.10 (m, 2H), 7.96 – 7.87 (m, 2H), 7.62 (t, $J$ = 7.8 Hz, 1H), 7.57 – 7.48 (m, 3H), 5.88 – 5.72 (m, 1H), 5.57 – 5.45 (m, 2H), 5.07 – 4.93 (m, 2H), 4.16 (t, $J$ = 8.1 Hz, 1H), 4.12 – 4.05 (m, 1H), 3.40 – 3.30 (m, 1H), 3.34 (s. 3H), 2.19 – 1.90 (m, 5H), 1.72 – 1.41 (m, 5H).

**$^{13}$C-NMR (101 MHz, $CDCl_3$):** $\delta$ = 174.7, 165.4, 152.7, 152.6, 138.5, 132.1, 131.8, 131.6, 129.4, 129.3, 127.6, 123.7, 123.1, 115.1, 100.1, 67.1, 66.2, 56.7, 48.1, 35.2, 34.5, 33.9, 29.8, 28.2, 24.9.

**HRMS (ESI$^+$, *m/z*):** $[M+H]^+$ for $C_{26}H_{30}N_3O_5S^+$: calc.: 496.1901, found: 496.1896.

**IR (ATR):** $\tilde{v}$ [$cm^{-1}$] = 3223 (w), 3074 (w), 2939 (w), 1716 (s), 1683 (s), 1437 (w), 1353 (w), 1294 (m), 1272 (m), 1209 (m), 1175 (w), 1153 (m), 1094 (m), 1074 (m), 1035 (m), 911 (w), 819 (w), 769 (m), 723 (w).

**Synthesis of (2*R*,4*R*,6*R*)-2-hydroxy-2-((*R*)-2-oxothiazolidin-4-yl)-6-(pent-4-en-1-yl)tetra hydro-2*H*-pyran-4-yl 4-(phenyldiazenyl)benzoate (5):**

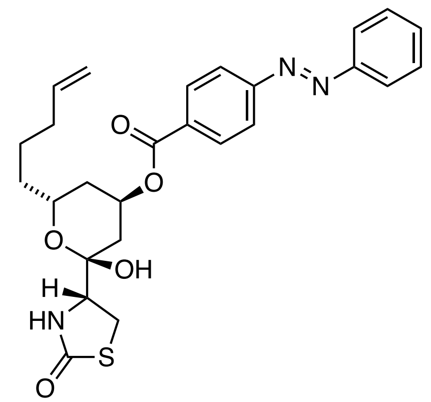


**SI-13** (12.8 mg, 25.1 µmol, 1.00 eq.) was dissolved in THF (0.40 ml) and $H_2O$ (0.40 ml) and AcOH (1.20 ml) was added. the reaction was heated to 45 °C and stirred for 4 h. after cooling to room temperature, $NaHCO_3$ (5 ml) was added, and the reaction was extracted with EtOAc (3 x 10 ml). The combined organic layers were washed with brine, filtered, and concentrated. The crude product was purified by flash column chromatography ($SiO_2$, Hex:EtOAc = 8:2 to 2:8) and HPLC (semi-prep, 50-80% MeCN in $H_2O$ over 8 min, 3 min MeCN, no FA). **SI-13** was obtained as an orange oil (7.10 mg, 14.3 µmol, 57%).

**$R_f$** ($SiO_2$, Hex:EtOAc = 5:5; CAM) = 0.58

**LCMS** (50-100% MeCN in $H_2O$ over 5 min, 0.1% FA): 1.8 min (*cis*) 3.1 min (*trans*).

**HPLC** (semi-prep, 50-80% MeCN in $H_2O$ over 8 min, 3 min MeCN, no FA): 5.5 min (minor), 8.3 min (major).

**$^1$H-NMR (400 MHz, $CDCl_3$):** $\delta$ [ppm] = 8.20 – 8.10 (m, 2H), 8.08 – 7.91 (m, 4H), 7.55 (d, $J$ = 6.3 Hz, 3H), 5.80 (ddt, $J$ = 17.0, 9.5, 6.7 Hz, 1H), 5.70 (s, 1H), 5.69 – 5.61 (m, 1H), 5.08 – 4.93 (m, 2H), 4.29 – 4.17 (m, 1H), 3.86 (dd, $J$ = 8.8, 6.2 Hz, 1H), 3.72 (s, 1H), 3.52 (dd, $J$ = 11.4, 9.2 Hz, 1H), 3.44 (dd, $J$ = 11.8, 6.0 Hz, 1H), 2.24 – 1.96 (m, 5H), 1.70 – 1.34 (m, 5H).

**$^{13}$C-NMR (101 MHz, CDCl$_3$):** $\delta$ [ppm] = 174.8, 164.8, 155.6, 152.6, 138.6, 132.1, 131.4, 130.7, 129.4, 123.4, 123.1, 115.0z, 97.6, 68.9, 65.4, 61.7, 35.1, 34.5, 33.7, 31.4, 28.9, 24.6.

**HRMS (APCI$^+$, *m/z*):** [M+H]$^+$ for $C_{26}H_{30}N_3O_5S^+$: calcd.: 496.1901, found: 496.1896.

**IR (ATR):** ṽ [cm$^{-1}$] = 3323 (w), 3067 (w), 2924 (w), 1680 (s), 1604 (w), 1486 (w), 1444 (w), 1408 (w), 1346 (w), 1276 (s), 1221 (w), 1143 (w), 1096 (m), 1012 (w), 912 (w), 866 (w), 778 (m), 762 (w).

**Synthesis of (2*R*,4*R*,6*R*)-2-methoxy-2-((*R*)-3-(4-methoxybenzyl)-2-oxothiazolidin-4-yl)-6-(pent-4-en-1-yl)tetrahydro-2*H*-pyran-4-yl 3-(phenyldiazenyl)benzoate (SI-14):**

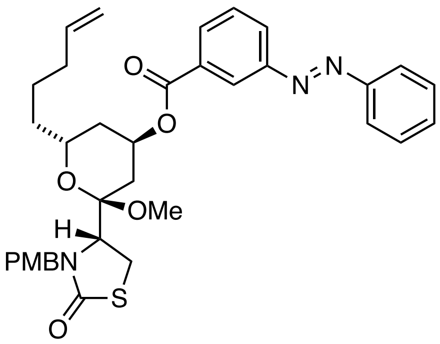


A flame dried vial with stir bar was charged with (*S*)-**SI-8** (30.0 mg, 71.2 µmol, 1.00 eq.), Azobenzene **SI-28**(193 mg, 85.4 µmol, 1.20 eq.) and $PPh_3$ (37.3 mg, 0.14 mmol, 2.00 eq.). The reaction was evacuated/ backfilled with nitrogen and dissolved in dry THF (1.42 ml). the reaction was cooled to 0 °C and DEAD (40% in toluene, 68.0 µl, 0.15 mmol, 2.10 eq.) was added dropwise. The reaction was stirred for 10 minutes at 0 °C and warmed to room temperature. After stirring for 4 h at r.t., the reaction mixture was concentrated and subjected to flash column chromatography ($SiO_2$, Hex:EtOAc = 9:1 to 5:5). The title compound was obtained as orange oil (39.8 mg, 63.2 µmol, 89%).

**R*f*** ($SiO_2$, Hex:EtOAc = 5:5) = 0.7

**$^{1}$H-NMR (400 MHz, $CDCl_3$):** $\delta$ [ppm] = 8.59 (d, $J$ = 2.0 Hz, 1H), 8.19 – 8.10 (m, 2H), 7.96 – 7.90 (m, 2H), 7.66 – 7.58 (m, 1H), 7.56 – 7.50 (m, 3H), 7.25 – 7.21 (m, 2H), 6.91 – 6.85 (m, 2H), 5.87 (dtd, $J$ = 17.9, 9.4, 8.5, 6.3 Hz, 1H), 5.48 (s, 1H), 5.14 (d, $J$ = 14.5 Hz, 1H), 5.11 – 4.97 (m, 2H), 4.31 (d, $J$ = 14.5 Hz, 1H), 4.15 – 4.07 (m, 1H), 3.87 (dd, $J$ = 7.1, 4.9 Hz, 1H), 3.82 (s, 3H), 3.28 – 3.25 (m, 2H), 3.23 (s, 3H), 2.33 – 2.12 (m, 3H), 2.07 – 1.94 (m, 2H), 1.83 – 1.49 (m, 5H).

**$^{13}$C-NMR (101 MHz, $CDCl_3$):** $\delta$ [ppm] = 173.2, 165.4, 159.3, 152.7, 152.6, 138.4, 132.2, 131.8, 131.6, 130.0, 129.3, 128.9, 127.7, 123.7, 123.1, 120.8, 115.2, 114.2 101.9, 67.2, 66.4, 59.3, 55.4, 47.8, 47.6, 35.7, 34.72, 34.0, 30.5, 25.6, 25.2.

**HRMS (**APCI$^+$, *m/z*): [M+H]$^+$ for $C_{35}H_{40}N_3O_6S^+$: calc.: 630.2632, found: 630.2609
[(M+Na)$^+$] for $NaC_{35}H_{39}N_3O_6S^+$: calc.: 652.2452, found: 652.2493.

**IR (ATR):** $\tilde{\nu}$ [cm$^{-1}$] = 2937 (w), 2835 (w), 1715 (m). 1667 (s), 1611 (w), 1586 (w), 1512 (m), 1441 (w), 1401 (w), 1354 (w), 1294 (m) 1269 (s), 1247 (s), 1210 (s), 1174 (s), 1152 (m), 1125 (m), 1091 (s), 1072 (s), 1032 (s), 999 (m), 913 (m), 845 (m), 818 (m), 769 (s), 756 (m), 734 (s).

**Synthesis of (2*R*,4*R*,6*R*)-2-methoxy-2-((*R*)-2-oxothiazolidin-4-yl)-6-(pent-4-en-1-yl)tetrahydro-2*H*-pyran-4-yl 3-(phenyldiazenyl)benzoate (SI-15 or 7 OMe):**

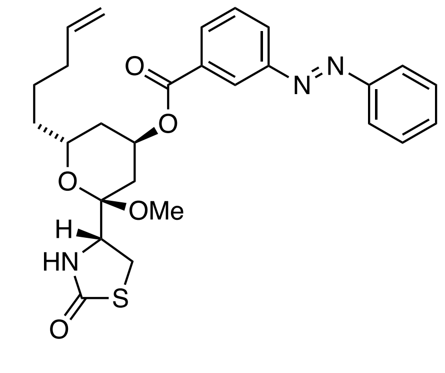


**SI-14** (32.4 mg, 51.4 µmol, 1.00 eq.) was dissolved in MeOH (2.6 ml) and MeCN (0.5 ml) and CAN (141 mg, 0.26 mmol, 5.00 eq.) was added at room temperature. The reaction was stirred over night and $NaHCO_3$ (sat. aq., 5 ml), $H_2O$ (5 ml) and EtOAc (5 ml) were added. The layers were separated, and the aqueous layer was extracted with EtOAc (3 x 5 ml). The combined organic layers were washed with brine, dried over $Na_2SO_4$, filtered and concentrated. The crude product was purified by flash column chromatography ($SiO_2$, Hex:EtOAc = 8:2 to 5:5) and HPLC (semi-prep, 70-90% MeCN in $H_2O$ over 8 min, 3 min MeCN, no FA) and furnished **SI-15** as an orange film (8.90 mg, 17.4 µmol, 34%).

**$R_f$** ($SiO_2$, Hex:EtOAc = 7:3; CAM) = 0.43

**LCMS** (50-100% MeCN in $H_2O$ over 5 min, 0.1% FA): 2.5 min (*cis*), 3.9 min (*trans*).

**HPLC** (semi-prep, 70-90% MeCN in $H_2O$ over 8 min, 3 min MeCN, no FA): 3.3 min (minor), 6,1 min (major).

**$^1$H-NMR (400 MHz, $CDCl_3$):** $\delta$ [ppm] = 8.20 (d, $J$ = 8.1 Hz, 2H), 8.00 – 7.91 (m, 4H), 7.58 – 7.49 (m, 3H), 5.81 (ddt, $J$ = 16.9, 9.6, 6.7 Hz, 1H), 5.50 (s, 1H), 5.45 ($s_{br}$, 1H), 5.08 – 4.94 (m, 2H), 4.15 (t, $J$ = 8.1 Hz, 1H), 4.11 – 4.01 (m, 1H), 3.41 – 3.27 (m, 2H), 3.30 (s, 3H), 2.20 – 1.86 (m, 5H), 1.72 – 1.40 (m, 5H).

**$^{13}$C-NMR (101 MHz, $CDCl_3$)**: $\delta$ [ppm] = 174.7, 165.4, 155.3, 152.7, 138.5, 132.5, 131.9, 130.8, 129.4, 123.3, 122.8, 122.8, 115.1, 100.1, 67.2, 66.2, 56.7, 48.1, 35.2, 34.5, 33.9, 29.7, 28.2, 24.9.

**HRMS** (APCI$^+$, *m/z*): $[M+H]^+$ for $C_{27}H_{32}N_3O_5S^+$: calc.: 510.2057, found: 510.2036

**Synthesis of (2R,4R,6R)-2-hydroxy-2-((R)-2-oxothiazolidin-4-yl)-6-(pent-4-en-1-yl)tetra hydro-2H-pyran-4-yl 3-((E)-phenyldiazenyl)benzoate (7):**

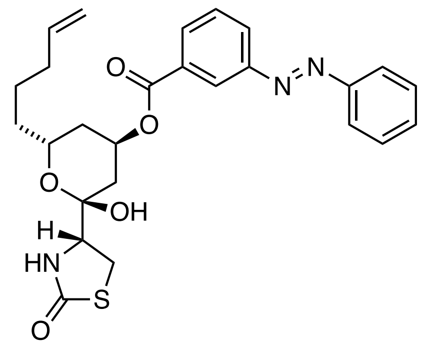

**SI-15** (9.00 mg, 17.7 µmol, 1.00 eq.) was dissolved in THF (0.30 ml) and water (0.30 ml). Next, AcOH (0.30 ml) was added and the reaction was heated to 45 °C for 4 h. After cooling to room temperature, $NaHCO_3$ (sat. aq., 5 ml) and EtOAc (5 ml) were added and separated. The aqueous layer was extracted with EtOAc (3 x 5 ml) and the combined organic layers were washed with brine, dried over $Na_2SO_4$, filtered and concentrated. The crude product was purified by flash column chromatography ($SiO_2$, Hex:EtOAc = 8:2 to 2:8) and HPLC (semi-prep, 50-80% MeCN in $H_2O$ over 8 min, 3 min MeCN, no FA) and gave **7** as an orange film (4.70 mg, 9.50 µmol, 54%).

**$R_f$** ($SiO_2$, Hex:EtOAc = 5:5; CAM) = 0.61

**LCMS** (50-100% MeCN in $H_2O$ over 5 min, 0.1% FA): $t_{ret}$ = 1.8 min (cis), 3.1 min (trans).

**HPLC** (semi-prep, 50-80% MeCN in $H_2O$ over 8 min, 3 min MeCN, no FA): $t_{ret}$ = 5.4 min (minor), 8.1 min (major).

**$^1$H-NMR (400 MHz, $CDCl_3$):** $\delta$ [ppm] = 8.52 (d, $J$ = 1.9 Hz, 1H), 8.18 – 8.08 (m, 2H), 7.95 (dd, $J$ = 7.4, 2.5 Hz, 2H), 7.67 – 7.60 (m, 1H), 7.60 – 7.49 (m, 3H), 5.80 (ddt, $J$ = 16.9, 8.9, 6.6 Hz, 1H), 5.73 – 5.63 (m, 2H), 4.99 (dd, $J$ = 21.0, 13.7 Hz, 2H), 4.29 – 4.19 (m, 1H), 3.86 (dd, $J$ = 8.8, 6.2 Hz, 1H), 3.72 (s, 1H), 3.56 – 3.39 (m, 2H), 2.25 – 1.96 (m, 5H), 1.71 – 1.41 (m, 5H).

**$^{13}$C-NMR (101 MHz, $CDCl_3$):** $\delta$ [ppm] = 174.7, 164.8, 152.9, 152.5, 138.6, 131.8, 131.5, 131.0, 129.7, 129.3, 127.1, 124.5, 123.3, 115.0, 97.6, 69.0, 65.4, 61.7, 35.1, 34.5, 33.7, 31.4, 28.9, 24.6.

**HRMS (ESI$^+$, *m/z*):** $[M+Na]^+$ for $C_{26}H_{29}N_3O_5SNa^+$: calcd.: 518.1720, found: 518.1743.

**IR (ATR):** $\tilde{\nu}$ [$cm^{-1}$] = 3317 (w), 2922 (w), 2855 (w), 1716 (m), 1679 (s), 1439 (w), 1349 (w), 1295 (m), 1271 (m), 1210 (m), 1176 (w), 1153 (m), 1100 (m), 1070 (m), 1021 (w), 913 (w), 814 (w), 806 (w), 770 (m), 759 (w), 716 (w).

**Synthesis of (2*R*,4*R*,6*R*)-2-methoxy-2-((*R*)-3-(4-methoxybenzyl)-2-oxothiazolidin-4-yl)-6-(pent-4-en-1-yl)tetrahydro-2*H*-pyran-4-yl 2-(phenyldiazenyl)benzoate (SI-16):**

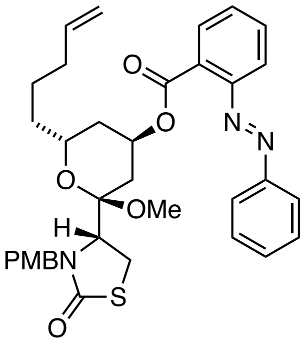


A flame-dried vial was equipped with a stir bar and charged with (*S*)-**SI-8** (30.0 mg, 71.2 µmol, 1.00 eq.), azobenzene **SI-27** (142 mg, 0.14 mmol, 2.00 eq.) and $PPh_3$ (93.3 mg, 0.36 mmol, 5.00 eq.). The reaction mixture was evacuated and backfilled with nitrogen, dissolved in dry toluene (1.4 ml) and cooled to 0 °C. DEAD (40% in toluene, 0.20 ml, 0.50 mmol, 7.00 eq.) was added dropwise and the reaction was stirred at 0 °C for 10 minutes, following 14 h at room temperature. The crude reaction mixture was loaded on $SiO_2$ (3.0 g), dried and subjected to flash column chromatography ($SiO_2$, Hex:EtOAc = 9.5:0.5 to 6:4). The desired product was inseparable from the $H_2$DEAD side product and subjected to the subsequent transformation without further purification.

**Synthesis of (2*R*,4*R*,6*R*)-2-methoxy-2-((*R*)-2-oxothiazolidin-4-yl)-6-(pent-4-en-1-yl)tetra hydro-2*H*-pyran-4-yl 2-(phenyldiazenyl)benzoate (SI-17):**

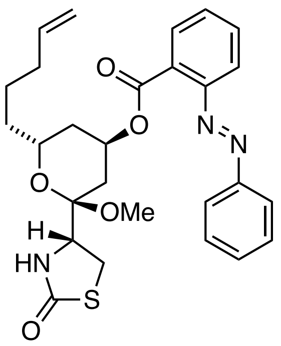


Crude **SI-16** was dissolved in MeOH (3.6 ml) and MeCN (0.7 ml) and CAN (195 mg, 0.36 mmol, 5.00 eq.) was added. The reaction was stirred over night at room temperature and $NaHCO_3$ (sat. aq., 10 ml) and water (10 ml) were added. The reaction was extracted with DCM (3 x 30 ml). the combined organic layers were washed with brine, dried over $Na_2SO_4$, filtered and concentrated. The crude product was purified by flash column chromatography ($SiO_2$, Hex:EtOAc = 8:2 to 5:5) and HPLC (semi-prep, 30 –80% MeCN in $H_2O$ over 8 min, 3 min MeCN, no FA). The title compound was obtained as an orange film (11.3 mg, 22.2 µmol, 31% over two steps).

$\mathbf{R}_f$ ($SiO_2$, Hex:EtOAc = 5/5) = 0.5

**LCMS** (50-100% MeCN in $H_2O$ over 5 min, 0.1% FA): tRet = 2.5 min. (*cis*), 3.5 min. (*trans*).

**HPLC** (semi-prep, 40-80% MeCN in $H_2O$ over 8 min, 3 min MeCN, no FA): $t_{ret}$ = 9.19 min (major).

**$^1$H-NMR (400 MHz, $CDCl_3$):** $\delta$ [ppm] = 7.94 – 7.87 (m, 3H), 7.64 – 7.57 (m, 1H), 7.55 – 7.47 (m, 5H), 5.75 (ddt, $J$ = 16.9, 10.1, 6.6 Hz, 1H), 5.45 (s, 1H), 5.44 – 5.40 (m, 1H), 5.03 – 4.92 (m, 2H), 4.08 – 4.00 (m, 1H), 3.85 – 3.74 (m, 1H), 3.28 – 3.07 (m, 2H), 3.11 (s, 3H), 2.15 – 1.79 (m, 5H), 1.56 – 1.10 (m, 5H).

**$^{13}$C-NMR (101 MHz, $CDCl_3$):** $\delta$ [ppm] = 174.7, 166.5, 152.8, 152.7, 138.5, 132.3, 131.6, 130.2, 129.7, 129.3, 128.9, 123.4, 118.5, 115.0, 99.9, 67.5, 66.0, 56.6, 47.8, 35.1, 34.5, 33.9, 29.5, 28.0, 24.8.

**HRMS (ESI$^+$, *m/z*):** $[M+Na]^+$ for $NaC_{27}H_{31}N_3O_5S^+$ calcd.: 532.1877, found: 532.1880.

**Synthesis of (2R,4R,6R)-2-hydroxy-2-((R)-2-oxothiazolidin-4-yl)-6-(pent-4-en-1-yl)tetrahydro-2H-pyran-4-yl 2-(phenyldiazenyl)benzoate (8):**

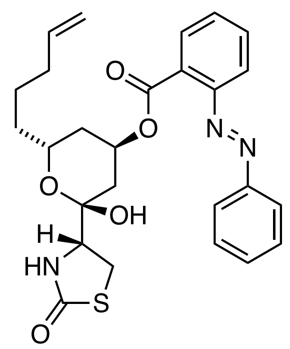


**SI-17** (10.0 mg, 19.6 µmol, 1.00 eq.) was dissolved in THF (0.6 ml), $H_2O$ (0.6 ml) and AcOH (0.9 ml) and stirred at 45 °C for 2 h. Then, AcOH (0.9 ml) was again added, and the reaction stopped after 2 h by pouring it into $NaHCO_3$ (sat. aq., 20 ml). the reaction was extracted with EtOAc (3 x 20 ml). The combined organic layers were washed with brine, dried over $Na_2SO_4$, filtered and concentrated. The crude product was purified twice by HPLC (semi-prep, 40 – 80% MeCN in $H_2O$ over 8 min, 3 min MeCN, no FA) and **8** was obtained as an orange film (1.00 mg, 2.00 µmol, 10%).
Note: during the reaction, decomposition was observed. The reaction was thus stopped to obtain a small quantity of the final product for biological evaluation by cell proliferation assay.

**R*f*** ($SiO_2$, Hex:EtOAc = 5:5; CAM) = 0.54 (major), 0.44 (minor).

**LCMS** (50-100% MeCN in $H_2O$ over 5 min, 0.1% FA): $t_{Ret}$ = 1.7 min (*cis*), 2.4 min (*trans*).

**HPLC** (semi-prep, 40 – 80% MeCN in $H_2O$ over 8 min, 3 min MeCN, no FA) $t_{ret}$ = 6.6 min (minor), 7.9 min (major).

**$^1$H-NMR (400 MHz, $CDCl_3$):** $\delta$ [ppm] = 7.96 – 7.83 (m, 3H), 7.74 – 7.45 (m, 6H), 5.77 – 5.59 (m, 3H), 5.51 (s, 1H), 5.08 – 4.80 (m, 3H), 4.39 ($s_{br}$, 1H), 3.79 – 3.70 (m, 1H), 3.70 – 3.63 (m, 1H), 3.56 – 3.36 (m, 1H), 3.25 (dd, $J$ = 11.7, 9.0 Hz, 1H), 2.95 (dd, $J$ = 11.7, 5.6 Hz, 1H), 2.18 – 1.05 (m, 7H).

**HRMS (ESI$^+$, *m/z*):** $[M+Na]^+$ for $C_{26}H_{29}N_3O_5S^+$: calcd.: 518.1720, found: 518.1742.

**Synthesis of (R)-4-((2R,4R,6R)-2-methoxy-6-(pent-4-en-1-yl)-4-(4-(phenyldiazenyl) phenoxy)tetrahydro-2H-pyran-2-yl)thiazolidin-2-one (SI-18):**

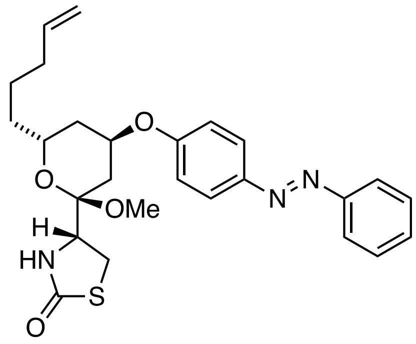


**20** (14.8 mg, 49.1 µmol, 1.00 eq.), $PPh_3$ (32.2 mg, 123 µmol, 2.50 eq.) and 4-hydroxy azobenzene (19.5 mg, 98.2 µmol, 2.00 eq.) were dissolved in dry THF (0.90 ml) and dry toluene (0.30 ml) and cooled to 0 °C using an ice-bath. To the orange reaction mixture, DEAD (40% in toluene, 56.0 µl, 123 µmol, 2.50 eq.) was added and dropwise and the reaction turned dark red. The reaction was stirred at 0 °C for 10 min. before it was warmed to room temperature and stirred over night. The reaction mixture was loaded on isolute, concentrated *in vaccuo* and submitted to purification by flash column chromatography ($SiO_2$, Hex:EtOAc 9.5:0.5 to 5:5, slow gradient). The title compound was repurified by HPLC (semi-prep, 50 – 90% MeCN in $H_2O$ over 8 min, 3 min MeCN, no FA) yielding the title compound as an orange film (13.0 mg, 27.0 µmol, 55%).

**$R_f$** ($SiO_2$, Hex:EtOAc = 7:3) = 0.44 (major), 0.25 (minor).

**LCMS** (50-100% MeCN in H2O over 5 min, 0.1% FA): $t_{Ret}$ = 2.4 min (*cis*), 3.8 min. (*trans*)

**HPLC** (semi-prep, 50 – 90% MeCN in $H_2O$ over 8 min, 3 min MeCN, no FA): 5.8 min (minor), 8.4 min (major).

**$^1$H-NMR (600 MHz, $CDCl_3$):** $\delta$ = 7.91 (d, $J$ = 8.9 Hz, 2H), 7.89 – 7.85 (m, 2H), 7.53 – 7.48 (m, 2H), 7.46 – 7.42 (m, 1H), 7.01 (d, $J$ = 8.9 Hz, 2H), 5.80 (ddt, $J$ = 16.9, 10.1, 6.7 Hz, 1H), 5.52 (s, 1H), 5.05 – 4.94 (m, 2H), 4.87 (s, 1H), 4.15 – 4.11 (m, 1H), 4.10 – 4.04 (m, 1H), 3.38 – 3.25 (m, 2H), 3.27 (s, 3H), 2.14 – 2.06 (m, 3H), 2.03 – 1.97 (m, 2H), 1.65 – 1.41 (m, 5H).

**$^{13}$C-NMR (151 MHz, $CDCl_3$):** $\delta$ = 174.8, 159.9, 152.9, 147.3, 138.5, 130.6, 129.2, 124.9, 122.7, 116.4, 115.1, 100.0, 69.3, 65.6, 56.7, 48.0, 35.1, 34.2, 33.9, 28.9, 28.2, 24.9.

NMRs are reported for major isomer (*trans*).

**HRMS (ESI$^+$, *m/z*):** $[M+Na]^+$ for $NaC_{26}H_{31}N_3O_4S^+$: calcd.: 504.1927, found: 504.1935.

**IR (ATR):** $\tilde{\nu}$ [cm$^{-1}$] = 3069 (w), 2938 (w), 2361 (w), 2343 (w), 1683 (s), 1599 (m), 1580 (w), 1497 (m), 1442 (w), 1417 (w), 1299 (w), 1248 (m), 1140 (w), 1096 (m), 1036 (w), 913 (w), 839 (w), 769 (w), 721 (w).

**Synthesis of (*R*)-4-((2*R*,4*R*,6*R*)-2-hydroxy-6-(pent-4-en-1-yl)-4-(4-(phenyldiazenyl) phenoxy)tetrahydro-2*H*-pyran-2-yl)thiazolidin-2-one (9):**

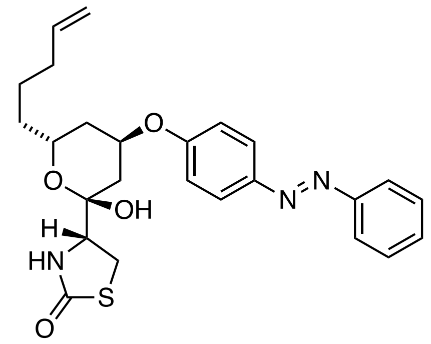

**SI-18** (12.8 mg, 26.6 µmol, 1.00 eq.) was dissolved in THF (0.40 ml) and water (0.40 ml) and AcOH (0.40 ml) was added. The reaction mixture was heated to 50 °C for 6.5 h. The reaction was poured into sat. aq. $NaHCO_3$ (30 ml) and Ethyl acetate (30 ml). The layers were separated and the aqueous phase was extracted with Ethyl acetate (2 x 20 ml). The combined organic layers were washed with brine, dried over $Na_2SO_4$, filtered and concentrated. The crude product was filtered through a plug of silica, eluting with EtOAc and then purified by HPLC (semi-prep, 40 – 80% MeCN in $H_2O$ over 8 min, 3 min MeCN, no FA). The title compound was obtained in 48% yield (6.0 mg, 12.8 µmol) as an orange film.

**$R_f$** ($SiO_2$, Hex:EtOAc = 4:6; CAM) = 0.59

**LCMS** (50-100% MeCN in $H_2O$ over 5 min, 0.1% FA): $t_{ret}$ (minor) = 1.7 min, $t_{ret}$ (major) = 3.1 min.

**HPLC** (semi-prep; 40 – 80% MeCN in $H_2O$ over 8 min., 3 min MeCN, no FA): $t_{ret}$ (minor) = 5.4 min, $t_{ret}$ (major) = 9.2 min.

**$^1$H-NMR (400 MHz, $CDCl_3$):** $\delta$ [ppm] = 7.95 (d, $J$ = 8.9 Hz, 2H), 7.89 (dd, $J$ = 8.4, 1.5 Hz, 2H), 7.56 – 7.43 (m, 3H), 7.05 (d, $J$ = 8.9 Hz, 2H), 5.83 – 5.71 (m, 1H), 5.72 (s, 1H), 5.09 – 4.90 (m, 4H), 4.20 – 4.11 (m, 1H), 3.88 (ddd, $J$ = 9.1, 5.5, 1.1 Hz, 1H), 3.55 (dd, $J$ = 11.7, 9.0 Hz, 1H), 3.44 (dd, $J$ = 11.8, 5.6 Hz, 1H), 2.25 – 2.18 (m, 1H), 2.12 – 1.99 (m, 4H), 1.64 – 1.35 (m, 5H).

**$^{13}$C-NMR (101 MHz, $CDCl_3$):** $\delta$ [ppm] = 174.9, 158.3, 152.7, 148.1, 138.6, 130.9, 129.2, 125.1, 122.8, 116.2, 115.0, 97.8, 72.5, 64.7, 61.0, 35.0, 33.7, 33.5, 31.7, 29.0, 24.6.

**HRMS (ESI$^+$, *m/z*):** $[M+H]^+$ for $C_{25}H_{30}N_3O_4S^+$: calcd.: 468.1952, found: 468.1960.

**Synthesis of (*R*)-4-((2*R*,4*R*,6*R*)-2-hydroxy-6-(pent-4-en-1-yl)-4-(2-((*E*)- phenyldiazenyl)-phenoxy)- tetrahydro-2*H*-pyran-2-yl)thiazolidin-2-one (10):**

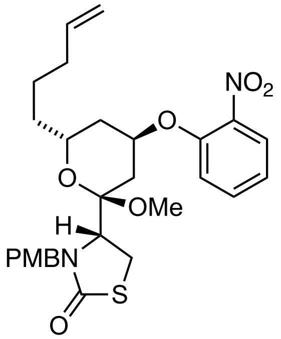


**SI-19**

(*S*)-**SI-8** (20.0 mg, 0.05 mmol, 1.00 eq.) was dissolved in DMF (1.00 ml) and cooled to 0 °C. NaH (2.30 mg, 0.10 mmol, 2.00 eq.) was added in one portion. The reaction was stirred for 30 min under nitrogen. 1-fluoro-2-nitrobenzene (13.4 mg, 0.10 mmol, 2.00 eq.) and the mixture was stirred at 50 °C for 3 h. $NH_4Cl$ (sat. aq., 2.00 ml), $H_2O$ (10 ml) and EtOAc (15 ml) were added, and the layers separated. The aqueous phase was extracted with EtOAc (2 x 10 ml). the combined organic layers were washed with brine, dried over $Na_2SO_4$, filtered and concentrated. Purification by flash column chromatography ($SiO_2$, Hex:EtOAc 9:1 to 3:1) furnished the $S_NAr$ product as a colorless solid (20.0 mg, 0.04 mmol, 80%).

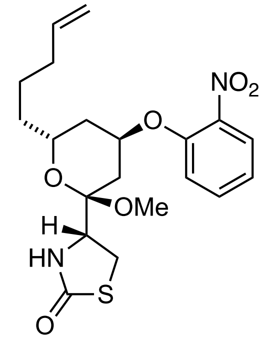


**SI-20**

**SI-19** (20.0 mg, 0.04 mmol, 1.00 eq.) was dissolved in MeOH (8.00 mL) and MeCN (2.00 ml) and CAN (0.16 g, 0.30 mmol, 8.00 eq.) was added in one portion. The reaction was stirred for 16 h under nitrogen. $NaHCO_3$ (sat. aq., 10 ml), $H_2O$ (10 mL) and EtOAc (20 ml) were added, and the layers separated. The aqueous phase was extracted with EtOAc (2 x 20 mL). the combined organic layers were washed with brine, dried over $Na_2SO_4$, filtered and concentrated. Purification by flash column chromatography ($SiO_2$, Hex:EtOAc 5:1 to 1:1) furnished **SI-20** compound as a colorless solid (9.3 mg, 0.02 mmol, 60%).

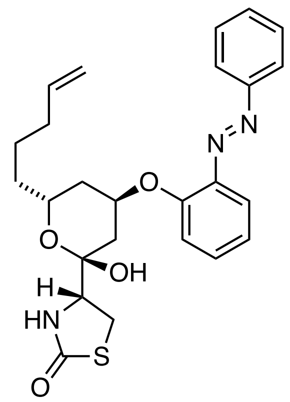

**SI-20** (9.3 mg, 0.02 mmol, 1.00 eq.) was dissolved in MeOH (3.00 ml) and THF (0.50 ml) and Lindlar catalyst (10.0 mg) was added in one portion. The reaction was stirred for 45 min under hydrogen. Then the catalyst was filtered out and the filtrate was concentrated under reduced pressure. Nitrosobenzene (5.1 mg, 0.05 mmol, 2.00 eq.) and AcOH (1.00 ml) was added, and the mixture was stirred at room temperature overnight. Water (0.60 ml) was added, and the reaction was heated at 60 °C for 3-4 h. The pH was made slightly basic (pH 7-8) by adding 5% NaHCO3 and EtOAc (10 ml) were added. The aqueous phase was extracted with EtOAc (2 x 50 mL). the combined organic layers were washed with brine, dried over $Na_2SO_4$, filtered and concentrated. The crude product was purified by HPLC and the title compound was obtained as an orange film (0.43 mg, 0.90 µmol, 5%*.)

**not optimized, yield over three steps: reduction of the nitro group, the Mills reaction and the hydrolysis of methyl ketal under acidic conditions.*

**$R_f$** ($SiO_2$, Hex:EtOAc = 4:6; CAM) = 0.59

**$^1$H NMR (400 MHz, CDCl3):** $\delta$ [ppm] = 7.92 (d, $J$ = 7.3 Hz, 2H), 7.67 (dd, $J$ = 8.1, 1.7 Hz, 1H), 7.55 - 7.46 (m, 3H), 7.41 (ddd, $J$ = 8.3, 7.3, 1.8 Hz, 1H), 7.19 – 7.13 (m, 1H), 7.13 – 7.03 (m, 1H), 5.92 – 5.57 (m, 2H), 5.16 – 4.93 (m, 2H), 4.89 (dt, $J$ = 11.1, 6.1 Hz, 1H), 3.98 – 3.87 (m, 1H), 3.79 (t, $J$ = 7.4 Hz, 1H), 3.45 (dd, $J$ = 7.3, 4.8 Hz, 1H), 2.46 – 2.37 (m, 1H), 2.30 – 2.21 (m, 1H), 2.05 (d, $J$ = 5.8 Hz, 2H), 1.58 – 1.44 (m, 6H).

**$^{13}$C-NMR (101 MHz, $CDCl_3$):** $\delta$ [ppm] = 174.6, 155.1, 153.0, 144.2, 138.4, 132.20, 131.0, 129.1, 123.1, 122.5, 119.0, 117.4, 114.9, 99.1, 74.1, 69.8, 62.5, 37.5, 35.0, 34.0, 33.6, 29.7, 28.8, 24.6.

**HRMS (ESI+, *m/z*):** [(M+H)+] for $C_{25}H_{30}N_3O_4S^+$: calcd.: 468.1952, found: 468.1965.

**IR (ATR):** ṽ [cm⁻¹] = 3853 (w), 3744 (w), 2925 (w), 1750 (w), 1717 (w), 1540 (w), 1506 (m), 1521 (m), 1363 (s), 1229 (w), 1217 (w), 1000 (w), 720 (w).

**Synthesis of (*R*)-4-((2*R*,4*R*,6*R*)-2-methoxy-6-(pent-4-en-1-yl)-4-(4-((3,4,5-trimethoxy phenyl)diazenyl)phenoxy)tetrahydro-2*H*-pyran-2-yl)thiazolidin-2-one (SI-21):**

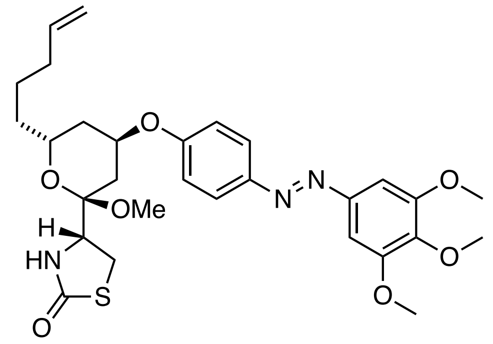


**20** (15.0 mg, 49.8 µmol, 1.00 eq.), $PPh_3$ (32.6 mg, 124 µmol, 2.50 eq.) and **SI-28** 54.2 mg, 99.6 µmol, 2.00 eq.) were dissolved in dry THF (0.90 ml) and dry toluene (0.30 ml) and cooled to 0 °C using an ice-bath. To the orange reaction micture, DEAD (40% in toluene, 57.0 µl, 124 µmol, 2.50 eq.) was added and dropwise and the reaction turned dark red. The reaction was stirred at 0 °C for 30 min. before it was warmed to room temperature and stirred over night. The reaction mixture was loaded on isolute, concentrated *in vaccuo* and submitted to flash column purification ($SiO_2$, Hex:EtOAc 9.5:0.5 to 5:5). The purified product contained $H_2$DEAD, which could be removed by HPLC purification (semi-prep, 50 – 90% MeCN in $H_2O$ over 8 min, 3 min MeCN, no FA). The title compound was obtained in 27% yield (7.60 mg, 13.3 µmol) as an orange film.

$\mathbf{R}_f$ ($SiO_2$, Hex:EtOAc = 7:3) = 0.24

**LCMS** (50-100% MeCN in $H_2O$, 0.1% FA) $t_{ret}$ = 2.1 min (*cis*), $t_{ret}$ = 3.3 min (*trans*).

**HPLC** (semi-prep, 50 – 90% MeCN in $H_2O$ over 8 min, 3 min MeCN, no FA): $t_{ret}$ = 7.4 min (major).

**$^1$H-NMR (400 MHz, CDCl$_3$):** $\delta$ [ppm] = 7.89 (d, $J$ = 8.9 Hz, 2H), 7.21 (s, 2H), 7.01 (d, $J$ = 9.0 Hz, 2H), 5.80 (ddt, $J$ = 17.0, 10.2, 6.7 Hz, 1H), 5.49 (s, 1H), 5.07 – 4.92 (m, 2H), 4.86 (s$_{br}$, 1H), 4.18 – 4.03 (m, 2H), 3.96 (s, 6H), 3.93 (s, 3H), 3.41 – 3.25 (m, 1H), 3.28 (s, 3H), 2.18 – 1.95 (m, 5H), 1.69 – 1.39 (m, 5H).

**$^{13}$C-NMR (101 MHz, CDCl$_3$):** $\delta$ [ppm] = 174.7, 159.8, 153.7, 148.8, 147.1, 140.3, 138.5, 124.8, 116.5, 115.1, 100.2, 100.1, 69.3, 65.7, 61.2, 56.7, 56.3, 48.0, 35.2, 34.3, 33.9, 29.0, 28.2, 24.9.

**HRMS (APCI$^+$, *m/z*):** [M+H]$^+$ for $C_{29}H_{38}N_3O_7S^+$: calcd.: 572.2425, found: 572.2419.

**IR (ATR):** ṽ [cm$^{-1}$] = 3219 (w), 3077 (w), 2933 (m), 1683 (s), 1597 (m), 1498 (s), 1467 (m), 1409 (w), 1330 (m), 1239 (s), 1146 (m), 1129 (s), 1098 (m), 1036 (m), 1006 (m), 913 (w), 847 (w).

**Synthesis of (*R*)-4-((2*R*,4*R*,6*R*)-2-methoxy-6-(pent-4-en-1-yl)-4-(4-((3,4,5-trimethoxy phenyl)diazenyl)phenoxy)tetrahydro-2*H*-pyran-2-yl)thiazolidin-2-one (13):**

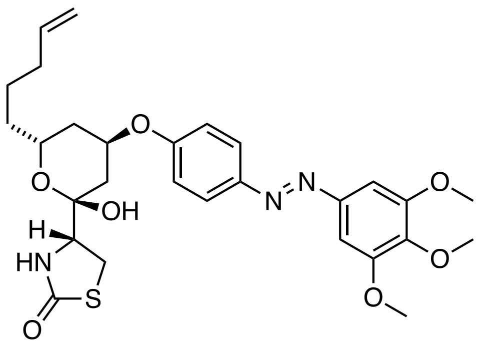


**SI-21** (7.5 mg, 13.1 µmol, 1.00 eq.) was dissolved in THF (0.20 ml) and water (0.20 ml) and AcOH (0.20 ml) was added. The reaction mixture was heated to 50 °C for 4 h. The reaction was poured into sat. aq. $NaHCO_3$ (30 ml) and Ethyl acetate (30 ml). The layers were separated and the aqueous phase was extracted with Ethyl acetate (2 x 20 ml). The combined organic layers were washed with brine, dried over $Na_2SO_4$, filtered and concentrated. The crude product was purified by HPLC (semi-prep, 45-75% MeCN in $H_2O$ over 8 min, 3 min MeCN, no FA) and the title compound was obtained in 55% yield (4.0 mg, 7.2 µmol) as an orange film.

**LCMS** (50-100% MeCN in $H_2O$ over 5 min, 0.1% FA) $t_{ret}$ (minor) = 1.4 min and $t_{ret}$ (major) 2.6 min.

**HPLC** (semi-prep, 45-75% MeCN in $H_2O$ over 8 min, 3 min MeCN, no FA): $t_{ret}$ = 8.3 min (major).

**$^1$H-NMR (600 MHz, $CDCl_3$):** $\delta$ [ppm] = 7.93 (d, $J$ = 8.9 Hz, 2H), 7.23 (s, 2H), 7.04 (d, $J$ = 8.9 Hz, 2H), 5.77 (ddt, $J$ = 16.9, 10.2, 6.7 Hz, 1H), 5.68 (s, 1H), 5.06 (t, $J$ = 3.0 Hz, 1H), 5.02 – 4.92 (m, 2H), 4.18 – 4.12 (m, 1H), 3.97 (s, 6H), 3.93 (s, 3H), 3.90 – 3.86 (m, 1H), 3.56 (dd, $J$ = 11.8, 9.1 Hz, 1H), 3.44 (dd, $J$ = 11.8, 5.4 Hz, 1H), 2.21 (dt, $J$ = 14.4, 2.4 Hz, 1H), 2.06 (dtd, $J$ = 17.6, 9.3, 8.2, 4.3 Hz, 4H), 1.55 – 1.34 (m, 5H).

**$^{13}$C-NMR (151 MHz, $CDCl_3$):** $\delta$ [ppm] = 174.9, 158.2, 153.7, 148.6, 147.9, 140.6, 138.6, 125.0, 116.2, 115.0, 100.3, 97.8, 72.5, 64.7, 61.2, 61.0, 56.4, 35.0, 33.7, 33.4, 31.6, 29.1, 24.6.

**HRMS** (APCI$^+$, *m/z*): $[M+H]^+$ for $C_{28}H_{36}N_3O_7S^+$ calcd.: 558.2268, found: 558.2258.

**IR (ATR):** $\tilde{\nu}$ [cm$^{-1}$] = 3524 (w), 3275 (w), 2934 (w), 2360 (w), 2340 (w), 1680 (s), 1597 (s), 1498 (s), 1467 (m), 1410 (m), 1330 (m), 1232 (s), 1147 (m), 1129 (s), 1005 (m), 902 (w), 847 (w), 806 (w), 773 (w), 720 (w).

**Synthesis of (*R*)-4-((2*R*,4*R*,6*R*)-4-((11,12-dihydrodibenzo[*c*,*g*][1,2]diazocin-3-yl)oxy)-2-methoxy-6-(pent-4-en-1-yl)tetrahydro-2*H*-pyran-2-yl)thiazolidin-2-one (SI-22):**

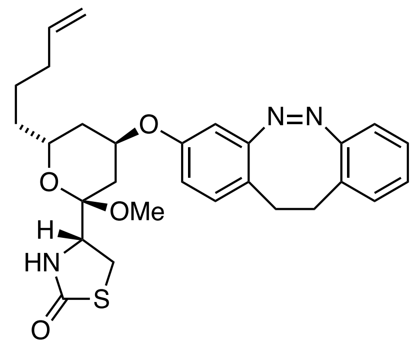


A flame-dried vial with stir bar was charged with **20** (10.0 mg, 33.2 µmol, 1.00 eq.). *meta*-Hydroxy diazocine[11] (14.9 mg, 66.4 mg, 2.00 eq.) and $PPh_3$ (21.8 mg, 82.3 µmol, 2.50 eq.). The reaction mixture was evacuated, backfilled with nitrogen and dissolved in dry toluene (0.6 ml) and dry THF (0.2 ml). The reaction was cooled to 0 °C and DEAD (40 wt% in toluene, 38.0 µl, 83.4 µmol, 2.51 eq.) was added dropwise. The reaction was stirred at 0 °C for 10 minutes and then at room temperature for 20 h. The reaction mixture was loaded onto celite, concentrated to dryness and submitted to flash column chromatography ($SiO_2$, Hex:EtOAc = 9/1 to 5/5) and HPLC (semi-prep, 50-90% MeCN in $H_2O$ over 8 min, 3 min MeCN, no FA). The title compound was obtained as a yellow film (2.72 mg, 5.40 µmol, 16%).

**$R_f$** ($SiO_2$, Hex:EtOAc = 5/5) = 0.38

**LCMS** (Azo; 5-100% MeCN in $H_2O$ over 5 min.): $t_{ret}$ = 4.85 min.

**HPLC** (semi-prep, 50-90% MeCN in $H_2O$ over 8 min, 3 min MeCN, no FA): $t_{ret}$ = 6.5 min.

**$^1$H-NMR (600 MHz, $CDCl_3$):** $\delta$ [ppm] = 7.11 (d, $J$ = 7.8 Hz, 1H), 7.04 – 6.95 (m, 2H), 6.87 (d, $J$ = 8.4 Hz, 1H), 6.80 (t, $J$ = 9.3 Hz, 1H), 6.55 (dd, $J$ = 8.5, 2.5 Hz, 1H), 6.39 (d, $J$ = 2.5 Hz, 1H), 5.83 – 5.73 (m, 1H), 5.41 (s, 1H), 5.04 – 4.94 (m, 2H), 4.62 – 4.59 (m, 1H), 4.12 – 4.05 (m, 1H), 4.02 – 3.91 (m, 1H), 3.33 (t, $J$ = 10.7 Hz, 1H), 3.23 (s, 3H), 3.25 – 3.16 (m, 1H), 2.98 – 2.89 (m, 2H),

---

[11] Maier M. S., Hüll K., Reynders M., Matsuura B. S., Leippe P., Ko T., Schäffer L., Trauner D. *J. Am. Chem. Soc.* **2019**, *141*, 17295–17304.

2.78 – 2.67 (m, 2H), 2.09 – 2.03 (m, 2H), 2.01 – 1.81 (m, 3H), 1.60 – 1.53 (m, 1H), 1.47 – 1.36 (m, 4H).

**$^{13}$C-NMR (151 MHz, $CDCl_3$):** $\delta$ [ppm] = 174.7, 156.4, 156.0*, 155.9*, 155.6, 138.5, 130.8, 129.8, 128.4, 127.3, 126.7, 121.2, 121.2, 118.8, 115.5*, 115.5*, 115.0, 107.5*, 107.4*, 100.0, 69.8*, 69.7*, 65.6, 56.6, 48.0, 35.1, 34.2, 33.9, 31.8, 31.1*, 31.1*, 29.1*, 28.7*, 28.1, 24.8.
* split signals.

**HRMS** (APCI$^+$, *m/z*): $[M+H]^+$ for $C_{28}H_{35}N_3O_4S^+$ calcd.: 508.2265, found: 508.2258.

**IR (ATR):** v (tilde) = 3206 (w), 3068 (w), 2925 (m), 2855 (w), 2361 (w), 1683 (s), 1607 (w), 1568 (w), 1492 (w), 1458 (w), 1354 (w), 1263 (w), 1241 (m), 1135 (w), 1100 (m), 1035 (m), 909 (w), 814 (w), 755 (w), 726 (w).

**Synthesis of (R)-4-((2R,4R,6R)-4-(((Z)-11,12-dihydrodibenzo[c,g][1,2]diazocin-3-yl)oxy)-2-hydroxy-6-(pent-4-en-1-yl)tetrahydro-2H-pyran-2-yl)thiazolidin-2-one (11):**

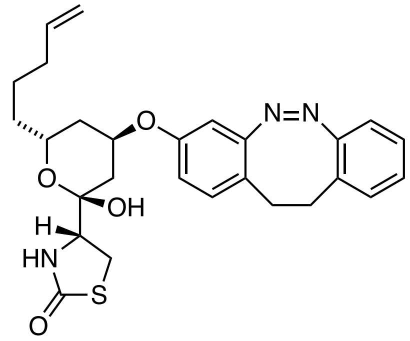


**SI-22** (2.72 mg, 5.30 µmol, 1.00 eq.) was dissolved in THF (0.1 ml), $H_2O$ (0.1 ml) and AcOH (0.1 ml) was added. the reaction was stirred for 3 h at 50 °C. The reaction was cooled to room temperature, poured into $NaHCO_3$ (sat. aq., 30 ml) and EtOAc (30 ml). The layers were separated and the aqueous layer was extracted with EtOAc (2 x 20 ml). The combined organic layers were washed with brine, dried over $Na_2SO_4$, filtered and concentrated. The crude product was purified by HPLC (semi-prep, 45-75% MeCN in $H_2O$ over 8 min, 3 min MeCN, no FA) and gave **11** as a pale yellow film (1.40 mg, 2.80 µmol, 53%).

**LCMS** (50-100% MeCN in H2O over 5 min, 0.1% FA): 2.2 (major) and 2.7 (minor)

**HPLC** (semi-prep, 45-75% MeCN in $H_2O$ over 8 min, 3 min MeCN, no FA): $t_{ret}$ = 7.3 min.

**$^1$H-NMR (600 MHz, $CDCl_3$):** $\delta$ [ppm] = 7.17 – 7.13 (m, 1H), 7.04 (td, $J$ = 7.5, 1.2 Hz, 1H), 7.00 – 6.97 (m, 1H), 6.92 (d, $J$ = 8.4 Hz, 1H), 6.85 – 6.81 (m, 1H), 6.57 (d, $J$ = 8.5 Hz, 1H), 6.38 (s, 1H), 5.83 – 5.70 (m, 1H), 5.60 (s, 1H), 5.03 – 4.93 (m, 3H), 4.81 (s, 1H), 4.07 – 4.00 (m, 1H), 3.83 (dd, $J$ = 9.1, 5.4 Hz, 1H), 3.52 (dd, $J$ = 11.8, 9.1 Hz, 1H), 3.41 – 3.31 (m, 1H), 2.98 – 2.90 (m, 2H), 2.79 – 2.71 (m, 2H), 2.11 – 1.83 (m, 5H), 1.50 – 1.28 (m, 5H).

**$^{13}$C-NMR (151 MHz, $CDCl_3$):** $\delta$ [ppm] = 174.8, 156.5*, 156.5*, 155.5*, 155.4*, 154.7, 138.6, 131.4*, 131.3*, 129.9, 128.2, 127.4, 126.8, 122.4*, 118.8, 118.8, 115.4*, 115.0, 114.7*, 106.8*, 106.2*, 97.8, 72.7, 64.6, 60.9, 35.0, 33.7, 33.2*, 31.8*, 31.7*, 31.5, 31.1, 29.9, 29.0, 24.5.

* split peak or two peaks that correspond to the same carbon. Splitting pattern due to conformation of diazocine relative to latrunculin core.

**HRMS** (APCI⁺, *m/z*): $[M+H]^+$ = calcd. for $C_{27}H_{32}N_3O_4S^+$: 494.2108; found: 494.2106.

**IR (ATR):** ṽ [$cm^{-1}$] = 3507 (w), 3227 (w), 3069 (w), 2923 (s), 2853 (m), 2361 (w), 2341 (w), 1678 (w), 1607 (m), 1571 (w), 1491 (m), 1458 (m), 1409 (m), 1349 (m), 1261 (s), 1237 (s), 1100 (s), 912 (m), 806 (s), 754 (m), 722 (m).

**Synthesis of (*R*)-4-((2*R*,4*R*,6*R*)-4-(4-((2,3-dihydrobenzo[*b*][1,4]dioxin-6-yl)diazenyl) phenoxy)-2-methoxy-6-(pent-4-en-1-yl)tetrahydro-2*H*-pyran-2-yl)thiazolidin-2-one (SI-23):**

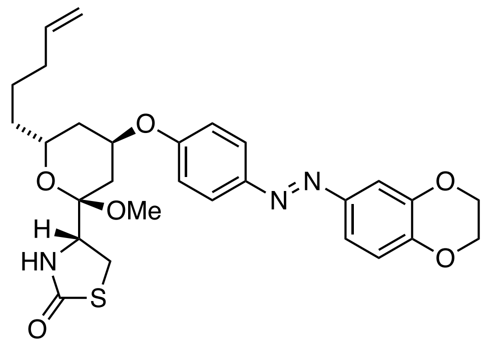


**20** (17.1 mg, 56.7 µmol, 1.00 eq.), $PPh_3$ (37.2 mg, 124 µmol, 2.50 eq.) and **SI-29** (29.1 mg, 0.114 mmol, 2.00 eq.) were dissolved in dry THF (0.34 ml) and dry toluene (1.00 ml) and cooled to 0 °C using an ice-bath. To the orange reaction mixture, DEAD (40% in toluene, 65.0 µl, 142 µmol, 2.50 eq.) was added and dropwise and the reaction turned dark red. The reaction was stirred at 0 °C for 30 min. before it was warmed to room temperature and stirred over night. The reaction mixture was loaded on isolute, concentrated *in vaccuo* and submitted to flash column chromatography ($SiO_2$, Hex:EtOAc 10:0 to 5:5 to 0:10, slow gradient). The purified product contained $H_2$DEAD and was submitted ketal cleavage without further purification.

**R*f*** ($SiO_2$, Hex:EtOAc = 5:5, CAM) = 0.57.

**LCMS** (5-100% MeCN in $H_2O$ over 5 min, 0.1% FA) $t_{ret}$ = 3.6 min (major).

**Synthesis of (R)-4-((2R,4R,6R)-4-(4-((E)-(2,3-dihydrobenzo[b][1,4]dioxin-6-yl)diazenyl) phenoxy)-2-hydroxy-6-(pent-4-en-1-yl)tetrahydro-2H-pyran-2-yl)thiazolidin-2-one (12):**

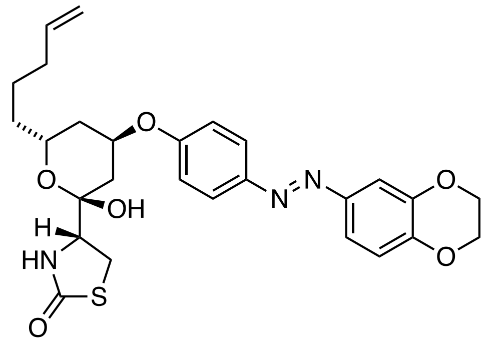

**SI-23** was dissolved in THF (0.90 ml) and water (0.90 ml) and AcOH (0.90 ml) was added. The reaction mixture was heated to 50 °C for 6 h. The reaction was poured into sat. aq. $NaHCO_3$ (30 ml) and EtOAc (30 ml). The layers were separated and the aqueous phase was extracted with EtOAc (2x 20 ml). The combined organic layers were washed with brine, dried over $Na_2SO_4$, filtered and concentrated. The crude product was purified by HPLC (semi-prep, 50-95% MeCN in $H_2O$ over 8 min, 3 min MeCN, no FA) and the title compound was obtained as an orange film (1.29 mg, 2.5 µmol, 4% over 2 steps).

**$R_f$** ($SiO_2$, Hex:EtOAc = 5:5, CAM) = 0.29.

**LCMS** (AZO, 50-100% MeCN in $H_2O$, 0.1% FA) $t_{ret}$ = 2.9 (major).

**HPLC** (semi-prep, 50-95% MeCN in $H_2O$ over 8 min, 3 min MeCN, no FA): $t_{ret}$ = 6.4 min.

**$^1$H-NMR (600 MHz, $CDCl_3$):** $\delta$ [ppm] = 7.89 (d, $J$ = 9.0 Hz, 2H), 7.49 (dd, $J$ = 8.6, 2.1 Hz, 1H), 7.46 (d, $J$ = 2.4 Hz, 1H), 7.02 (d, $J$ = 9.0 Hz, 2H), 6.98 (dd, $J$ = 8.6, 1.6 Hz, 1H), 5.77 (ddt, $J$ = 16.8, 9.7, 6.6 Hz, 1H), 5.67 – 5.59 (m, 1H), 5.05 (s, 1H), 5.03 (s, 1H), 5.02 – 4.92 (m, 2H), 4.36 – 4.29 (m, 4H), 4.18 – 4.11 (m, 1H), 3.88 (dd, $J$ = 9.2, 5.5 Hz, 1H), 3.55 (td, $J$ = 10.0, 9.0, 1.6 Hz, 1H), 3.43 (dd, $J$ = 11.8, 5.4 Hz, 1H), 2.23 – 2.18 (m, 1H), 2.09 – 2.02 (m, 4H), 1.52 – 1.37 (m, 5H).

**$^{13}$C-NMR (151 MHz, $CDCl_3$):** $\delta$ [ppm] = 174.8, 157.9, 148.1, 147.5, 146.4, 144.0, 138.6, 124.8, 118.2, 117.6, 116.2, 115.0, 110.8, 97.9, 72.6, 64.8, 64.7, 64.4, 61.0, 35.1, 33.7, 33.5, 31.6, 29.1, 24.6.

**HRMS** (APCI$^+$, *m/z*): [(M+H)$^+$] for $C_{27}H_{32}N_4O_6S^+$ calcd.: 526.2006, found: 526.1983.

**IR (ATR):** $\tilde{v}$ [cm$^{-1}$] = 3521 (w), 3229 (w), 3073 (w), 2924 (m), 2853 (m), 2363 (w), 1676 (s), 1596 (m), 1581 (m), 1494 (s), 1456 (m), 1320 (m), 1286 (s), 1257 (s), 1231 (s), 1137 (m), 1095 (s), 1063 (s), 883 (m), 838 (m), 803 (s), 724 (m).

**Synthesis of (R)-4-((2R,4R,6R)-4-(4-((4-benzyl-3,4-dihydro-2H-benzo[b][1,4]oxazin-7yl)diaz-enyl)phenoxy)-2-methoxy-6-(pent-4-en-1-yl)tetrahydro-2H-pyran-2-yl)thiazolidin-2-one (SI-24):**

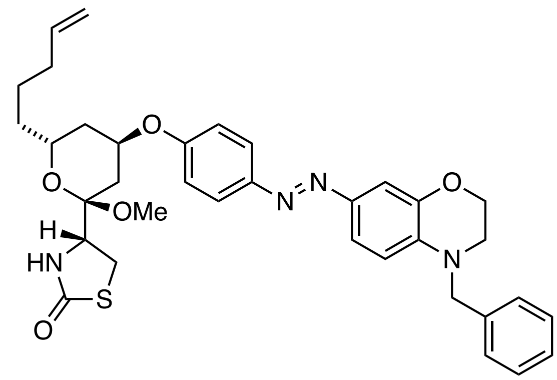


**20** (10.0 mg, 0.03 mmol, 1.00 eq.), $PPh_3$ (26.1 mg, 0.10 mmol, 3.00 eq.) and **21** (22.9 mg, 0.07 mmol, 2.00 eq.) were dissolved in dry THF (0.34 ml) and dry toluene (1.00 ml) and cooled to 0 °C using an ice-bath. To the orange reaction mixture, TMAD (17.1 mg, 0.10 mmol, 3.00 eq.) was added dropwise and the reaction turned dark red. The reaction was stirred at 0 °C for 30 min. before it was warmed to room temperature and stirred overnight. The reaction mixture was loaded on isolute, concentrated *in vaccuo* and submitted to flash column chromatography ($SiO_2$, Hex:EtOAc = 10:0 to 5:5 to 0:10, slow gradient). The purified product was immediately submitted ketal cleavage.

**$R_f$** ($SiO_2$, Hex:EtOAc = 5:5, CAM) = 0.59

**LCMS** (50-100% MeCN in $H_2O$ over 5 min, 0.1% FA): $t_R$ (major) = 4.3 min (trans).

**Synthesis of (R)-4-((2R,4R,6R)-4-(4-((4-benzyl-3,4-dihydro-2H-benzo[b][1,4]oxazin-7yl)diaz enyl)phenoxy)-2-hydroxy-6-(pent-4-en-1-yl)tetrahydro-2H-pyran-2-yl)thiazolidin-2-one (14):**

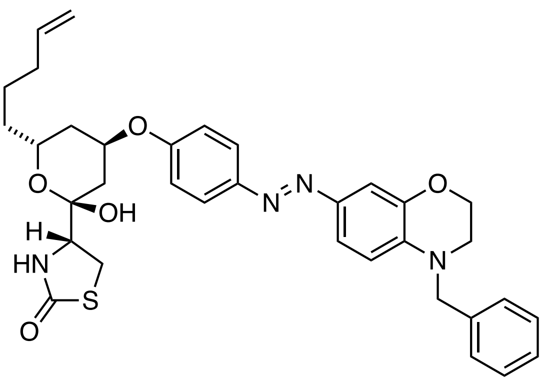


**SI-24** was dissolved in THF (0.90 ml) and water (0.90 ml) and AcOH (0.90 ml) was added. The reaction mixture was heated to 50 °C for 6 h. The reaction was poured into sat. aq. $NaHCO_3$ (30 ml) and EtOAc (30 ml). The layers were separated and the aqueous phase was extracted with EtOAc (2x 20 ml). The combined organic layers were washed with brine, dried over $Na_2SO_4$, filtered and concentrated. The crude product was purified by HPLC (semi-prep, 50-96% MeCN in $H_2O$ over 8 min, 3 min MeCN, no FA) and the title compound was obtained as an orange film (9.8 mg, 15.9 µmol, 48% over 2 steps).

$\mathbf{R}_f$ ($SiO_2$, Hex:EtOAc = 5:5, CAM) = 0.31.

**LCMS** (5-100% MeCN in $H_2O$ over 5 min, 0.1% FA) $t_{ret}$ = 5.3 min.

**HPLC** (semi-prep, 50-96% MeCN in $H_2O$ over 8 min, 3 min MeCN, no FA): $t_{ret}$ = 7.7 min.

**$^1$H-NMR (600 MHz, $CDCl_3$):** $\delta$ [ppm] = 7.84 (d, $J$ = 8.5 Hz, 2H), 7.46 (d, $J$ = 2.0 Hz, 1H), 7.45 (s, 1H), 7.36 (t, $J$ = 7.5 Hz, 2H), 7.28 (d, $J$ = 7.8 Hz, 3H), 7.00 (d, $J$ = 8.6 Hz, 2H), 6.74 (d, $J$ = 8.4 Hz, 1H), 5.77 (ddt, $J$ = 16.9, 10.2, 6.7 Hz, 1H), 5.64 (s, 1H), 5.09 (s, 1H), 5.02 (s, 1H), 5.01 – 4.92 (m, 2H), 4.59 (s, 2H), 4.30 (t, $J$ = 4.5 Hz, 2H), 4.14 (td, $J$ = 7.7, 7.1, 4.1 Hz, 1H), 3.88 (dd, $J$ = 9.1, 5.6 Hz, 1H), 3.55 (dd, $J$ = 11.7, 9.2 Hz, 1H), 3.50 (t, $J$ = 4.5 Hz, 2H), 3.43 (dd, $J$ = 11.8, 5.5 Hz, 1H), 2.19 (d, $J$ = 14.3 Hz, 1H), 2.08 – 2.01 (m, 4H), 1.53 – 1.37 (m, 5H).

**$^{13}$C-NMR (151 MHz, $CDCl_3$):** $\delta$ [ppm] = 174.8, 157.2, 148.5, 144.7, 144.0, 138.6, 137.2, 129.0, 127.6, 127.0, 124.4, 121.0, 116.3, 115.0, 111.2, 108.6, 97.9, 72.6, 64.7, 64.4, 61.0, 54.7, 47.5, 35.1, 33.7, 33.5, 31.6, 29.1, 24.6.

**HRMS (ESI$^+$, *m/z*):** [(M+H)$^+$] for $C_{34}H_{39}N_4O_5S^+$ calcd.: 615.2636, found: 615.2645.

**IR (ATR):** ṽ [cm$^{-1}$] = 3528 (w), 3227 (w), 3070 (w), 2924 (m), 2854 (w), 1678 (s), 1596 (s), 1515 (s), 1496 (s), 1452 (m), 1396 (m), 1350 (m), 1319 (s), 1232 (s), 1201 (s), 1152 (m), 1093 (s), 1051 (s), 884 (m), 838 (m), 802 (m), 727 (m).

**Synthesis of (R)-4-((2R,4R,6R)-4-(4-((E)-(4-benzyl-3,4-dihydro-2H-benzo[b][1,4]oxazin-6-yl)diazenyl)phenoxy)-2-hydroxy-6-(pent-4-en-1-yl)tetrahydro-2H-pyran-2-yl)thiazolidin-2-one (15):**

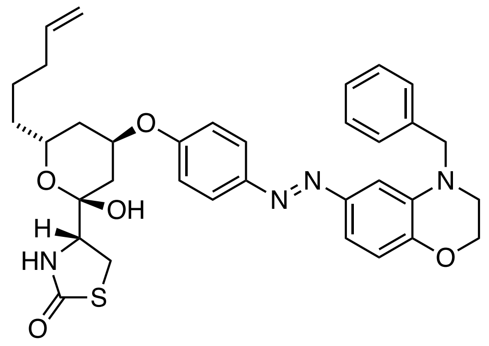


**20** (5.00 mg, 17.0 µmol, 1.00 eq.), $PPh_3$ (13.1 mg, 50.0 µmol, 3.00 eq.) and (*E*)-4-((4-benzyl-3,4-dihydro-2H-benzo[b][1,4]oxazin-6-yl)diazenyl)phenol[12] (11.5 mg, 33.0 µmol, 2.00 eq.) were dissolved in dry THF (0.17 ml) and dry toluene (0.50 ml) and cooled to 0 °C using an ice-bath. To the orange reaction mixture, TMAD (8.60 mg, 50.0 µmol, 3.00 eq.) was added dropwise and the reaction turned dark red. The reaction was stirred at 0 °C for 30 min. before it was warmed to room temperature and stirred overnight. The reaction mixture was loaded on isolute, concentrated *in vaccuo,* and submitted to flash column chromatography ($SiO_2$, Hex:EtOAc = 10:0 to 5:5 to 0:10, slow gradient). The purified product was submitted ketal cleavage without further purification.

The hemiacetal intermediate was dissolved in THF (0.45 ml) and water (0.45 ml) and AcOH (0.45 ml) was added. The reaction mixture was heated to 50 °C for 6 h. The reaction was poured

[12] prepared analogously to **21**.

into sat. aq. $NaHCO_3$ (10 ml) and EtOAc (10 ml). The layers were separated, and the aqueous phase was extracted with EtOAc (2x 10 ml). The combined organic layers were washed with brine, dried over $Na_2SO_4$, filtered and concentrated. The crude product was purified by HPLC (semi-prep, 50-96% MeCN in $H_2O$ over 8 min, 3 min MeCN, no FA) and the title compound was obtained as an orange film (4.9 mg, 8.00 µmol, 48% over 2 steps).

**$R_f$** ($SiO_2$, Hex:EtOAc = 5:5, CAM) = 0.31.

**LCMS** (5-100% MeCN in $H_2O$ over 5 min, 0.1% FA) $t_{ret}$ = 5.3 min.

**HPLC** (semi-prep, 50-96% MeCN in $H_2O$ over 8 min, 3 min MeCN, no FA): $t_{ret}$ = 7.7 min.

**$^1$H-NMR (600 MHz, $CDCl_3$):** $\delta$ [ppm] = 7.84 (d, $J$ = 8.6 Hz, 1H), 7.38 – 7.32 (m, 5H), 7.28 (m, 1H), 7.00 (d, $J$ = 8.7 Hz, 2H), 6.93 (d, $J$ = 8.4 Hz, 1H), 5.82 – 5.70 (m, 1H), 5.05 (s, 1H), 5.08 – 4.92 (m, 3H), 4.55 (s, 2H), 4.32 (t, $J$ = 4.4 Hz, 2H), 4.18 – 4.0 (m, 1H), 3.87 (dd, $J$ = 9.1, 5.5 Hz, 1H), 3.54 (dd, $J$ = 11.8, 9.1 Hz, 1H), 3.43 (dd, $J$ = 11.8, 5.5 Hz, 1H), 3.35 (t, $J$ = 4.5 Hz, 2H), 2.19 (d, $J$ = 14.2 Hz, 1H), 2.09 – 2.0 (m, 4H), 1.53 – 1.34 (m, 6H).

**$^{13}$C-NMR (151 MHz, $CDCl_3$):** $\delta$ [ppm] = 174.7, 157.4, 148.1, 147.71, 147.0, 138.4, 137.5, 136.0, 128.7, 127.4, 127.4, 124.5, 116.6, 116.0, 115.1, 114.8, 105.3, 97.7, 72.4, 65.0, 64.5, 60.8, 54.6, 46.3, 34.9, 33.5, 33.3, 31.5, 28.9, 24.4.

**HRMS (ESI$^+$, *m/z*):** [$(M+Na)^+$] for $C_{34}H_{38}N_4NaO_5S^+$ calcd.: 637.2455, found: 637.2480.

**IR (ATR):** $\tilde{\nu}$ [$cm^{-1}$] = 2931 (w), 2862 (w), 1683 (w), 1598 (m), 1505 (w), 1452 (s), 1345 (s), 1151 (s), 1094 (m), 1040 (m), 852 (s).

**Synthesis of (4-(phenyldiazenyl)phenyl)methanol (SI-25)**

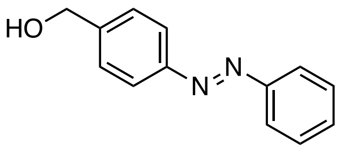

The title compound was synthesized following a reported procedure[13]. Yield: 54%; The analytical data matched those reported.

**$R_f$** ($SiO_2$, Hex:EtOAc = 8:2; UV) = 0.18

**$^1$H-NMR (400 MHz, $CDCl_3$):** $\delta$ [ppm] 7.97 – 7.88 (m, 4H), 7.57 – 7.43 (m, 5H), 4.80 (d, $J$ = 5.6 Hz, 2H), 1.75 (t, $J$ = 5.9 Hz, 1H).

**$^{13}$C-NMR (101 MHz, $CDCl_3$):** $\delta$ [ppm] = 152.8, 152.3, 144.0, 131.2, 129.2, 127.6, 123.2, 123.0, 65.1.

[13] Rao, J.; Hottinger, C.; Khan, A. *J. Am. Chem. Soc.* **2014**, *136*, 5872–5875.

**Synthesis of 2-(4-(phenyldiazenyl)phenyl)ethan-1-ol (SI-26):**

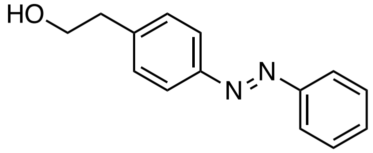

The title compound was synthesized following a reported procedure[14]. Yield: 59%; The analytical data matched those reported.

**$R_f$** ($SiO_2$, Hex:EtOAc = 8:2; UV) = 0.15

**$^1$H-NMR (400 MHz, $CDCl_3$):** $\delta$ [ppm] = 7.95 – 7.82 (m, 4H), 7.59 – 7.43 (m, 3H), 7.43 – 7.32 (m, 2H), 3.98 – 3.87 (m, 2H), 2.96 (t, $J$ = 6.5 Hz, 2H), 1.45 – 1.39 (m, 1H).

**$^{13}$C-NMR (101 MHz, $CDCl_3$):** $\delta$ [ppm] = 152.8, 151.6, 142.1, 131.0, 129.9, 129.2, 123.3, 122.9, 63.6, 39.2.

**Synthesis of 2-(phenyldiazenyl)benzoic acid (SI-27):**

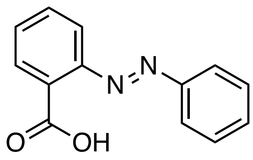

2-Aminobenzoic acid (461 mg, 3.36 mmol, 1.20 eq.) was dissolved in glacial acetic acid (4 ml) at room temperature and a solution of nitroso benzene (300 mg, 2.80 mmol, 1.00 eq.) in acetic acid (4 ml) was added dropwise over 5 min. The brown reaction mixture was stirred at room temperature for 48 h and then concentrated under reduced pressure. The crude product was

[14] Schönberger, M; Trauner, D. *Angew. Chem. Int. Ed.* **2014**, *53*, 3264–4367.

purified by flash column chromatography ($SiO_2$, DCM:MeOH 10:0 to 9:1) and gave the title compound as an orange-brown solid.

**R**$_f$ (DCM; UV) = 0.54 (major)

**$^1$H-NMR (400 MHz, $CDCl_3$):** $\delta$ [ppm] = 12.83 ($s_{br}$, 1H), 8.51 – 8.44 (m, 1H), 8.09 – 8.02 (m, 1H), 7.92 – 7.86 (m, 2H), 7.75 – 7.67 (m, 2H), 7.64 – 7.58 (m, 3H).

**$^{13}$C-NMR (101 MHz, $CDCl_3$):** $\delta$ [ppm] = 166.2, 151.7, 149.5, 134.0, 133.7, 133.4, 132.9, 130.0, 127.3, 123.8, 115.9.

**HRMS (ESI$^+$, *m/z*):** $[M+H]^+$ for $C_{13}H_{11}N_2O_2^+$ calcd.: 227.0815, found: 227.0816.

**Synthesis of 3-(phenyldiazenyl)benzoic acid (SI-27):**

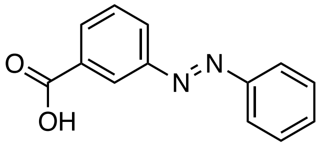


3-Aminobenzoic acid (400 mg, 3.70 mmol, 1.00 eq.) was dissolved in glacial acetic acid (3 ml) at room temperature and a solution of nitroso benzene (615 mg, 4.50 mmol, 1.20 eq.) in acetic acid (4.5 ml) was added dropwise over 5 min. The brown reaction mixture was stirred at room temperature for 48 h and then concentrated under reduced pressure. The crude product was purified by flash column chromatography ($SiO_2$, DCM:MeOH 10:0 to 9:1) and gave the title compound as an orange solid (559 mg, 2.48 mmol, 55%).

**R**$_f$ ($SiO_2$, DCM:MeOH = 95:5; UV) = 0.4

**$^1$H-NMR (400 MHz, $CDCl_3$):** $\delta$ [ppm] = 8.66 (d, $J$ = 2.1 Hz, 1H), 8.28 – 8.20 (m, 1H), 8.18 (dd, $J$ = 7.9, 1.7 Hz, 1H), 8.02 – 7.92 (m, 2H), 7.65 (t, $J$ = 7.8 Hz, 1H), 7.60 – 7.44 (m, 4H).

**$^{13}$C-NMR (101 MHz, CDCl$_3$):** $\delta$ [ppm] = 170.6, 152.7, 152.4, 132.2, 131.5, 130.3, 129.4, 129.2, 127.7, 124.8, 123.1.

**HRMS (ESI$^+$, *m/z*):** for $C_{13}H_{11}N_2O_2^+$ calcd.: 227.0815, found: 227.0823.

**IR (ATR):** ṽ [cm$^{-1}$] = 2817 (w), 2565 (w), 1674 (s), 1598 (w), 1585 (m), 1473 (w), 1448 (w), 1423 (m), 1326 (m), 1307 (m), 1277 (m), 1211 (m), 1159 (m), 1093 (w), 1077 (m), 1019 (w), 998 (m), 937 (m), 917 (m), 820 (m), 785 (m), 762 (s)

**Synthesis of 4-((3,4,5-trimethoxyphenyl)diazenyl)phenol (SI-28):**

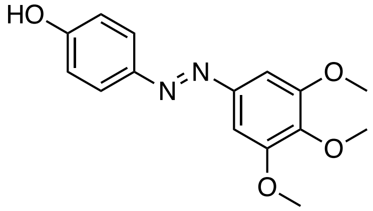


Following a reported procedure[15] for a related compound, 3,4,5-trimethoxyaniline (366 mg, 2.00 mmol, 1.00 eq.) was dissolved in THF (5.0 ml) and HCl (1M, 6.0 ml) and cooled to 0 °C. $H_2O$ (4.0 ml) was added and an aqueous solution of $NaNO_2$ (2.0 M in $H_2O$, 1.2 ml, 2.40 mmol, 1.20 eq.) was added dropwise and the resulting dark-red suspension was stirred for 30 min. at 0 °C. A solution of phenol (226 mg, 2.40 mmol, 1.20 eq.) was added dropwise and the reaction mixture was stirred for 1 h at 0 °C. The volatiles were removed under reduced pressure and the residue was redissolved in EtOAc (20 ml) and $H_2O$ (20 ml) and the phases separated. The aqueous layer was extracted with EtOAc (3 x 30 ml) and the combined organic layers were washed with water and brine, dried over $MgSO_4$, filtered and concentrated. The crude product was purified by flash column chromatography ($SiO_2$, Hex:EtOAc = 8:2 to 5:5) and gave the title compound as a brown solid (526 mg, 1.82 mmol, 91%).

---

[15] Eisel, B.; Hartrampf, F. W. W.; Meier, T.; Trauner, D. *FEBS lett.* **2018**, *592*, 343–355.

**$R_f$** ($SiO_2$, Hex:EtOAc = 6:4; UV) = 0.33 (major), 0.47 (minor).

**$^1$H-NMR (400 MHz, $CDCl_3$)**: $\delta$ [ppm] = 7.86 (d, $J$ = 8.8 Hz, 2H), 7.22 (s, 2H), 6.95 (d, $J$ = 8.8 Hz, 2H), 5.17 ($s_{br}$, 1H), 3.96* (s, 6H), 3.93* (s, 3H). *main peaks reported

**$^{13}$C-NMR (101 MHz, $CDCl_3$):** $\delta$ [ppm] = 158.3, 153.7, 148.7, 147.2, 140.3, 125.0, 116.0, 100.3, 61.2, 56.4.

**HRMS (ESI$^+$, *m/z*):** $[M+H]^+$ for $C_{16}H_{16}N_2O_4^+$ calcd.: 289.1183, found: 289.1180.

**IR (ATR):** ṽ [$cm^{-1}$] = 3277 (w), 2950 (w), 2839 (w), 1599 (m), 1585 (m), 1496 (m), 1472 (m), 1461 (m), 1430 (w), 1411 (m), 1332 (m), 1316 (w), 1302 (w), 1278 (m), 1225 (m), 1208 (m), 1180 (m), 1145 (m), 1128 (s), 1100 (m), 991 (s), 922 (w), 866 (m), 847 (s), 781 (m).

**Synthesis of 4-((2,3-dihydrobenzo[b][1,4]dioxin-6-yl)diazenyl)phenol (SI-29):**

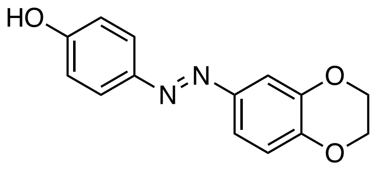


2,3-dihydrobenzo[*b*][1,4]dioxin-6-amine (500 mg, 3.31 mmol, 1.00 eq.) was dissolved in THF (8.3 ml), water (6.3 ml) and 1 M HCl (9.9 ml) and cooled to 0 °C. An aqueous solution of $NaNO_2$ (2 M, 2.0 ml, 3.97 mmol, 1.20 eq.) was added dropwise and the reaction was stirred for 30 minutes at 0 °C. Phenol (374 mg, 3.97 µmol, 1.20 eq.) was dissolved in aqueous NaOH (1 M, 9.9 ml, 9.92 mmol, 3.00 eq.) and added dropwise. The reaction was stirred for 2 h at 0 °C and upon completion, the volatiles were removed under reduced pressure. The residue was taken up in EtOAc (100 ml) and $H_2O$ (50 ml), the layers were separated and the organic layer was extracted with EtOAc (2 x 100 ml). The combined organic layers were washed with brine, dried over $MgSO_4$, filtered and concentrated. The crude product was purified by flash column chromatography ($SiO_2$,

Hex:DCM:EtOAc = 6:3:1 to 0:6:2) and the title compound was obtained as an orange solid (531 mg, 2.07 mmol, 63%).

**R*f*** ($SiO_2$, Hex:DCM:EtOAc = 6:3:1; UV) = 0.22

**$^1$H-NMR (400 MHz, $CDCl_3$)**: $\delta$ = 7.86 – 7.79 (m, 2H), 7.49 – 7.43 (m, 2H), 6.97 (d, $J$ = 8.4 Hz, 1H), 6.95 – 6.89 (m, 2H), 5.17 (s, 1H), 4.34 – 4.27 (m, 4H).

**$^{13}$C-NMR (101 MHz, $CDCl_3$):** δ 157.9, 147.6, 147.3, 146.1, 144.0, 124.8, 117.9, 117.5, 115.9, 110.7, 64.7, 64.4.

**HRMS** (ESI$^+$, *m/z*): [(M+H)$^+$] for $C_{14}H_{13}N_2O_3^+$ calcd.: 257.0921, found: 257.0920.

**IR (ATR):** ṽ [cm$^{-1}$] = 3378 (w), 1592 (m), 1491 (s), 1434 (w), 1318 (w), 1284 (s), 1268 (m), 1255 (s), 1239 (m), 1212 (m), 1190 (m), 1138 (m), 1115 (m), 1099 (w), 1059 (s), 1037 (m), 923 (m), 896 (m), 884 (m), 867 (m), 835 (s), 809 (m), 762 (m), 741 (m), 724 (w).

**Synthesis of 7-nitro-3,4-dihydro-2*H*-benzo[*b*][1,4]oxazine (SI-30):**

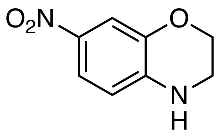


2-Amino-5-nitrophenol (10.0 g, 64.9 mmol, 1.00 eq.) and $K_2CO_3$ (35.7 g, 260 mmol, 4.00 eq.) were dissolved in dry DMF (130 ml) and 1,2-dibromoethane (22.4 ml, 260 mmol, 4.00 eq.) was added. The reaction was heated to 120 °C over night. Upon cooling to room temperature, DMF was removed under reduced pressure and by trituation with hexanes. The resulting crude product was partitioned between $H_2O$ (400 ml) and DCM (400 ml) and the layers were separated. The organic layer was dried over $MgSO_4$ and concentrated. Excess DMF was removed by co evaporation with hexanes. The title compound was obtained as a yellow solid (9.43 g, 52.3 mmol, 81%).

**$^1$H-NMR (400 MHz, $CDCl_3$):** $\delta$ [ppm] = 7.78 – 7.66 (m, 2H), 6.52 (d, $J$ = 8.7 Hz, 1H), 4.29 – 4.21 (m, 2H), 3.61 – 3.49 (m, 2H), 2.96 (s, 1H), 2.88 (s, 1H).

**Synthesis of 4-benzyl-7-nitro-3,4-dihydro-2H-benzo[b][1,4]oxazine (SI-31):**

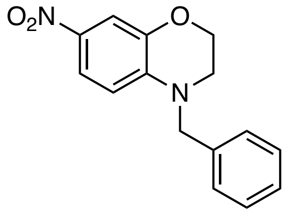

Following a known procedure[16], 7-nitro-3,4-dihydro-2*H*-benzo[*b*][1,4]oxazine **SI-30** (400 mg, 2.22 mmol, 1.00 eq.) and $K_2CO_3$ (1.04 g, 7.55 mmol, 3.40 eq.) were dissolved in dry DMF (3.2 ml) and benzyl bromide (0.53 ml, 4.44 mmol, 2.00 eq.) was added. The reaction mixture was heated to 80 °C for 22 h. After cooling to room temperature the reaction mixture was partitioned between $H_2O$ (100 ml) and EtOAc (100 ml). The phases were separated and the aqueous layer was extracted with EtOAc (2 x 100 ml). The combined organic layers were washed with brine, dried over $Na_2SO_4$, filtered and concentrated. The crude product was purified by flash column chromatography ($SiO_2$, Hex:EtOAc = 8:2 to 0:10) and gave the title compound as a yellow solid (348 mg, 1.29 mmol, 58%).

**$R_f$** ($SiO_2$, Hex:EtOAc = 8:2; UV) = 0.32

**$^1$H-NMR (400 MHz, $CDCl_3$):** $\delta$ [ppm] = 7.58 (s, 1H), 7.60 – 7.55 (m, 1H), 7.42 – 7.33 (m, 2H), 7.33 – 7.26 (m, 3H), 6.84 (dt, $J$ = 8.9, 1.3 Hz, 1H), 4.51 (s, 2H), 4.36 – 4.32 (m, 2H), 3.42 – 3.37 (m, 2H).

---

[16] Beijing University of Technology; Hu, L.; Rong, J.; Mao, Z.; Wang, Y.; Zeng C. CN104693216, 2017, B; Location in patent: Paragraph 0062-0066

**$^{13}$C-NMR (101 MHz, CDCl$_3$):** $\delta$ [ppm] = 149.5, 142.6, 136.5, 135.6, 129.1, 127.9, 127.4, 116.2, 114.4, 107.3, 65.2, 54.8, 46.2.

**HRMS (ESI$^+$, *m/z*):** [M+H]$^+$ for $C_{15}H_{15}N_2O_3^+$: calcd.: 271.1077, found: 271.1075.

**IR (ATR):** $\tilde{\nu}$ [cm$^{-1}$] = 2850 (w), 1722 (w), 1617 (w), 1575 (w), 1512 (s), 1452 (m), 1321 (s), 1235 (s), 1215 (s), 1161 (m), 1105 (s), 1076 (w), 1039 (m), 985 (w), 935 (w), 913 (m), 855 (m), 803 (m), 741 (s), 714 (m).

**Synthesis of 4-benzyl-3,4-dihydro-2H-benzo[b][1,4]oxazin-7-amine (SI-32):**

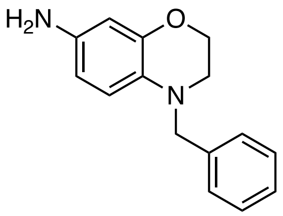


**SI-31** (348 mg, 1.29 mmol, 1.00 eq.) and $SnCl_2 \cdot 2\,H_2O$ (1.16 g, 5.15 mmol, 4.00 eq.) were dissolved in ethanol (5.1 ml) and conc. HCl (1.0 ml) and heated to 80 °C for 16 h. The reaction was cooled to room temperature and poured slowly into sat. aq. $NaHCO_3$ (80 ml). The mixture was basified with NaOH (2.0 M) and $H_2O$ (30 ml) was added. The aqueous layer was extracted with EtOAc (3 x 100 ml), washed with brine, dried over $Na_2SO_4$, filtered and concentrated. The crude product was purified by flash column chromatography ($SiO_2$, Hex:DCM:EtOAc 7:2:1 to 0:8:2) and furnished the title compound as a light beige solid (285 mg, 1.19 mmol, 92%). The product was unstable and darkened when kept under air for prolonged times and was thus kept under high vacuum for 24 h and used immediately.

**$R_f$** ($SiO_2$, Hex:DCM:EtOAc = 4:4:2) = 0.30.

$^1$**H-NMR (400 MHz, CDCl$_3$):** $\delta$ [ppm] = 7.35 – 7.31 (m, 4H), 7.29 – 7.23 (m, 1H), 6.56 (d, $J$ = 8.4 Hz, 1H), 6.26 (d, $J$ = 2.4 Hz, 1H), 6.20 (dd, $J$ = 8.5, 2.6 Hz, 1H), 4.31 (s, 2H), 4.26 – 4.21 (m, 2H), 3.32 ($s_{br}$, 2H), 3.22 – 3.17 (m, 2H). *contains residual EtOAc.

**Synthesis of 4-((4-benzyl-3,4-dihydro-2H-benzo[b][1,4]oxazin-7-yl)diazenyl)phenol (21):**

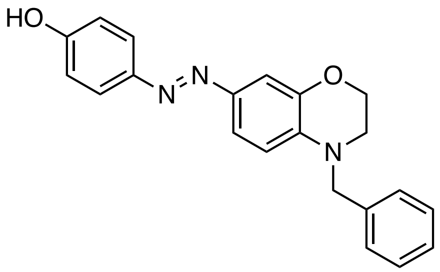


Aniline **SI-32** (150 mg, 0.62 mmol, 1.00 eq.) was dissolved in THF (1.6 ml), water (1.2 ml) and 1.0 M HCl (1.9 ml) and cooled to 0 °C. An aqueous solution of $NaNO_2$ (2.0 M, 0.38 ml, 0.75 µmol, 1.20 eq.) was added dropwise and the reaction was stirred for 30 minutes at 0 °C. Phenol (70.5 mg, 0.75 µmol, 1.20 eq.) was dissolved in aqueous NaOH (1.0 M, 1.9 ml, 1.87 mmol, 3.00 eq.) and added dropwise. The reaction was stirred for 2 h at 0 °C and upon completion poured into water (80 ml) and extracted with Ethyl acetate (3 x 80 ml, 1 x 30 ml). The combined organic layers were washed with brine, dried over $MgSO_4$, filtered and concentrated. The crude product was purified by flash column chromatography ($SiO_2$, Hex:EtOAc 10:0 to 5:5) and the title compound was obtained as a dark red solid (190 mg, 0.55 mmol, 88%).

$\mathbf{R}_f$ ($SiO_2$, Hex:EtOAc = 7:3; UV/ CAM) = 0.24.

**LCMS** (5-100% MeCN in $H_2O$ over 5 min, 0.1% FA): $t_{ret}$ = 4.5 min.

$^1$**H-NMR (400 MHz, CDCl$_3$):** $\delta$ [ppm] = 7.77 (d, $J$ = 8.6 Hz, 24H), 7.47 – 7.43 (m, 2H), 7.38 – 7.32 (m, 2H), 7.28 (d, $J$ = 8.2 Hz, 3H), 6.90 – 6.86 (m, 2H), 6.73 (d, $J$ = 9.2 Hz, 1H), 5.61 ($s_{br}$, 1H), 4.57 (s, 2H), 4.29 (t, $J$ = 4.5 Hz, 2H), 3.48 (t, $J$ = 4.5 Hz, 2H).

**$^{13}$C-NMR (101 MHz, $CDCl_3$):** $\delta$ [ppm] = 157.4, 147.5, 144.7, 143.9, 138.3, 137.2, 129.0, 127.5, 127.0, 124.3, 120.5, 115.8, 111.3, 108.7, 64.4, 54.7, 47.5.

**HRMS** (ESI$^+$, *m*/*z*): [(M+H)$^+$] for $C_{21}H_{20}N_3O_2^+$ calcd.: 346.1550, found: 346.1554.

**IR (ATR):** $\tilde{\nu}$ [cm$^{-1}$] = 1586 (m), 1515 (m), 1451 (w), 1396 (w0, 1348 (w), 1317 (s), 1274 (m), 1248 (s), 1216 (s), 1150 (s), 1123 (s), 1091 (m), 1048 (m), 973 (w), 911 (w), 877 (m), 836 (s), 802 (m), 754 (w), 712 (m).

### UV-Vis Spectra (in DMSO)

Note: methoxy acetals had the same optical properties as their hemi acetals, therefore only the representative spectrum of the hemi acetal is shown.

**3**

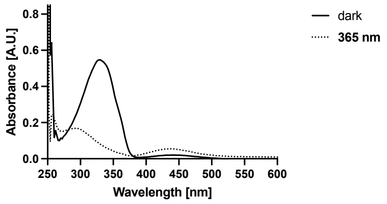


**4**

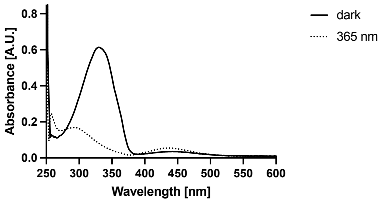


**5**

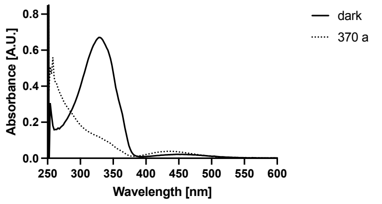


**4**

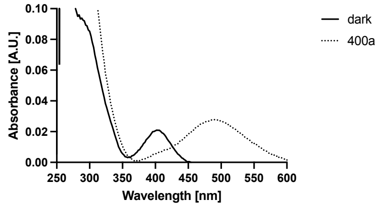


**7**

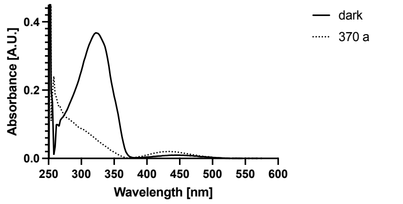


**8**

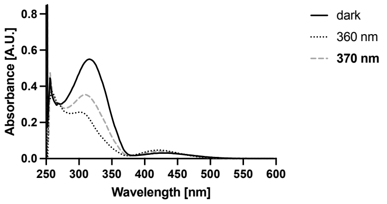


**9**

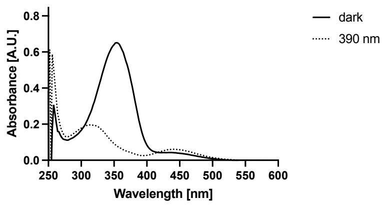


**10**

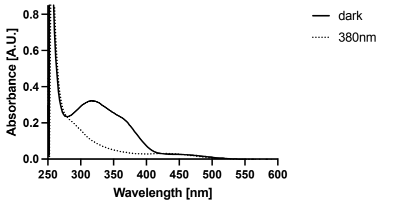


**11**

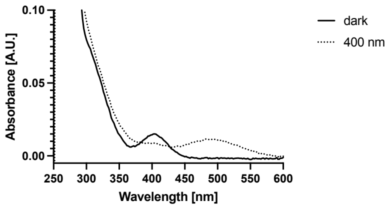


**12**

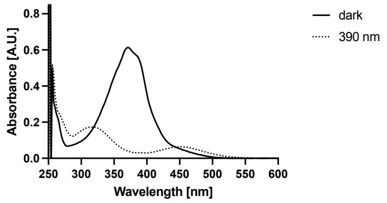


**13**

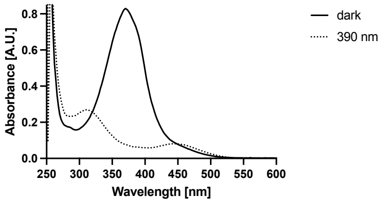


**15**

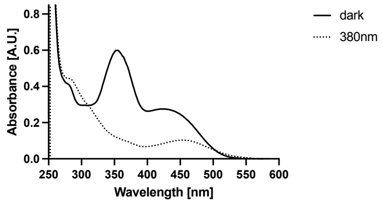

### General Methods, Biology

**Cell Culture:** HeLa (ATCC), MDA-MB-231 (ATCC) and mCherry-LifeAct MDA-MB-231 cells (gift from A. Akhmanova, Utrecht University) were cultured in antibiotic free Dulbecco's Modified Eagle Medium (DMEM) (Gibco, Thermo Fisher cat# 10566024) supplemented with 10% fetal bovine serum (FBS) (Gibco, Thermo Fischer cat# 10437036) for the first 2 passages after thawing. Thereafter they were conditioned to phenol-red free DMEM (Thermo Fisher cat# 31053036), supplemented with 10% FBS (Gibco, Thermo Fischer cat# 10437036), 1% penicillin-streptomycin-glutamine (Gibco, Thermo Fischer cat# 10378016) and a final concentration of 4 mM L-glutamine (Gibco, Thermo Fischer cat# 25030081) for 2 passages before use in assays. Cells were grown in a cell culture incubator at 37 °C in a 5% $CO_2$ atmosphere and passaged at 70-90% confluency every 2-4 days. Cells were used for up to 25 passages for cell proliferation assays and up to 15 passages for imaging studies.

**Handling of Photoswitchable Compounds:** Test compounds were dissolved in DMSO (sterile filtered) to the desired stock-concentration (e.g. 10 mM) and stored at –20 °C. In case of long half-lives, the compound stock was left in the dark at room temperature for an appropriate time to ensure full thermal relaxation of the photoswitch. Compounds were protected from light and only handled in the dark or under red-light to avoid isomerization.

**MTT Cell Proliferation Assays:** Cells (HeLa: 5000, MDA-MB-231: 4000 cells) were seeded in 96 well-plates using 90 µl phenolred-free DMEM (Thermo Fisher cat# 31053036), supplemented with 10% FBS (Gibco, Thermo Fischer cat# 10437036), 1% penicillin-streptomycin-glutamine (Gibco, Thermo Fischer cat# 10378016) and a final concentration of 4 mM L-glutamine (Gibco, Thermo Fischer cat# 25030081). After 24 h, cells were treated with compound stocks (1% DMSO, 2% MeCN as cosolvents for better solubility), which were applied as 10x concentrations in 10 µl medium. Light-dependent assays were performed as duplicates where one plate was kept in light-proof boxes, shielded from light (dark) and the second one was exposed to a specific irradiation protocol (slow: 75 ms per 15s; fast: 25 ms per 0.5 s) using a cell DISCO as described previously[17].

---

[17] Borowiak, M.; Nahaboo, W.; Reynders, M.; Nekolla, K.; Jalinot, P.; Hasserodt, J.; Rehberg, M.; Delattre, M.; Zahler, S.; Vollmar, A.; Trauner, D.; Thorn-Seshold, O. *Cell* **2015**, *162*, 403–411.

After 48 h of treatment, 3-(4,5-dimethylthiazol-2-yl)-2,5-diphenyl tetrazolium bromide (MTT; Invitrogen, Thermo Fischer Cat# M6494; 10 µl, 5 mg/ml in PBS) was added to each well and incubated for 3 h (37 °C, 5% $CO_2$). The wells were emptied and the purple formazan crystals at the bottom of the wells were dissolved in 100 µl DMSO (incubated 10 minutes, 37 °C), followed by colorimetric read-out using a FLUOstar Omega microplate reader (BMG LABTECH) (120 sec. shaking, readout at 570 nm, blank corrected).
Viability was reported as mean percentage of viable cells relative to control. ± standard deviation (SD) was reported and performed in triplicates for lead compounds. $EC_{50}$-values were determined by four-parameter curve fitting for sigmoidal dose-response with a variable slope.

### Dose response curves (MTT) for photoswitchable latrunculin derivatives and reference compounds

**3**

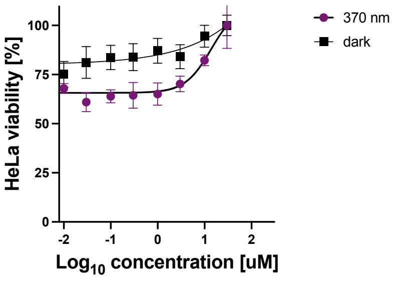


**4**

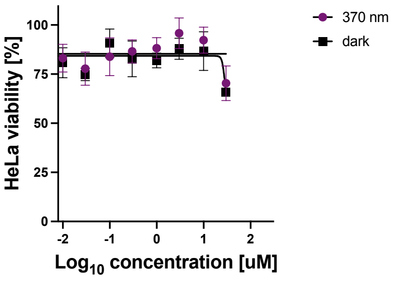


**5 OMe**

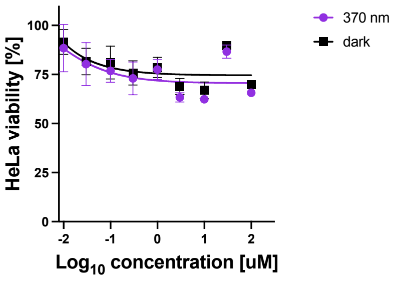


**5**

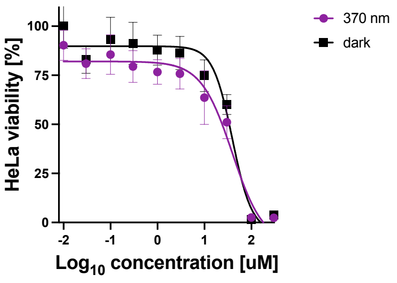


**6**

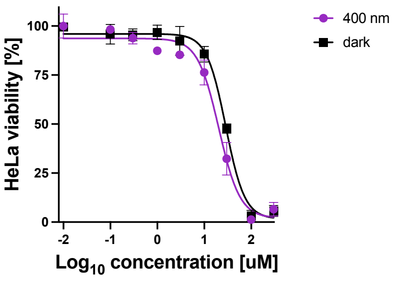


**7 OMe**

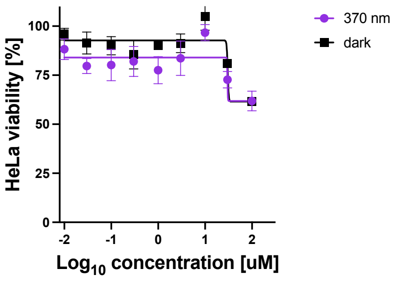

**7**

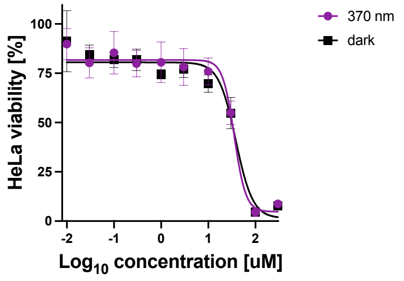


**8**

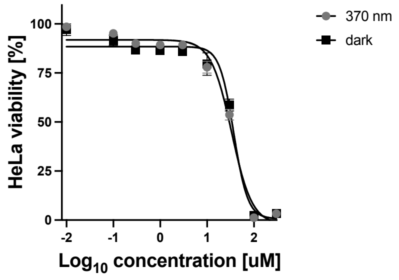


**10**

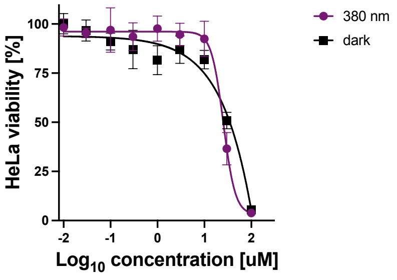


**11**

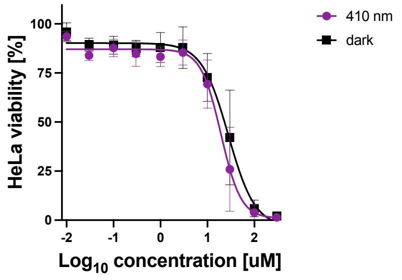


**12**

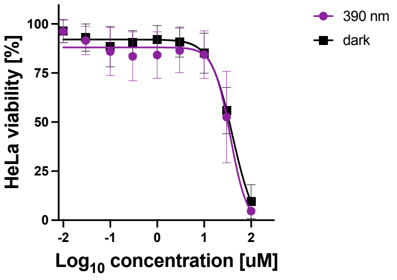


**13**

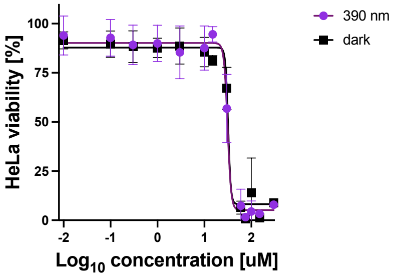


**15**

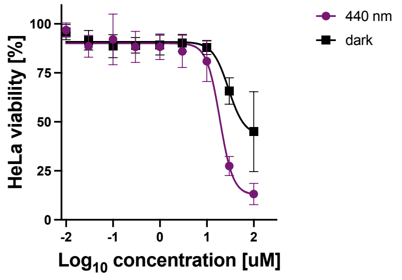


**Latrunculin A**

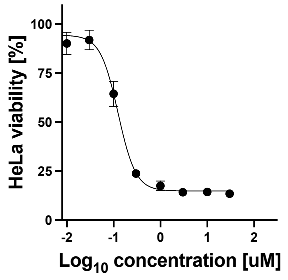


**Latrunculin B**

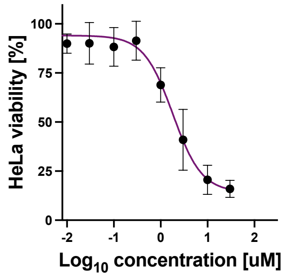


**8-*Nor*-LatB**

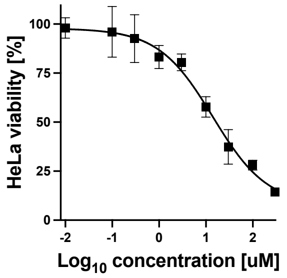

For statistical analysis and graphical representation, GraphPad Prism 9 for macOS (San Diego, CA, USA) was used. The absorbance values for untreated controls (cosolvent only) were normalized to 100%.

### Wound Healing Assays

MDA-MB-231 cells were plated in 96 well-plates at a concentration of 40k cells/well in 100 µl phenolred-free DMEM (Thermo Fisher cat# 31053036), supplemented with 10% FBS, 1% penicillin-streptomycin and a final concentration of 4 mM L-glutamine. The next day, cells were starved over night by exchanging the medium for phenolred-free DMEM (Thermo Fisher cat# 31053036), supplemented with 1% FBS (Gibco, Thermo Fischer Cat# 10437036), 1% penicillin-streptomycin-glutamine (Gibco, Thermo Fischer cat# 10378016) and a final concentration of 4 mM L-glutamine (Gibco, Thermo Fischer cat# 25030081). After a fully confluent monolayer was obtained, the monolayer was scratched using a sterilized wooden toothpick, the medium was removed, the cells washed with PBS (pH 7.4, Gibco, Thermo Fischer cat# 10010023; 3 x 100 µl, 37 °C) and 90 µl full growth medium was added. The obtained scratches were imaged using a Leica DMI6000B inverted fluorescent microscope with a Tokai Hit stage-top incubator at 37 °C at 2.5 x magnification, DIC filter (timepoint t=0), before the compounds were added (10x concentration, 10 µl). Experiments were run in sets of two duplicates (dark/ light protocol) using the cell DISCO system with a 24-LED array as previously reported. Quantification of scratch area was performed using a custom macro script for ImageJ (National Institutes of Health, USA). The final selection area was adjusted to fit the wound in every image of the image-stack. Relative wound closure per well was referenced to timepoint $t_0$ and further analyzed using GraphPad Prism 9 for macOS (San Diego, CA, USA). Overall, three independent experiments were performed. Fig, X shows data points that were combined from two independent experiments under optimized conditions by internal normalization to the wound closure (dark) after 24 h. Statistical analysis was performed using GraphPad Prism 9 for macOS (San Diego, CA, USA). Error bars are reported as ± standard deviation (S.D.) from 2 independent experiments (n = 6).
After 24 h, cell viability was assessed using PrestoBlue viability assay (Thermo Fischer cat# A13261; 10 µl/ well), incubated for 30 minutes and analyzed by fluorescence readout $\lambda_{Ex}$ 544 nm/ $\lambda_{Em}$ 590 nm using a FLUOstar Omega microplate reader (BMG LABTECH, Ortenberg, Germany).

**18 M OLat** conc: significant but p value smaller because of small set of data points (1 biol. replicate).

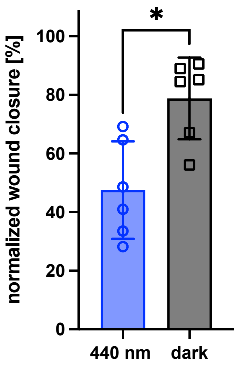


**Macro:**

```
run("Images to Stack", "name=Stack title=[] use");
run("Find Edges", "stack");
setOption("BlackBackground", true);
run("Make Binary", "method=Default background=Default calculate black");
run("Dilate", "stack");
run("Fill Holes", "stack");
run("Dilate", "stack");
run("Fill Holes", "stack");
run("Invert", "stack");
makeRectangle(657, 753, 768, 576);
```

### Live-Cell Imaging with Local Illumination

MDA-MB-231 cells that stably express mCherry LifeAct were seeded using phenol-red free DMEM (Thermo Fisher cat# 31053036), supplemented with 10% FBS, 1% penicillin- streptomycin and a final concentration of 4 mM L-glutamine at a density of 40k cells/ 2 ml in a ø35 mm imaging dish (previously coated with poly-L-lysine; see procedure for cover slips) and left to adhere for 16-24 h. Next, the medium was removed and new medium containing **OptoLat** was added. The cells were left to incubate for 16-24 h in the dark. When transporting cells, it was ensured that cells were kept in a warm environment and in the dark.

The environmental chamber of a Zeiss LSM880 Airyscan microscope system (ZEISS, Oberkochen, Germany) was preheated to 37 °C prior to imaging. Cells were transported as quickly as possible in a tightly sealed styrofoam box with 2 x 500 ml water bottles that had been equilibrated to 37 °C and cells immediately transferred to the environmental chamber of the microscope. The imaging dish was mounted on the stage under minimal ambient light and using red head lamps to allow for better vision. Cells were focused using a red filter foil in the light path oft he microscope to shield blue light from reaching the sample during the light exposure that was required to focus.

Images were acquired using a 63x objective. 440 nm light pulses and ROI were specified using the FRAP tool and different acquisition blocks were programmed in row using the experiment designer in ZEN (ZEISS, Oberkochen, Germany).

Image analysis and processing was performed using ImageJ and Fiji – ImageJ (National Institutes of Health, USA)[18].

### Experiments with Oligodendrocytes

Oligodendrocyte precursor cells were purified from enzymatically dissociated, mixed-sex P5-P7 Sprague-Dawley (Charles Rivers) rat brains by immunopanning and grown in serum-free defined medium at 10%$CO_2$ 37˚C, as described previously[19]. PDGF (10 ng/ml, PeproTech) and NT-3 (1 ng/ml, PeproTech) were added to the media to proliferate OPCs for 3-5 days. Before confluency was reached, OPCs were trypsinized (0.0025% trypsin-EDTA, Gibco), re-plated at 4K cells/12mm PDL-coated glass coverslip and incubated with thyroid hormone (triiodothyronine, T3; 40 ng/ml; Sigma) to induce differentiation. After 24-30 hours in culture, optolatrunculin or DMSO was added to appropriate wells and incubated overnight in the dark. Unless otherwise stated, coverslips were then moved to a climate-controlled incubator with 1Hz flashing 430nm LED for 6 hours. Latrunculin-A treated cells were treated for 4 hours with 100nM LatA (Invitrogen). Coverslips were fixed for 15 min in 4% paraformaldehyde (PFA) and rinsed 3 times in PBS. Cells were permeabilized in 0.1% Triton X-100 in PBS for 3 min and rinsed 3 times in PBS. Coverslips were incubated in PBS with 50 nM Alexa Fluor 488-phalloidin (Invitrogen) to visualize actin filaments, and 5ug/mL HCS CellMask™ Blue Stain (invitrogen) to reveal cellular morphology. Cells were rinsed three times in PBS and mounted with Fluoromount-G (Southern Biotech). Cells were visualized by epifluorescence using an inverted Zeiss Axio Observer with a 40x oil objective and Zen Blue Software. Slides were blinded and individual cells were chosen based on the CellMask channel. Identical illumination and acquisition conditions were used for each experiment.

Regions of interest (ROIs) were defined by thresholding on phalloidin in ImageJ by blinded investigators. Mean gray value (average intensity) was measured for each cell. To normalize values across different biological replicates, the value for each cell was divided by the average value for DMSO-treated

[18] Schindelin, J.; Arganda-Carreras, I.; Frise, E.; Kaynig, V.; Longair, M.; Pietzsch, T.; Preibisch, S.; Rueden, C.; Saalfeld, S.; Schmid, B.; Tinevez, J.-Y.; White, D. J.; Hartenstein, V.; Eliceiri, K.; Tomancak, P.; Cardona, A. *Nat Methods* **2012**, *9* (7), 676–682.
[19] Dugas, J.C., and Emery, B. (2013). Purification of oligodendrocyte precursor cells from rat cortices by immunopanning. Cold Spring Harb. Protoc. 2013, 745–758.

cells in the same technical replicate and reported as one individual cell in the superplot. Data from 2-3 technical replicate coverslips from a given experimental day were averaged for the value reported as one biological replicate. Data were analyzed and plotted using Prism (GraphPad Software). Unless otherwise stated, error bars are SD, and p values were calculated using one-way ANOVA followed by Dunnett's multiple comparison test.

**Basal optolatrunculin activity in oligodendrocytes**

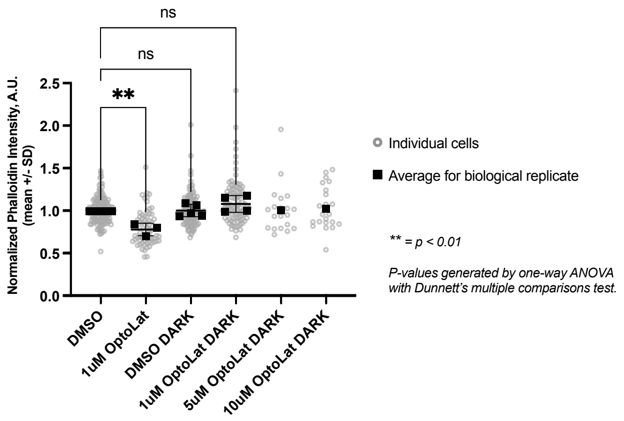


### Experiments with Budding Yeast Cells

**Yeast Cell growth conditions.** This study was conducted using the yeast BY4741 strain (*MATa his3Δ1 leu2Δ0 met15Δ0 ura3Δ0*) (Open Biosystems, Huntsville, AL). The cultures used for **OptoLat** experiments were grown to mid-log phase ($OD_{600}$ 0.1-0.3) in glucose-based synthetic complete (SC) medium [0.67% (w/v) yeast nitrogen base without amino acids and with ammonium sulfate, amino acids, 2% (w/v) glucose] at 30°C.

**Visualization of F-actin cytoskeleton.** Mid-log phase yeast cells ($OD_{600}$ 0.5) were treated with 100 μM **OptoLat** dissolved in DMSO or the same amount of DMSO (HybriMax, Sigma Aldrich, St. Louis, MO), and exposed to pulsed blue light (430 nm) or kept in the dark for 0, 5, and 20 minutes in 12-well plates (Costar, Corning, NY). Cells were then fixed in 3.7% paraformaldehyde (Electron Microscope Sciences, Hatfield, PA) at 30°C for 30 min. Fixed cells were washed three times with 500 μL of 1x PBS (137 mM sodium chloride, 2.7 mM potassium chloride, 10.14 mM sodium phosphate dibasic, and 1.77 mM potassium phosphate monobasic, and adjusted to pH

8), and then washed with 500 μL 1x PBT (1x PBS with 1% w/v BSA, 0.1 % v/v Triton X-100, 0.1% w/v sodium azide), and actin was stained with 2.5 μM AlexaFluor488 (AF488)-conjugated phalloidin (Thermo Fisher Scientific, Waltham, MA) for 25 min at room temperature (RT) in the dark. Stained cells were then washed three times with 100 μL 1x PBS and resuspended in 5 μL of SlowFade Diamond anti-fade mounting medium (Thermo Fisher Scientific, Waltham, MA). 1.8 μL of phalloidin-stained cells were mounted on a glass side and covered with a #1.5 coverslip, and the edges were sealed with nail polish.

**Microscopy and image acquisition**

Fluorescence microscopy was performed using a Zeiss Axioskop 2 Plus upright fluorescence microscope equipped with a 100x/ 1.4 Numerical Aperture (NA) Zeiss Plan-Apochromat objective lens (Carl Zeiss Inc., White Plains, NY), a Light-Emitting Diode (LED) module (CoolLED pE-400, Andover, UK) and an Orca ER cooled charge-coupled device (CCD) camera (Hamamatsu Photonics, Hamamatsu City, Japan). The imaging system was controlled using NIS Elements 4.60 Lambda software (Nikon, Melville, NY). AF488-conjugated phalloidin was visualized using 470 nm illumination, and emitted fluorescence was filtered using a dual eGFP/mCherry cube (#59222, Chroma, Bellows Falls, VT). Z-stacks of AF488-phalloidin stained F-actin structures were captured at 0.3 μm steps using 1x1 binning, 116 digital gain, and 200-300 ms exposure time. Images were deconvolved using Volocity 5.5 (Quorum Technologies Ltd, Puslinch, ON, Canada) with a constrained iterative restoration algorithm (507 nm emission wavelength, 100% confidence limit, and 60 iterations). Images shown were contrast-enhanced using the same parameters.

**Quantification and Statistical Analysis**

F-actin content in actin cables was quantified by measuring the fluorescence intensity of AF488 phalloidin-stained actin cables in a region of interest in mother yeast cells using Volocity 5.5 (Quorum Technologies Ltd, Puslinch, ON, Canada). Statistical analysis and generation of Super plots from the data were carried out using GraphPad Prism 8 (GraphPad Software, San Diego, CA). Two-tailed statistical analysis was applied to samples analyzed. For multiple group comparisons and control studies, *p* values were determined using the 2-way ANOVA Kruskal-Wallis test with Tukey's multiple comparisons. For all tests, *p* values are classified as follows: *$p$

< 0.05; ***p* < 0.01; *** *p* < 0.001; *****p* < 0.0001, unless exact *p* values are noted in the figure legends.

### Experiments with Microglia

#### Mouse hippocampal slice culture

Mice were housed and bred at the University Medical Center Hamburg-Eppendorf. All procedures were performed in compliance with German law and the guidelines of Directive 2010/63/EU. The study was approved by the local authorities (Amt für Verbraucherschutz, Lebensmittelsicherheit und Veterinärwesen, Hamburg; permission # 42/17). Mice carrying a tamoxifen-inducible Cre-recombinase in microglia and the floxed fluorescent marker tdTomato (B6.129 - Cx3cr1$^{tm2.1(cre/ERT2)Jung}$ Gt(ROSA26)Sortm$^{9(CAG\text{-}tdTomato)Hze}$) (JAX 020940; JAX 007909) were used to prepare hippocampal slice cultures at postnatal day 4–7 as described[20]. Briefly, mice were anesthetized with 80% $CO_2$ 20% $O_2$ and decapitated. Hippocampi were dissected in cold dissection medium containing (in mM): 248 sucrose, 26 $NaHCO_3$, 10 glucose, 4 KCl, 5 $MgCl_2$, 1 $CaCl_2$, 2 kynurenic acid and 0.001% phenol red. pH was 7.4, osmolarity 310-320 mOsm/kg, saturated with 95% $O_2$, 5% $CO_2$. Tissue was cut into 410 $\mu$M thick sections on a tissue chopper and cultured at the medium/air interface on membranes (Millipore PICMORG50) at 37° C in 5% $CO_2$. For the first 24 h of incubation, 1 $\mu$M (Z)-4-hydroxytamoxifen was added to the slice culture medium to activate Cre. Slice culture medium was partially exchanged (60-70%) twice per week and contained (for 500 ml): 394 ml Minimal Essential Medium, 100 ml heat inactivated donor horse serum, 1 mM L-glutamine, 0.01 mg $ml^{-1}$ insulin, 1.45 ml 5M NaCl, 2 mM $MgSO_4$, 1.44 mM $CaCl_2$, 0.00125% ascorbic acid, 13 mM D-glucose. Slice cultures were used for experiments between 12 and 28 days *in vitro*. Before light stimulation, cultures were incubated in slice culture medium with 30 uM NAV-872 in 0.5% DMSO for 16 h or with 0.5% DMSO only (vehicle control).

#### Two photon imaging

Organotypic hippocampal slice cultures were placed in the recording chamber of a custom-built two-photon laser scanning microscope based on an Olympus BX51WI microscope with HC Fluotar L 25x 0.95 NA (Leica) objective, controlled by ScanImage 2017b[21] (Pologruto, Sabatini, and Svoboda 2003). A Ti:Sapphire laser (Chameleon Ultra, Coherent) controlled by an electro-optic modulator (350-80, Conoptics) was used

---

[20] Gee, Christine E., Iris Ohmert, J. Simon Wiegert, and Thomas G. Oertner. 2017. “Preparation of Slice Cultures from Rodent Hippocampus.” *Cold Spring Harbor Protocols* 2017 (2). https://doi.org/10.1101/pdb.prot094888.

[21] Pologruto, Thomas A., Bernardo L. Sabatini, and Karel Svoboda. 2003. “ScanImage: Flexible Software for Operating Laser Scanning Microscopes.” *Biomedical Engineering Online* 2 (1): 13.

to excite tdTomato in microglia at 980 nm. During imaging, slice cultures were continuously perfused with a HEPES-buffered solution (in mM): 135 NaCl, 2.5 KCl, 10 Na-HEPES, 12.5 D-glucose, 1.25 $NaH_2PO_4$, 2 $CaCl_2$, 1 $MgCl_2$ (pH 7.4, 308 mOsm) at 31-33°C. Image stacks were acquired at 2 min intervals for 50 min. An LED light engine (Lumencor) was used for epifluorescence and activation of OptoLat (475 nm). During blue light pulses (250 ms pulses at 1 Hz, pulse intensity 22 mW/mm$^2$), PMTs were protected by a computer-controlled shutter system (Vincent Associates).

#### Analysis of microglia ramification and surveillance

We combined several analysis steps into a semi-automated workflow[22]. Image data were background-subtracted (rolling ball radius = 30 pixels) and median filtered (radius = 1 pixel) with ImageJ. In order to correct for small positional shifts that occurred during acquisition, we registered all images (Efficient subpixel image registration by cross-correlation, version 1.1, MATLAB Central File Exchange). Images were then binarized using Otsu's method (threshold set to 50% calculated value). Based on a maximum projection of both images and binary images, a mask was drawn to define the region considered for motility analysis. Within the masked region we counted the number of pixels that remained stable, that were lost, and that were gained throughout the time series. From the covered areas and perimeters, we calculated ramification and surveillance indices

[22] Laprell, Laura, Christian Schulze, Marie-Luise Brehme, and Thomas G. Oertner. 2021. "The Role of Microglia Membrane Potential in Chemotaxis." *Journal of Neuroinflammation* 18 (1): 21.
